# Background selection does not mimic the patterns of genetic diversity produced by selective sweeps

**DOI:** 10.1101/2019.12.13.876136

**Authors:** Daniel R. Schrider

## Abstract

It is increasingly evident that natural selection plays a prominent role in shaping patterns of diversity across the genome. The most commonly studied modes of natural selection are positive selection and negative selection, which refer to directional selection for and against derived mutations, respectively. Positive selection can result in hitchhiking events, in which a beneficial allele rapidly replaces all others in the population, creating a valley of diversity around the selected site along with characteristic skews in allele frequencies and linkage disequilibrium (LD) among linked neutral polymorphisms. Similarly, negative selection reduces variation not only at selected sites but also at linked sites—a phenomenon called background selection (BGS). Thus, discriminating between these two forces may be difficult, and one might expect efforts to detect hitchhiking to produce an excess of false positives in regions affected by BGS. Here, we examine the similarity between BGS and hitchhiking models via simulation. First, we show that BGS may somewhat resemble hitchhiking in simplistic scenarios in which a region constrained by negative selection is flanked by large stretches of unconstrained sites, echoing previous results. However, this scenario does not mirror the actual spatial arrangement of selected sites across the genome. By performing forward simulations under more realistic scenarios of BGS, modeling the locations of protein-coding and conserved noncoding DNA in real genomes, we show that the spatial patterns of variation produced by BGS rarely mimic those of hitchhiking events. Indeed, BGS is not substantially more likely than neutrality to produce false signatures of hitchhiking. This holds for simulations modeled after both humans and *Drosophila*, and for several different demographic histories. These results demonstrate that appropriately designed scans for hitchhiking need not consider background selection’s impact on false positive rates. However, we do find evidence that BGS increases the false negative rate for hitchhiking—an observation that demands further investigation.

## 2 Introduction

The impact of natural selection on genetic diversity within and between species has been debated for decades (Kimura 1968, 1983; Gillespie 1984, 1991; Kern and Hahn 2018; Jensen *et al*. 2019). Perhaps the strongest evidence that selection influences the amount of diversity at linked neutral alleles comes from the correlation between diversity levels and recombination rates across the genome (Begun and Aquadro 1992; Smukowski and Noor 2011; McGaugh *et al*. 2012; Corbett-Detig *et al*. 2015). This observation is consistent with genetic hitchhiking, in which a beneficial mutation rapidly increases in population frequency and carries its genetic background along with it. These events, which are also referred to as selective sweeps, result in the complement of genetic diversity in the vicinity of the selected site to be largely replaced by descendants of the chromosome(s) that acquired the adaptive mutation. The width of the resulting valley of diversity will depend in part on the recombination rate, as crossover events will allow linked variation to “escape” by shuffling alleles onto and off of the set of sweeping chromosomes (Maynard Smith and Haigh 1974; Kaplan *et al*. 1989). The correlation between recombination rate and diversity can also be explained by background selection (BGS), wherein neutral alleles linked to a deleterious mutation are purged via negative (or purifying) selection unless they can escape via recombination (Charlesworth *et al*. 1993). Whether primarily due to hitchhiking, BGS, or—more likely—a combination of the two (Elyashiv *et al*. 2016; Booker and Keightley 2018), there is growing evidence that natural selection has a profound impact on the amount and patterns of diversity across the genome in a variety of species (Begun *et al*. 2007; Lohmueller *et al*. 2011; Langley *et al*. 2012; Corbett-Detig *et al*. 2015; Booker and Keightley 2018). Indeed, it appears that natural selection is in part responsible for the limited range of levels of genetic diversity observed across species (Lewontin 1974; Leffler *et al*. 2013; Corbett-Detig *et al*. 2015), though other forces are likely at play as well (Coop 2016).

There is ample reason to suspect that BGS may substantially reduce the amount of polymorphism genome-wide. For example, approximations have been derived for the expected percent reduction in diversity at a given site, termed *B*, due to negative selection acting on linked sites (Hudson and Kaplan 1994, 1995; Nordborg *et al*. 1996). The extent to which BGS effects diversity can thus be predicted from a genome annotation and estimated distribution of fitness effects (DFE; e.g. McVicker *et al*. 2009). While there may be some uncertainty over the true DFE, such *B*-maps predict that BGS has a sizeable impact, removing an estimated ∼20% and ∼45% of diversity on the autosomes in humans and *Drosophila*, respectively (McVicker *et al*. 2009; Comeron 2014).

More controversial is the potential role of positive selection in shaping the landscape of diversity across the genome (Stephan 2010). A number of approaches exist for detecting positive selection in population genomic data. For example, variants of the McDonald-Kreitman test, which searches for an excess of nonsynonymous divergence between species, have found that in many organisms a large fraction of amino acid substitutions are beneficial (Smith and Eyre-Walker 2002; Charlesworth and Eyre-Walker 2006; Galtier 2016; Enard *et al*. 2016). Efforts have also been made to fit genome-wide parameters of recurrent hitchhiking models to population genetic data in *Drosophila*, in some cases suggesting appreciable rates of hitchhiking events (Andolfatto 2007; Jensen *et al*. 2007; Li and Stephan 2006). An alternative approach to assess the frequency of adaptive substitutions is to directly search for recent selective sweeps. For this reason, and because identifying hitchhiking may provide clues about recent adaptations and selective pressures, a large number of methods for locating selective sweeps in the genome have been devised (Hudson *et al*. 1994; Fay and Wu 2000; Kim and Stephan 2002; Sabeti *et al*. 2002; Kim and Nielsen 2004; Nielsen *et al*. 2005; Voight *et al*. 2006; Lin *et al*. 2011; Ronen *et al*. 2013; Ferrer-Admetlla *et al*. 2014; Pybus *et al*. 2015; Schrider and Kern 2016; Mughal and DeGiorgio 2018). Nevertheless, detecting signatures of hitchhiking remains a major challenge. For example, it is well known that demographic events such as population bottlenecks can mirror selective sweeps (*Simonsen et al*. 1995; Jensen *et al*. 2005; Nielsen *et al*. 2005) and the feasibility of detecting hitchhiking in the presence of non-equilibrium demography remains hotly debated (Harris *et al*. 2018; Schrider and Kern 2018). It has also been suggested that BGS could be mistaken for selective sweeps (e.g. Comeron *et al*. 2012; DeGiorgio *et al*. 2016), and it is this possibility that we investigate here.

Intuitively one may expect BGS to resemble hitchhiking because both forces can create localized reductions of polymorphism. Moreover, BGS is a very flexible model in part because of the astronomical number of possible arrangements of selected sites across a chromosome—it is straightforward to construct a BGS scenario that somewhat mirrors the valley of diversity caused by hitchhiking by placing a large cluster of selected sites in a region flanked by vast stretches of unselected sites (e.g. Mughal and DeGiorgio 2018). At first blush this may suggest that distinguishing between hitchhiking and BGS should be extremely challenging (reviewed in Stephan 2010). However, in practice we often know (with some degree of uncertainty) where selected sites reside in genomes with high-quality annotations of genes and conserved non-coding elements (CNEs); such information is used to create the *B*-maps alluded to above. Thus, rather than focusing on the most pessimistic scenario in which BGS could mirror selective sweeps, it is possible to ask how often BGS would be expected to actually mirror hitchhiking in real genomes. In addition, hitchhiking events affect diversity in ways other than just removing polymorphism: they can also dramatically skew the site frequency spectrum towards low- and high-frequency derived alleles (Braverman *et al*. 1995; Fay and Wu 2000) and increase linkage disequilibrium (LD) on either flank of the selected site while reducing LD between polymorphisms on opposite sides of the sweep site (Kim and Nielsen 2004). While BGS may also influence these aspects of polymorphism, such effects may be subtle (Charlesworth *et al*. 1995; Tachida 2000; Zeng 2013). Thus, it is unclear whether BGS will resemble hitchhiking when additional features of genetic variation are examined.

With this in mind, here we examine the separability of BGS and hitchhiking models via simulations of large genomic regions summarized by a number of statistics capturing the amount of nucleotide and haployptic diversity, the shape of the site frequency spectrum, and patterns of linkage disequilibrium. Our approach is to simulate BGS scenarios designed to match annotated genomes with respect to their locations of selected sites and estimated DFEs, and to compare the resulting patterns of diversity to those expected under selective sweeps. In addition to qualitative comparisons of these models using various summaries of diversity, we also use a classification approach to ask how often BGS simulations are mistaken for hitchhiking events, as this is informative about how frequently realizations of these two models will resemble one another. We first examine simulations modeled after the human genome, including both equilibrium and non-equilibrium demographic histories. We then consider the *Drosophila melanogaster* genome, which has both a much larger density of selected sites and a different estimated DFE. We then conclude with a discussion of the implications of our results for efforts to detect positive selection in the face of potentially widespread BGS.

## 3 Methods

### 3.1 Annotation Data

We downloaded annotation data from the UCSC Table Browser (Karolchik *et al*. 2004) for both the human (Lander *et al*. 2001) and *D. melanogaster* (Adams *et al*. 2000) genome assemblies (GRCh37/hg19 and Release 5/dm3 coordinate spaces, respectively; all data accessed on Dec 21, 2018). These data included refSeq protein-coding gene coordinates for both genomes and phastCons elements (Siepel *et al*. 2005) conserved across vertebrates and insects, respectively, as well as the locations of gaps within both genome assemblies. The human genetic map from Kong et al. (2010) was also obtained from this resource. In addition to data from the UCSC Table Browser, we used the *D. melanogaster* genetic map from Comeron *et al*. 2012.

### 3.2 Overview of simulation strategies

We used four different simulation strategies to assess the similarity between BGS and hitchhiking models. These include: 1) forward simulations of various scenarios of BGS in moderately sized chromosomal windows (e.g. 1 cM in humans); 2) forward simulations of BGS in larger chromosomal regions, 3) coalescent simulations of selective sweeps used both to qualitatively compare to BGS and to train a classifier to more formally quantify the similarity of BGS and hitchhiking (by asking how often simulations of BGS are misclassified as selective sweeps); and 4) forward simulations containing both BGS and selective sweeps, used for assessing the extent to which the signatures of hitchhiking events are weakened in the presence of BGS.

### 3.3 Forward simulations of background selection

We used `fwdpy11` version 0.1.4 (Thornton 2014) to perform forward simulations of 1.1 Mb regions modeled after human populations and 110 kb regions modeled after *D. melanogaster*. In all simulations with BGS, 75% of mutations within selected regions (either exons or CNEs) were deleterious (i.e. the selection coefficient, *s*, was drawn from the appropriate DFE), while the remaining 25% were selectively neutral. The selection coefficients for deleterious mutations were gamma-distributed: the DFE for deleterious mutations in humans had a mean of −0.030 and shape parameter of 0.206 (as estimated by Boyko *et al*. 2008) and for *Drosophila* the mean and shape were −0.000133 and 0.35 respectively (Huber *et al*. 2017). We simulated populations under four different scenarios (Figure 1): 1) No selection 2) a scenario we refer to as Central BGS, wherein the central ∼5% of the simulated region is a coding sequence; 3) a scenario where each simulated replicate is modeled after a randomly selected genomic region as described below—we refer to this scenario as Real BGS because it should more accurately model BGS in real genomes than does scenario 2; 4) and a scenario identical to 3 but where the selection coefficients in CNEs are 10-fold lower than in coding regions, though the DFEs have the same shape (Real BGS– Weak CNE). Each of our simulated scenarios contained a single fixed dominance coefficient, *h*, for all deleterious mutations. We simulated replicates of each Real BGS scenario (scenarios 3 and 4 above) with dominance values of 0, 0.25, 0.5, and 1.0. The fitness values of homozygous wild-type, heterozygous, and homozygous mutant individuals were 1, 1-*hs*, and 1-*s*, respectively. Unless otherwise noted, we report results from simulations with dominance of 0.25 because deleterious mutations may often be partially recessive (GarcÍa-Dorado and Caballero 2000; Peters *et al*. 2003; Agrawal and Whitlock 2011), although our results do not appear to change qualitatively with different dominance values as discussed in the Results. All forward simulations began with a burn-in period of 10*N* generations where *N* is the ancestral population size. Although this may not have been a sufficient burn-in period to ensure that all lineages coalesce normally in the ancestral population, the strong concordance between the mean values of summary statistics calculated from our neutral forward and coalescent simulations (e.g. Figure 3) implies that the burn-in duration did not substantially impact our results. At the end of each simulation we randomly sampled 100 chromosomes from the population.

**Figure 1:**
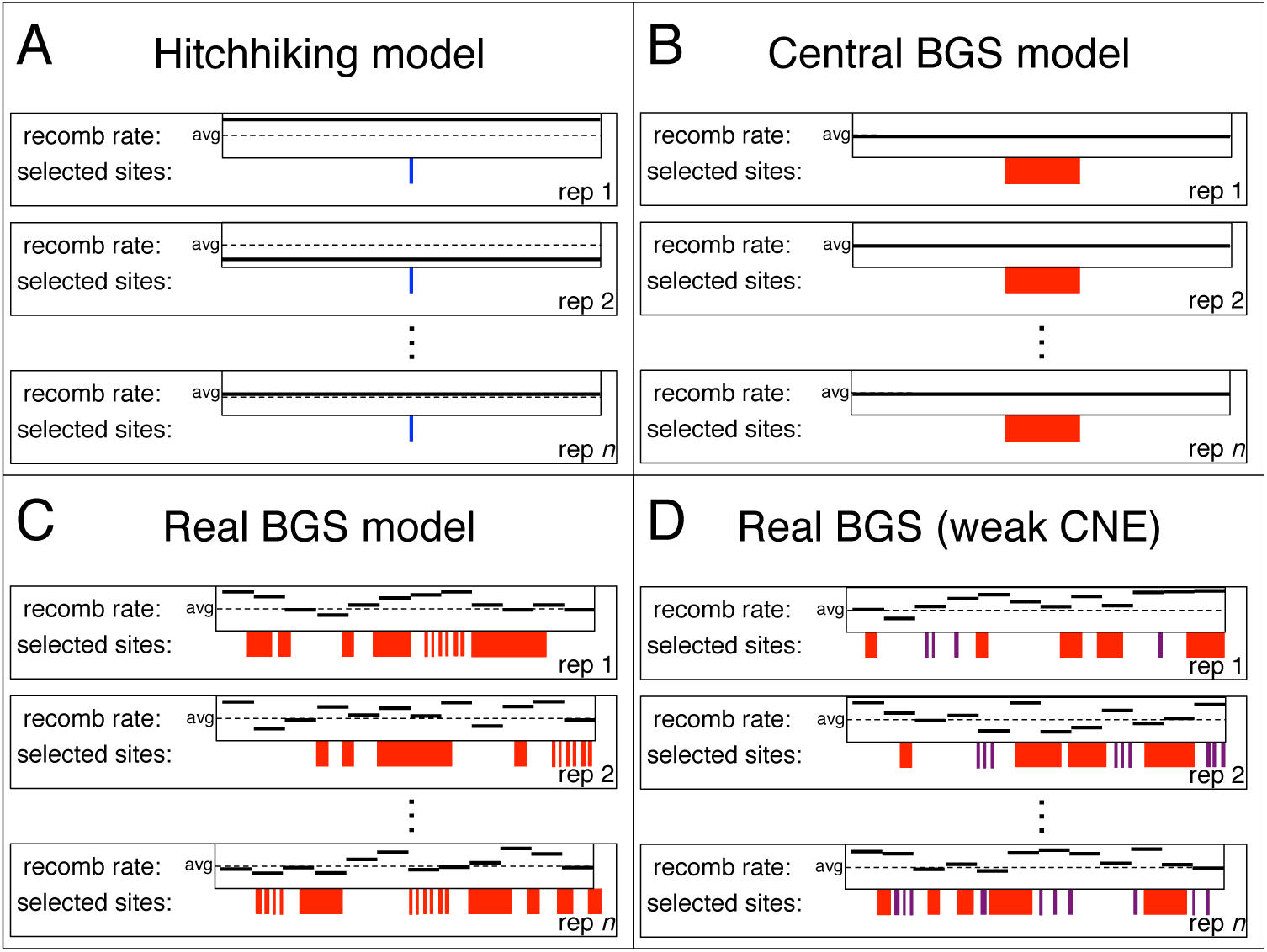
Overview of selection models examined in this study. (A) Hitchhiking models (either hard or soft sweeps) in which there is a recent sweep in the center of the simulated region (blue site), and the recombination rate is flat but varies across simulated replicates. (B) The Central BGS model in which mutations in the central ∼10% of the region can be deleterious but all other mutations are neutral. (C) The Real BGS model in which for each replicate both the recombination map and the locations of sites experiencing negative selection (red regions) are drawn from a randomly selected region in the genome, but all selected sites have the same DFE. (D) The Real BGS model with weak selection on CNEs, which is the same as the Real BGS model but mutations at CNEs (purple) have 10-fold lower *s* than those within exons (red) on average.

**Figure 2:**
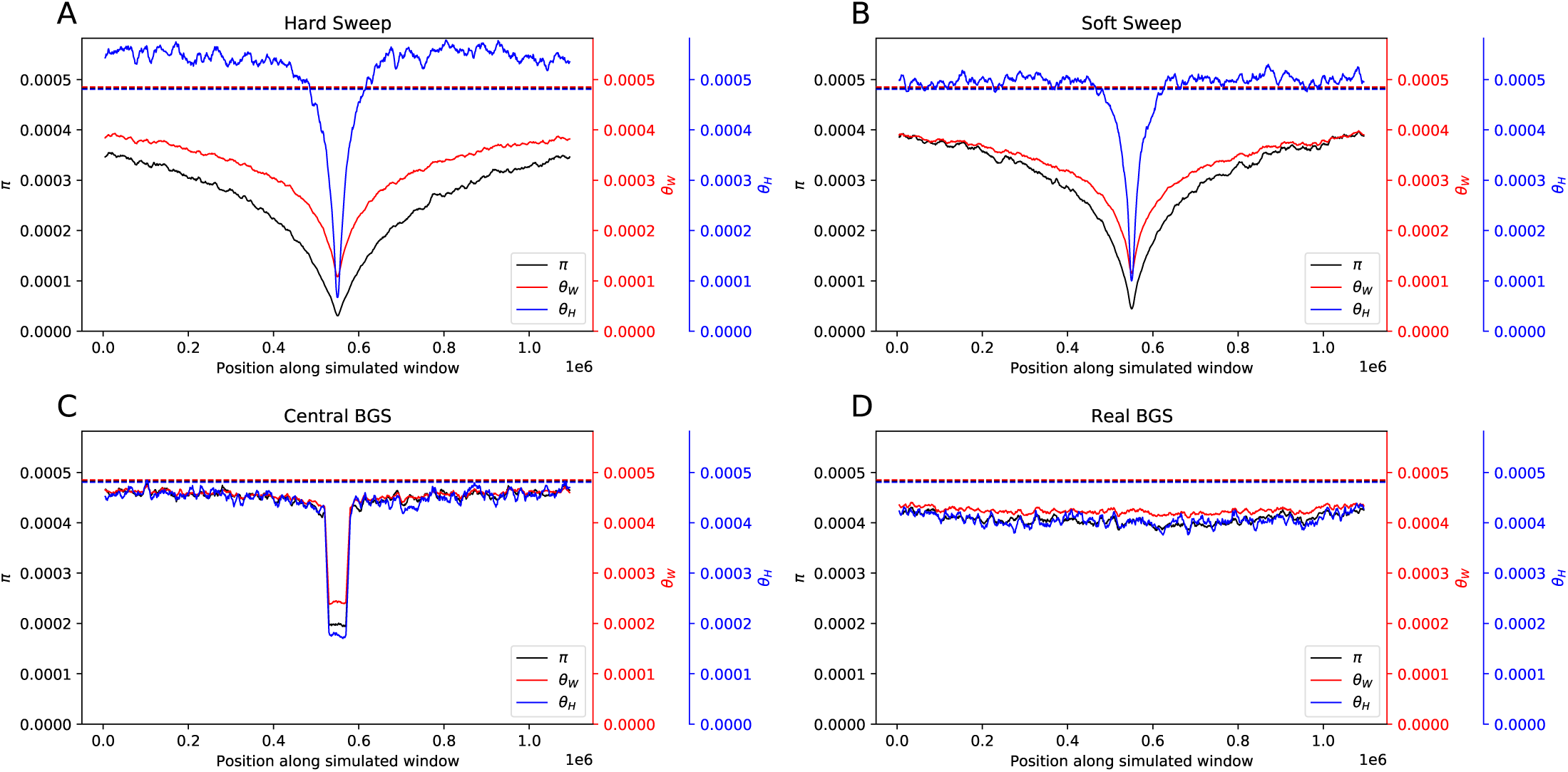
Patterns of diversity produced by different models of linked selection under constant population size. (A) Values of three estimators of *θ* in regions experiencing a recent hard sweep at the center of the window. (B) Values of these estimators in regions with a recent soft sweep at the center. (C) Values in regions where purifying selection is acting on a central 50 kb element. (D) Values in regions with purifying selection acting on elements distributed across the chromosome in a manner that mirrors the locations of exons and CNEs in randomly selected 1.1 Mb regions in the human genome. Each panel shows windowed statistics calculated as described in the Methods and averaged across 1000 simulated replicates, and the mean value of each statistic across the neutral forward simulations is also shown as a dashed line.

**Figure 3:**
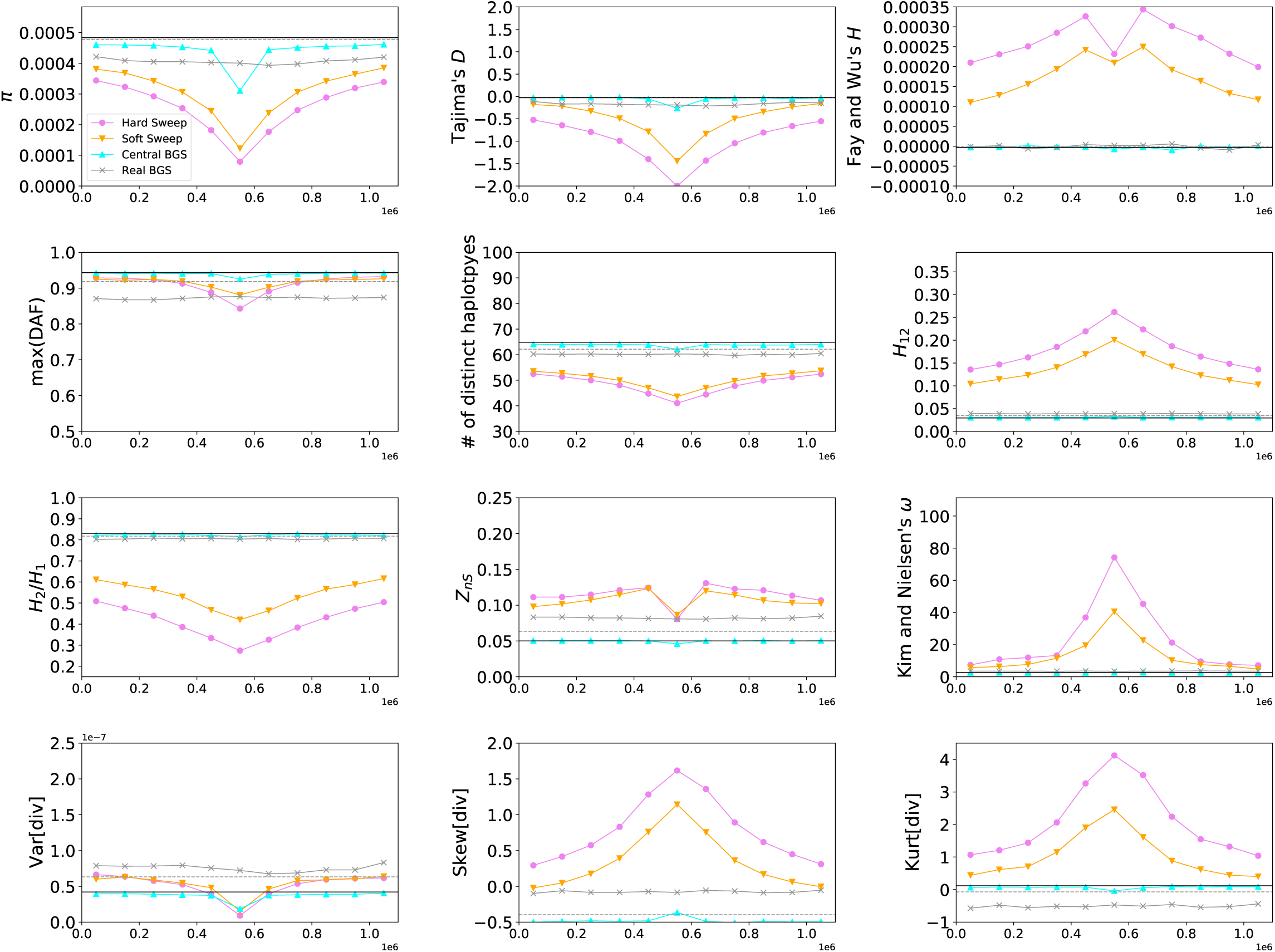
Values of 15 statistics calculated in each of the simulation conditions shown in Figure 2. Here, the mean value of each statistic (calculated from 1000 simulation replicates) is shown in the 11 adjacent 100 kb windows within the 1.1 Mb chromosomal region. In each panel the neutral expectation of the statistic obtained from forward simulations are shown as a horizontal black line, while the neutral expectation from coalescent simulations is shown as a dashed gray line—although for most statistics these two expectations were essentially identical, for some there was a slight difference perhaps reflecting the subtle differences in the models used by the two simulators, and/or incomplete burn-in.

For each replicate of our Real BGS simulations, we modeled a random genomic window of the appropriate size for the two species. This was done by first selecting the endpoint of the window, which we constrained to be a multiple of 100 kb for humans and 10 kb for *Drosophila*. Window locations were drawn with replacement, and all possible locations across the genome had equal probability of being selected, though windows with at least 75% of positions in assembly gaps were disallowed. The simulation replicate was then modeled after the selected region by taking the locations of annotated exons and phastCons elements within the region and allowing deleterious mutations to occur at these sites only, i.e. all mutations outside of these elements were neutral. Our neutral simulations followed the same procedure as the Real BGS simulations for randomly selecting a region to model but used only the region’s recombination landscape.

For our human simulations we used two different demographic histories: a constant-sized population with *N*_*e*_ = 10, 000, and the European population size history estimated by Tennessen et al. (2012); the latter model contains two successive population contractions followed by a period of exponential growth and then a phase of more rapid growth continuing until the present. For *Drosophila*, we followed the 3-epoch demographic model estimated by Sheehan and Song (2016), wherein the population experiences a protracted but moderate bottleneck followed by a nearly full recovery. Note that all of these demographic histories are single-population models with no gene flow. Our average mutation and recombination rates (*µ* and *r*, respectively) in humans were 1.2 × 10^−8^ (Kong *et al*. 2012) and 1.0 × 10^−8^, while in *Drosophila* these rates were set to 5 × 10^−9^ (Schrider *et al*. 2013; Assaf *et al*. 2017) and 2.3 × 10^−^, respectively (based on Comeron 2014). To allow the mutation rate to vary across simulated replicates, for each simulation we drew the rate uniformly from a range spanning a full order of magnitude and centered around the specified mean—thus mutation rates varied considerably among replicates but were constant across the simulated region within each replicate. In the interest of computational tractability we reduced our population sizes and simulation durations (in generations) 10-fold and 100-fold for the human and *Drosophila* simulations, respectively. Concordantly, mutation rates, recombination rates, and selection coefficients were increased by the same factor so that *θ* = 4*Nµ, ρ* = 4*Nr*, and *α* = 2*Ns* were unaffected by this rescaling.

### 3.4 Simulations of BGS in larger chromosomal regions

The simulation strategy above generated thousands of replicates of 1.1 Mb and 110 kb in length for humans and *Drosophila*. Because the impact of BGS can extend beyond these distances, we also simulated larger chromosomes of length 12.1 Mb and 1.21 Mb in humans and *Drosophila*. For these simulations we used the approach of the Real BGS and Real BGS–weak CNE scenarios described above, in which the locations of selected sites are based on randomly selected 12.1 Mb and 1.21 Mb regions of the human and *Drosophila* genomes, respectively. This approach allowed us to examine the impact of BGS on sweep detection when the region being examined is affected by both proximal and distal negatively selected sites spread across a chromosome. These simulations were carried out for each combination of demographic model and DFE described above.

### 3.5 Coalescent simulations of hitchhiking

Our goal was to compare the results of the BGS simulations described above to recent hard and soft selective sweeps. In order to rapidly simulate positive selection while conditioning on fixation of the adaptive allele, we used the coalescent simulator `discoal` (Kern and Schrider 2016). These simulations used the same demographic histories and average values of locus-wide *θ* and *ρ* as the forward simulations for BGS above. Again, *θ* varied uniformly across an order of magnitude from replicate to replicate. Rather than following a particular genetic map, *ρ* was drawn from a truncated exponential with a maximum value fixed to three times the mean—larger values of *ρ* require more memory and therefore sometimes cause the simulation to crash. This strategy allowed for variation in recombination rate while skewing toward lower rates. For each demographic history, we simulated 4000 examples of neutral evolution, hard sweeps occurring at the center of each of 11 adjacent equally sized windows partitioning the simulated chromosome, and soft sweeps at the center of each window.

Our soft sweeps consisted of selection on a previously neutral allele that is segregating at a specified frequency at a time at which it becomes beneficial and sweeps to fixation(Hermisson and Pennings 2005), rather than the alternative model of soft sweeps from recurrent adaptive mutations (Pennings and Hermisson 2006). Note that this model does not ensure that multiple independent lineages that harbor the adaptive allele at the onset of selection will participate in the sweep (i.e. it is possible that all lineages but one may go to extinction, making the outcome somewhat more similar to selection on a *de novo* mutation). We also note an alternative definition of hard and soft sweeps is commonly used in the literature, where a sweep is defined as hard if all sweeping lineages trace their ancestry to a single individual at the onset of selection and defined as soft otherwise (e.g. Hermisson and Pennings 2017). However, here we simply equate hard sweeps with selection on a *de novo* mutation and soft sweeps with selection on standing variation. For each hard and soft sweep replicate the selection coefficient of the beneficial mutation was drawn uniformly from between 0.0001 and 0.05, while the initial selected frequency for soft sweeps raged from between 0 and 0.05. The fixation time for each sweep was randomly chosen from between 0 and 200 generations ago, thereby modeling relatively recent selective sweeps (e.g. completing within the last ∼5000 years in humans). As with our forward simulations, our sample size was set to 100 haploid genomes.

### 3.6 Forward simulations with hitchhiking and background selection

We also sought to compare the influence of hitchhiking on diversity in regions with and without BGS. We therefore used forward simulations to model the Real BGS scenario described above while also conditioning on the recent fixation of an adaptive mutation near the center of the region. The results of these simulations were then compared to simulated hitchhiking events on otherwise neutrally evolving chromosomes. The procedure for this simulation was as follows: first the simulation runs up until a randomly selected time drawn from ∼ *U* (201, 5000) generations before the present, and the simulation state is saved. Next, the selected phase begins either by introducing a *de novo* beneficial mutation at the center of a randomly selected chromosome in the case of hard sweeps, or by changing the selection coefficient of the polymorphism nearest to the center of the chromosome having a frequency within a specified range in the case of a soft sweep. If the selected mutation is lost or does not reach fixation by the end of the simulation, the simulation is restarted from the point at which the state was previously saved, and this process repeats until fixation is achieved.

We sought to use the same uniform distributions as the coalescent simulations described above for the selection coefficient, fixation time, and initial selected frequency (in the case of soft sweeps). Because some combinations of these parameters are more likely than others to yield simulation replicates matching our acceptance criteria, we downsampled our set of completed replicates after splitting them into bins that were equally sized with respect to the fraction of each parameter range encompassed. For hard sweeps, we split our parameter ranges for the selection coefficient and fixation time into thirds, and drew an equal number of replicates from each bin in the resulting two-dimensional grid of nine parameter ranges. For soft sweeps we split our three parameter ranges (selection coefficient, fixation time, and initial selected frequency) into halves, and drew an equal number of replicates from each bin in the three-dimensional grid of 8 parameter ranges. The resulting distribution of accepted replicates is then somewhat similar to that produced under our coalescent simulations, although our binning procedure is fairly coarse and also reduces our total number of replicates considerably. We performed this procedure for both hard and soft sweeps with and without BGS under the Tennessen et al. (2012) model of European demography, using the same rescaling factor as above (0.1). The resulting number of replicates were as follows: 342 for hard sweeps on a neutrally evolving background, 104 for soft sweeps on a neutrally evolving background, 378 for hard sweeps on a background experiencing BGS, and 128 for soft sweeps with BGS.

### 3.7 Summary statistics and visualization

For each coalescent and forward simulation we calculated the following statistics: *π* (Nei and Li 1979; Tajima 1983), 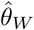 (Watterson 1975), 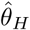 and Fay and Wu’s *H* (Fay and Wu 2000), Tajima’s *D* (Tajima 1989), the maximum derived allele frequency (max(DAF); Li 2011), the number of distinct haplotypes, *H*_12_ and *H*_2_/*H*_1_ (Garud *et al*. 2015), Kelly’s *Z*_*nS*_ (i.e. average *r*^2^; Kelly 1997), Kim and Nielsen’s *ω* (Kim and Nielsen 2004), and the variance, skewness, and kurtosis of the distribution of densities of pairwise differences between chromosomes. All of these can be calculated using `diploS/HIC` (Kern and Schrider 2018) in haploid mode. For our human simulations, these statistics were calculated both within 100 kb and 1 kb windows to visualize variation across coarse and fine scales, respectively. In *Drosophila*, these window sizes were 10 kb and 100 bp. The smaller window sizes produced plots that were quite noisy, so we further smoothed values by plotting running averages across 10 windows.

### 3.8 Classifying sweeps

We adopted a classification approach in order to ask how often our forward simulations resembled selective sweeps on the basis of the set of summary statistics described above. In particular, we used the `diploS/HIC` software in haploid mode to classify each simulated region as a hard sweep, a soft sweep, linked to a hard sweep, linked to a soft sweep, or neutrally evolving. A classifier was trained to discriminate among these five classes as follows. Note that none of these classes contain BGS, but this classifier can still be used to classify simulations with BGS, thereby revealing which of the five classes a BGS replicate most closely resembles.

Training was performed by first dividing the coalescent simulations described above into 11 windows. Hard and soft sweep simulations with a selected mutation located in the central window were then labeled as “hard” and “soft”, respectively, while those in other windows were labeled as “hard-linked” and “soft-linked”, respectively. Next, a balanced training set having 2000 examples of each of the five classes was constructed from these simulations, and a separate set of the same size was set aside for testing. We then trained our classifier using `diploS/HIC`’s `train` command with default parameters, before using the `predict` command to obtain classifications, which tell us which of the five classes above most closely resembles a given simulation outcome according to `diploS/HIC`. Three classifiers were trained in total: one for the human equilibrium model, one for the human model of European demography from Tennessen et al. (2012), and one for the *D. melanogaster* model of African demography from Sheehan and Song (2016). Each classifier was then applied to additional simulations generated under the corresponding demographic model as described in the Results.

### 3.9 Availability of Data and Code

All code for generating forward and coalescent simulations, calculating and visualizing summary statistics, training and applying `diploS/HIC` are available at https://github.com/SchriderLab/posSelVsBgs. In addition, all simulated data, along with statistics from each replicate in both text and graph form, are available at https://figshare.com/projects/posSelVsBgs/72209.

## 4 Results

### 4.1 Simplistic models of background selection in humans can resemble selective sweeps

We begin by simulating constant-sized populations with mutation and recombination rates matching estimates from the human genome, and comparing average patterns of diversity after a selective sweep to those from two different models of BGS. The first model of BGS that we examined (dubbed “Central BGS”) includes a 50 kb coding region in the center of the simulated locus, flanked by non-functional DNA on either side occupying the remainder of the 1.1 Mb locus—note that similar models have been used to describe the expected patterns of polymorphism under BGS (e.g. Mughal and DeGiorgio 2018). In the second model (“Real BGS”), for each replicate simulation a 1.1 Mb window was randomly selected from the human genome, and our simulation was designed to match several features of this genomic window including the locations of selected sites (exons and CNEs) and the recombination landscape (Methods). Although parameter values varied across replicates, the overall locus-wide mean population-scaled mutation and recombination rates, *θ* and *ρ*, were set to roughly match values expected in a human population of effective size 10,000 and a total locus size of 1.1 Mb. Note that for our BGS simulations, polymorphism at selected sites is included in our observations—our results therefore reflect the action of direct purifying selection as well as BGS. Selective sweeps were generated via coalescent simulation, while the BGS scenarios were modeled via forward simulation (Methods).

In Figure 2 we show levels of diversity as measured by three estimators of *θ* within simulated regions experiencing different modes of linked selection (averaged across 1000 replicates). We see that hard selective sweeps produce a large valley of diversity at the center of the simulated region with a gradual recovery toward equilibrium moving away from the selected site (Figure 2A), as expected (Maynard Smith and Haigh 1974). Note that this valley is more pronounced for *π* than for *θ*_*W*_, due to the expected deficit of intermediate-frequency alleles produced by a hitchhiking event (Braverman *et al*. 1995). *θ*_*H*_ on the other hand is elevated in the regions flanking the selective sweep due to the excess of high-frequency derived alleles that escaped the sweep via recombination (Fay and Wu 2000). For soft sweeps (Figure 2B), we see a qualitatively similar pattern but with a less pronounced valley of diversity. Again, *π* is lower than *θ*_*W*_ in the simulated region, indicating a deficit of intermediate-frequency alleles. It has been observed that soft sweeps with a fairly high initial selected frequency (e.g. 5% or more) can sometimes yield an excess of intermediate-frequency alleles (Teshima *et al*. 2006; Schrider *et al*. 2015), but here our initial selected frequency is constrained to values ≤5%, so closer concordance between hard and soft sweeps is expected. Additional statistics summarizing information about the site frequency spectrum, haplotype diversity, and linkage disequilibrium are shown in Figure 3. These statistics show patterns concordant with expectations under a sweep: we observe peaks in the number of high frequency derived alleles (Fay and Wu 2000; Hahn 2018) and in LD (Kim and Nielsen 2004) in regions flanking the sweep, decaying haplotype homozygosity with increasing distance from the selected site (Garud *et al*. 2015), and characteristic spatial patterns of the variance, skewness, and kurtosis of pairwise diversity around the sweep (Kern and Schrider 2018).

In the Cental BGS scenario (Figure 2C) we observe a strong localized reduction in diversity caused by direct selection against deleterious mutations, and then a rapid increase in diversity as we move away from the selected sites. Still, these diversity levels are somewhat reduced relative to the neutral expectation due to their linkage with the selected region, and recover gradually as we move further away as expected under BGS (Charlesworth *et al*. 1993). There are also apparent changes to the site frequency spectrum in the selected region (e.g. reduced Tajima’s *D* and max(DAF)), although these quickly recover toward neutral expectations with increasing distance from the selected region. Thus, while the average realization of this particular scenario does not perfectly match the predictions of a selective sweep, it is consistent with the possibility that regions experiencing BGS may commonly be mistaken for sweeps, especially if negatively selected sites are also examined—as may typically be the case when scanning for positive selection in practice.

Finally, in Figure 2D we show the mean values of the three estimators of *θ* across the Real BGS simulations, wherein each replicate draws its recombination map and locations of selected sites from a random region of the human genome. We see that diversity is somewhat reduced in these simulations relative to the neutral expectation due to the combination of direct and linked negative selection. Because each replicate models a different genomic region, there is no consistent spatial pattern shown in Figure 2D, but this does not mean that individual replicates in this set do not resemble selective sweeps—we examine this possibility below.

### 4.2 Expected patterns of diversity produced by BGS in particular regions of the human genome

In the previous section we examined the average values of different summary statistics under four different evolutionary models (Figures 2–3), including one model of BGS where the chromosomal locations of exons and CNEs as well as the recombination map were chosen to match randomly selected regions in the human genome. Because these regions may differ dramatically in the number and locations of selected sites, and thus the expected impact of selection on diversity, rather than looking at the average across regions, a more useful question to ask is whether any particular region’s measures of diversity are expected to resemble selective sweeps. We examine this in Supplementary Figures 1–10, where we show the values of 15 summary statistics calculated from sets of simulations each modeled after a particular randomly chosen region of the human genome, with 1000 replicates for each region. These plots also show the density of exonic and conserved non-coding sites in 100 kb windows, revealing the concordance between the peaks and valleys in the density of selected sites and the average values of the summary statistics. Among these 10 examples, we see considerable variation in the number and arrangement of selected sites along the chromosome. There are corresponding differences in the mean values of summary statistics from region to region, and generally across windows within a region we see subtle shifts in the amount of nucleotide diversity that coincide with changes in the density of selected sites (i.e. peaks in conserved elements correspond to slight dips in *π*). However, in none of these regions do the expected patterns of summary statistics resemble a selective sweep, or even the Central BGS scenario. An examination of the arrangements of selected sites along these chromosomal regions reveals why this is so: in none of these 10 regions do we see a high density of selected sites in the center flanked by largely unconstrained sequence. Thus these results suggest that scenarios of background selection that are most likely to resemble selective sweeps may not be appropriate models for the typical manner in which BGS shapes diversity across the human genome. We investigate this possibility more systematically in the following section.

### 4.3 BGS rarely produces patterns of diversity resembling selective sweeps in equilibrium populations

In the previous section we examined simulated data based on 10 randomly selected 1.1 Mb regions of the human genome, finding that none are expected to produce patterns of variation mimicking a recent hitchhiking event. However, the human genome is large, consisting of ∼3000 such regions. Thus, even if a small minority of regions have an arrangement of selected sites and a recombination map that lend themselves to producing large valleys of diversity at their center, then BGS could still result in a large number of regions somewhat resembling selective sweeps. We sought to examine this directly by asking how many of our simulated examples from the Real BGS set are mistaken for sweeps on the basis of their spatial patterns of population genetic summary statistics. To do this, we used the `S/HIC` framework (Schrider and Kern 2016), which represents a genomic region as a large vector of population genetic summary statistics calculated in and normalized across each of a number of windows within this region (Methods); we refer to this set of statistics as our feature vector. `S/HIC` then classifies the central window of this region into one of five distinct evolutionary models: a hard sweep (i.e. a hard selective sweep recently occurred at the region’s center), a soft sweep, linked to a hard sweep (i.e. a hard sweep recently occurred within or near the region, but not within the central sub-window), linked to a soft sweep, or evolving neutrally (i.e. no recent sweep in the vicinity of the region). This inference is made via supervised machine learning: we first train a classifier on the basis of feature vectors calculated from genomic regions whose true class is known prior to applying the trained classifier to data whose true class may be unknown. In our case, the training data are obtained via coalescent simulation of regions with a sweep in a center, surrounded by unselected sequence (Methods). Because of the design of its feature vector, `S/HIC` is well suited for determining whether a given genomic window resembles a selective sweep or not on the basis of its spatial patterns of genetic variation. We note that there are similar approaches that may be equally suitable for this task (e.g. Lin *et al*. 2011; Mughal and DeGiorgio 2018).

After training our `S/HIC` classifier, we applied it to the 1000 replicates from our Central BGS and Real BGS sets of forward simulations (i.e. the same data examined in Figure 2D). First, we assessed `S/HIC`’s ability to perform the task for which it was trained—discriminating between selective sweeps, regions linked to selective sweeps, and neutrally evolving regions (top 5 rows of Figure 4). Overall the classifier performed quite well, although discrimination between hard and soft sweeps is difficult for both the sweep and sweep-linked classes; this is not unexpected given that our soft had a fairly low initial selected frequency (≤5%), making them more similar to hard sweeps where the initial frequency is 1*/*2*N*. Moreover, here we equate soft sweeps with selection on standing variation regardless of the number of independent copies that participate in the sweep (Methods), raising the possibility that in some cases only a single ancestral copy will reach fixation. However our primary concern is the extent to which sweeps of any type can be distinguished from alternative evolutionary models. Importantly, we see that 4% of neutrally evolving regions are misclassified as selective sweeps (all as soft; Figure 4)—these results are based on forward simulations but similar numbers are obtained when we use a test set coalescent simulations generated in the same manner as those used to train `S/HIC`. Thus, due to the stochasticity of the evolutionary process, despite the vast difference in the expectations between sweep models and neutrality, we can expect to occasionally see neutrally evolving regions whose spatial patterns of genetic diversity resemble selective sweeps closely enough for `S/HIC` to misclassify them.

**Figure 4:**
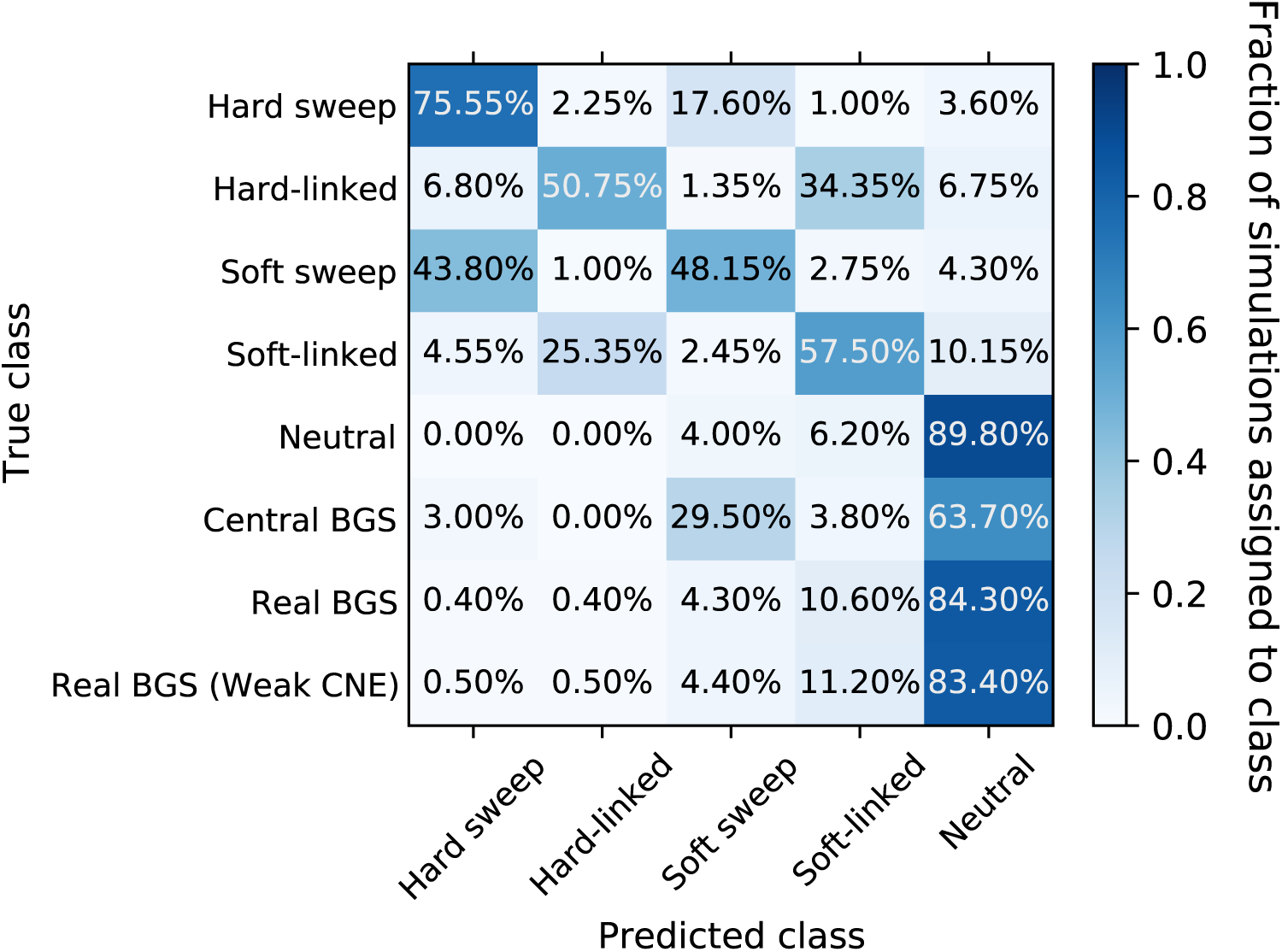
Confusion matrix showing the fraction of constant-size simulations assigned to each class by `S/HIC`. The *y*-axis shows the true evolutionary model of the simulated region, and the *x*-axis shows the model selected by `S/HIC`. Thus a given row shows the fraction of simulated examples of a given scenario that were assigned to each of `S/HIC`’s five classes. In the top five rows, the diagonal corresponds to correct classifications, while in the bottom three rows (BGS scenarios) there is no possible correct classification and our concern is how often a region is misclassified as a sweep. Note that the examples with positive selection were simulated with `discoal` while those with negative selection or neutral evolution were generated with `fwdpy`

Next we assessed `S/HIC`’s behavior on BGS models not included in training (bottom 3 rows of Figure 4), first asking how often examples in our Central BGS set were mistaken for selective sweeps by `S/HIC`. Perhaps unsurprisingly given the sharp valley of diversity observed in Figure 2C, we find that 32.5% of these simulated regions are classified as sweeps by `S/HIC`, with the majority classified as soft sweeps (29.5% classified as soft versus 3% as hard). However, when examining the Real BGS simulations, which are designed to more accurately model BGS in the human genome, we find that 4.7% of examples are classified as sweeps (4.3% as soft and 0.4% as hard)—similar to the corresponding fraction of neutrally evolving examples (*P* = 0.51; Fisher’s exact test). When simulating weaker selection on CNEs than on protein-coding exons, we again find no significant elevation in the rate of false sweep calls (4.9% of of examples classified as sweeps; *P* = 0.39). However, we note that the Real BGS models do result in regions being classified as affected by linked soft selective sweeps (i.e. the soft-linked class) at a substantially higher rate than are neutral regions (∼11% for Real BGS models versus 6.2% under neutrality). In addition to running our classifier on each simulated replicate, we have created plots similar to Supplementary Figures 1–10 but rather than showing the mean values of each statistic we plot each simulation replicate separately. Readers curious about the extent of variability in patterns across individual realizations of each of our simulated scenarios—which can be considerable—may wish to explore these plots of each individual simulation (available at https://figshare.com/projects/posSelVsBgs/72209).

Thus far our simulations do not support the claim that BGS frequently alters diversity in a manner consistent with selective sweeps. This may imply that, at least in the case of paramterizations relevant for the human genome, BGS should be readily separable from models of selective sweeps by examining summaries of variation taken across a large chromosomal region. However, up to this point we have only considered a constant-size population. Given that non-equilibrium population dynamics can have a profound impact on genetic diversity and the effect of BGS (Torres *et al*. 2018, 2019) we examine two such models in the following sections.

### 4.4 Selective sweeps and background selection are readily distinguishable in the presence of drastic population size change

To this point we have shown that BGS and sweep models are readily distinguishable in simulated constant-sized populations with genomic regions modeled after those randomly selected from the human genome. It is known that humans populations have experienced a number of demographic changes that have reshaped patterns of diversity genome-wide (Gravel *et al*. 2011). These include recent explosive population growth (Tennessen *et al*. 2012) and in non-African populations a severe bottleneck associated with the migration out of Africa (Marth *et al*. 2004). Thus if our goal is to model BGS in humans we should consider the effects of dramatic population size changes. We therefore repeated all of the analyses described above under a model of European population size history (Tennessen *et al*. 2012).

Panels A and B of Figure 5 show that selective sweeps under the European model produce a fairly similar pattern to sweeps in a constant population (Figure 2), with one noticeable difference in that *θ*_*H*_ is depressed rather than elevated in regions flanking the sweep (though it remains considerably higher than *π*). Again, we see a sharp dip in diversity around the selected region in the Central BGS model (Figure 5C) and on average a global reduction in diversity in the Real BGS model (Figure 5D). When comparing the values of a larger set of population genetic statistics across the full 1.1 Mb region, we see that on average the Central BGS model bears a passing resemblance to sweeps for some statistics but not others, echoing our results from the constant population size case. (Supplementary Figure 11).

**Figure 5:**
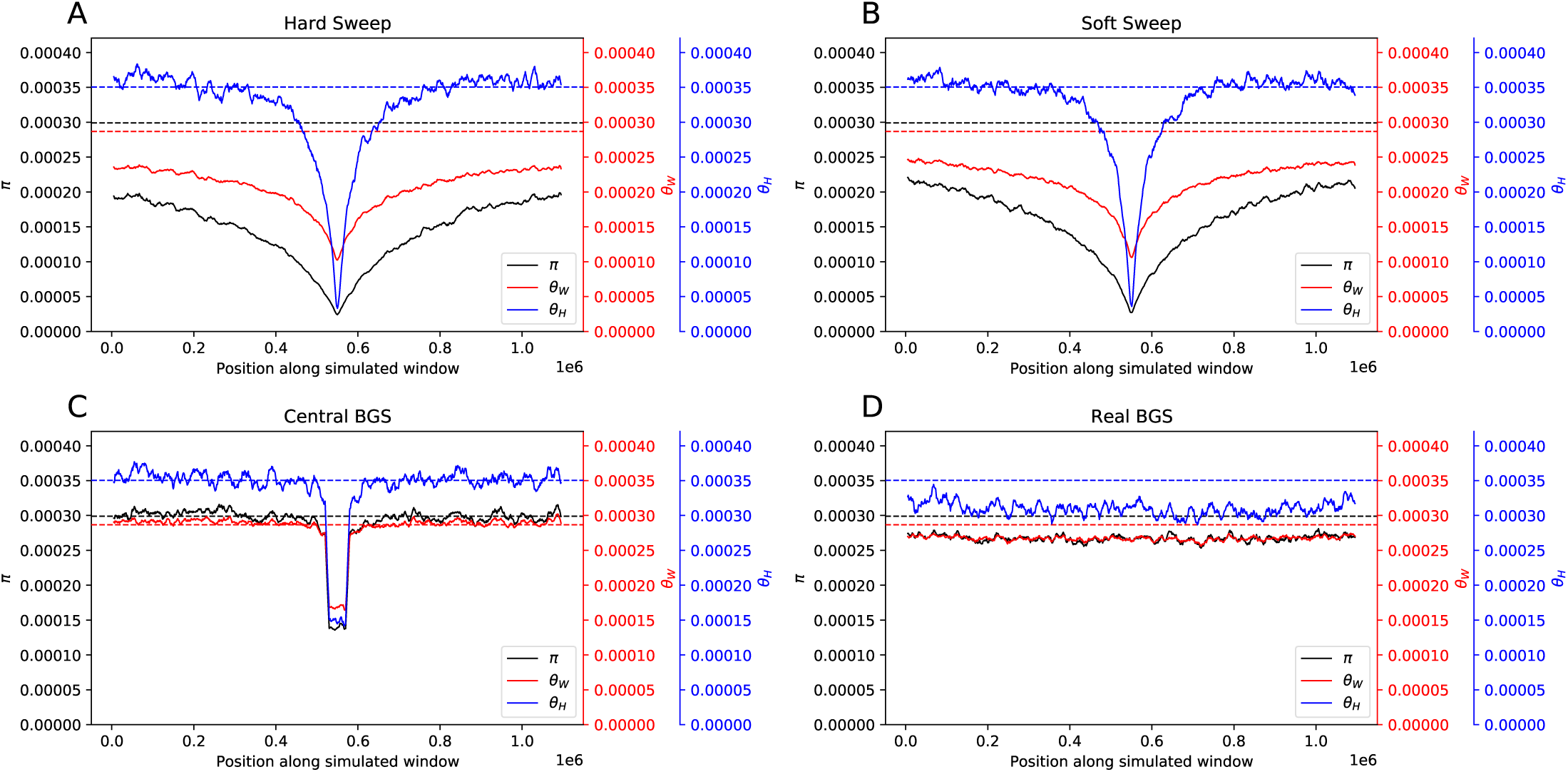
Patterns of diversity produced by different models of linked selection under Tennessen et al.’s model of European population size history (Tennessen *et al*. 2012). (A) Values of three estimators of *θ* in regions experiencing a recent hard sweep at the center of the window. (B) Values of these estimators in regions with a recent soft sweep at the center. (C) Values in regions where purifying selection is acting on a central 50 kb element. (D) Values in regions with purifying selection acting on elements distributed across the chromosome in a manner that mirrors the locations of exons and CNEs in randomly selected 1.1 Mb regions in the human genome. Each panel shows windowed statistics calculated as described in the Methods and averaged across 1000 simulated replicates, and the mean value of each statistic across the neutral forward simulations is also shown as a dashed line.

We also re-examined the same 10 randomly selected genomic regions shown in Supplementary Figures 1–10, this time simulating BGS under Tennessen et al.’s European model (Supplementary Figures 12–21). Again, none of these 10 regions show the appearance of a sweep. To more formally ask how often examples of each of our scenarios resemble hitchhiking events, we trained a `S/HIC` classifier under the Tennessen et al. model (Methods) and recorded the number of simulations that were misclassified as a selective sweep (Figure 6). Population bottlenecks are expected to produce sweep-like signatures (Simonsen *et al*. 1995; Jensen *et al*. 2005; Nielsen *et al*. 2005), and we do find that under this demographic model we observe a slightly higher false positive rate for neutrally evolving regions than under constant population size (6.2% in total versus 4% under constant population size; *P* = 0.033). In addition, we do have greater difficulty distinguishing between hard and soft sweeps, perhaps because population bottlenecks during a sweep can reduce diversity among chromosomes harboring the beneficial allele, thereby “hardening” the sweep (Wilson *et al*. 2014). As observed under equilibrium demographic history, the false positive rate is considerably higher under the Central BGS model than under neutrality (0.9% and 22.7% of simulations classified as hard and soft, respectively Figure 4). Under the Real BGS model, the false positive rate is similar to that under neutrality (5.8% in total, with 5.7% and 0.1% of simulations classified as hard and soft, respectively; *P* = 0.78 for the comparison with neutrality); we find similar results when simulating regions with weaker selection on CNEs (5.6% false positives, with 4.9% classified as hard and 0.7% as soft; *P* = 0.64 when compared with neutrality). In sum, under neither Real BGS model do we see an excess of regions classified as being linked to a sweep. Thus, under realistic arrangements of selected sites, models of BGS in humans do not appear to produce signatures of selective sweeps even in the context of severe population size change.

**Figure 6:**
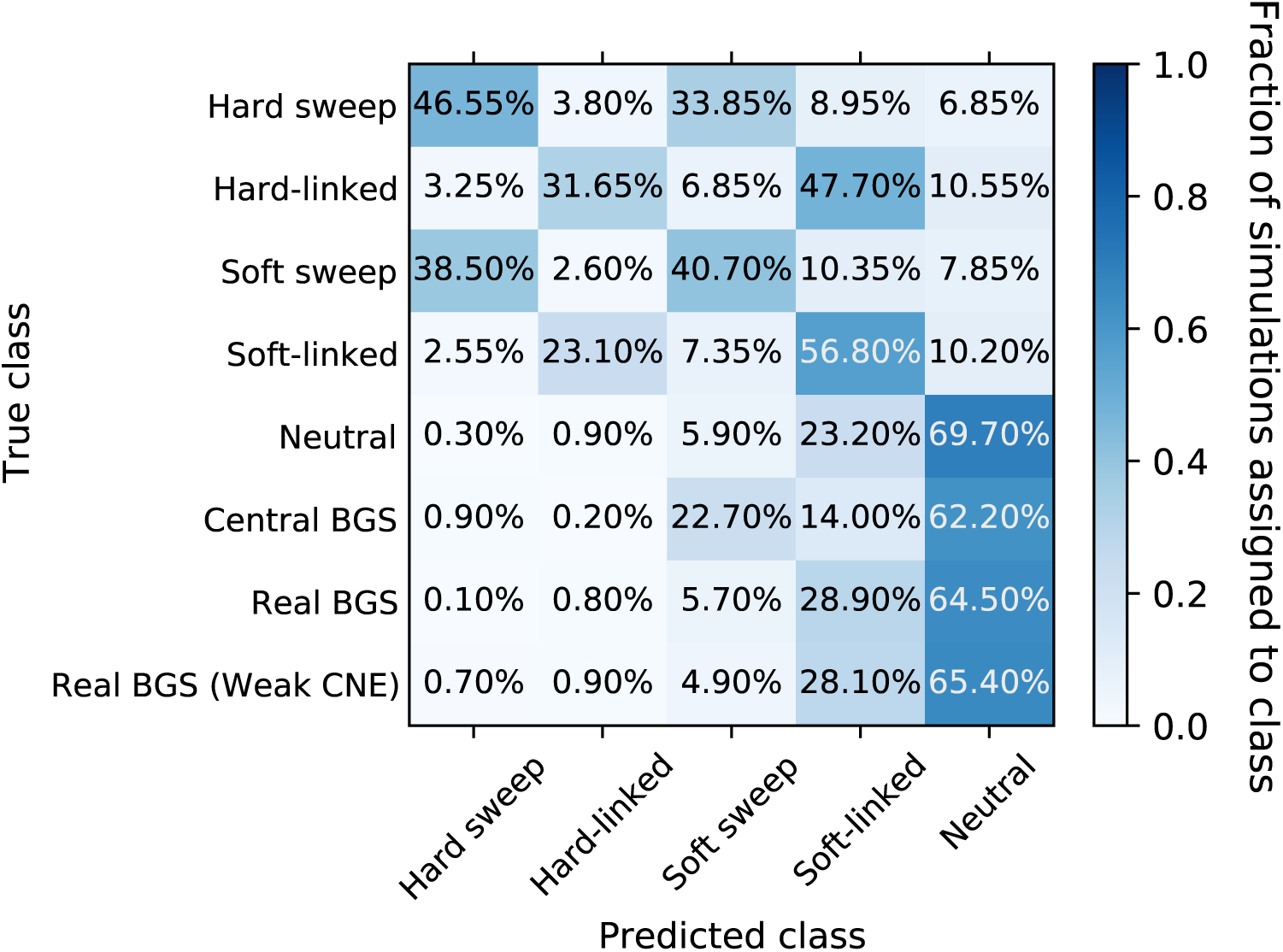
Confusion matrix showing the fraction of simulations under the Tennessen et al. (2012) model of European demography assigned to each class by `S/HIC`, following the same schema as Figure 4.

### 4.5 BGS rarely mimics sweeps in *Drosophila*

Thus far our results are based on simulations modeled after the human genome which has a low density of genes and conserved DNA (∼5%; Siepel *et al*. 2005) compared to more compact genomes. To examine the impact of BGS in a genome with a higher density of conserved elements, we simulated BGS in regions modeled after 110 kb windows in the *Drosophila melanogaster* genome (see Methods), in which over 50% sites show evidence of purifying selection (Andolfatto 2005; Halligan and Keightley 2006). For these simulations we used Sheehan and Song’s (2016) three-epoch demographic model of a Zambian population sample (Lack *et al*. 2015). In Figure 7 we again show values of three estimators of *θ* under our four difference scenarios. For this demographic history, there is a more pronounced difference between hard and soft sweeps (Figure 7A–B). This may be a consequence of the larger effective population size (*N*_*e*_) for *Drosophila*, which yields a much stronger effect observed for hard sweeps than under previous scenarios, while soft sweeps contain a drift phase whose duration is constant (in coalescent units) across values of *N*_*e*_. Under the Central BGS scenario (Figure 7C), we again see a strong reduction in diversity in the immediate vicinity of the selected sites flanked by a rapid recovery. In the Real BGS scenario (Figure 7D) we again do not see any spatial pattern on average, as expected given our random sampling across loci. However we do see a larger mean reduction in variation relative to expectations in the absence of selection than seen in the human scenarios due to the denser placement of selected sites in *Drosophila*. The average patterns of additional summary statistics are shown in Supplementary Figure 22; these statistics show a strong hitchhiking effect for hard sweeps (a valley of nucleotide diversity, a plateau of LD, an SFS skewed towards low- and high-frequency derived alleles, etc.), and a somewhat different pattern for soft sweeps (e.g. elevated Tajima’s *D*, a fairly high ratio of *H*2*/H*1). Although for Central BGS the depth of the valley of diversity closely resembles that of soft sweeps, as expected there is no spatial pattern on average for any statistics under the Real BGS scenario.

**Figure 7:**
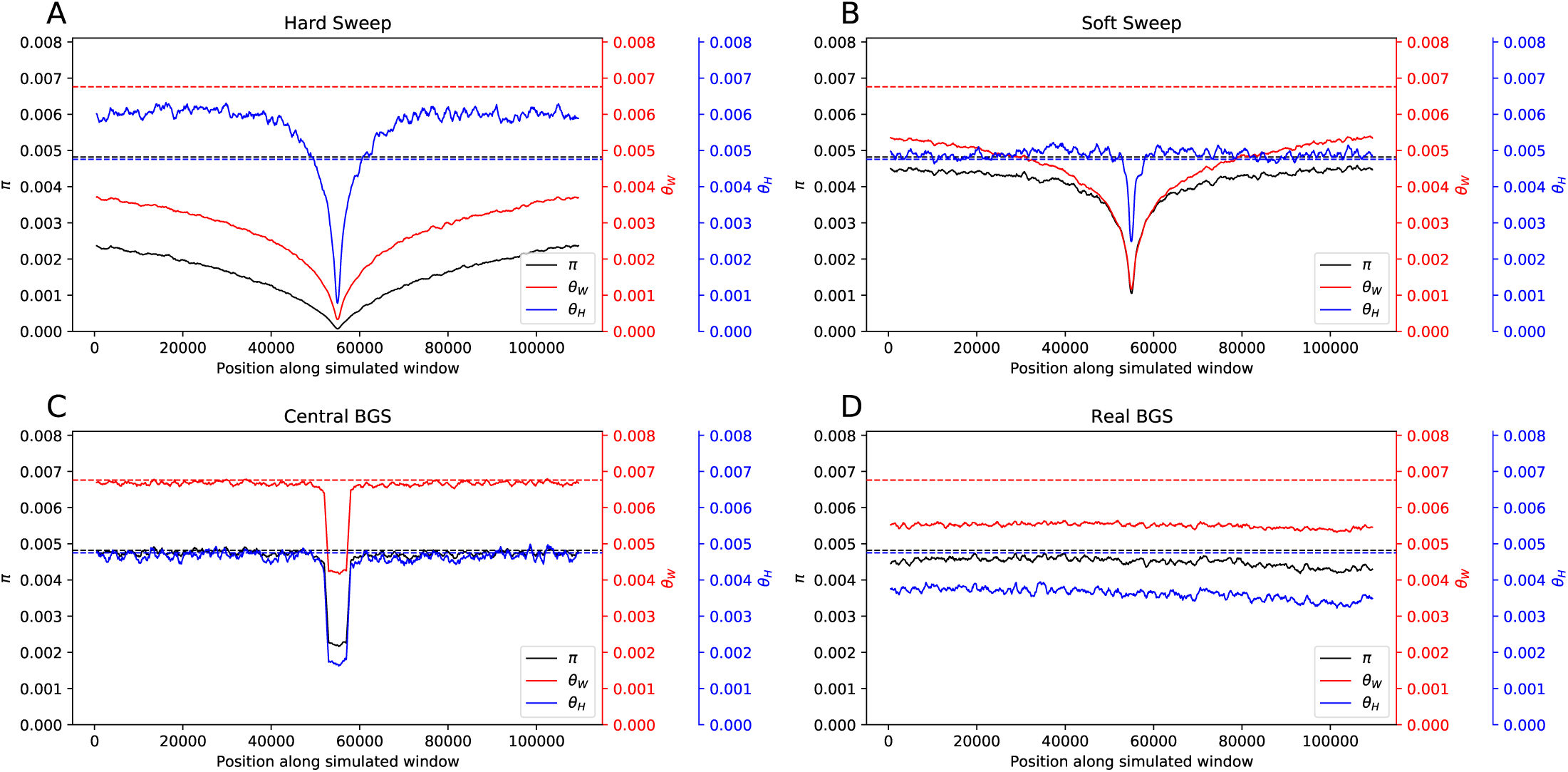
Patterns of diversity produced by different models of linked selection under the Sheehan and Song (2016) model of African *Drosophila melanogaster* population size changes. (A) Values of three estimators of *θ* in regions experiencing a recent hard sweep at the center of the window. (B) Values of these estimators in regions with a recent soft sweep at the center. (C) Values in regions where purifying selection is acting on a central 5 kb element. (D) Values in regions with purifying selection acting on elements distributed across the chromosome in a manner that mirrors the locations of exons and CNEs in randomly selected 110 kb regions in the *Drosophila* genome. Each panel shows windowed statistics calculated as described in the Methods and averaged across 1000 simulated replicates, and the mean value of each statistic across the neutral forward simulations is also shown as a dashed line.

We examine expected values of summary statistics in 10 randomly selected individual regions in Supplementary Figures 23–32. Here we see a much more conspicuous relationship between summaries of diversity and the density of conserved elements, in part because this density is an order of magnitude higher than in our human simulations. Again, none of our 10 randomly selected regions at all resemble a selective sweep. Using a `S/HIC` classifier trained on coalescent simulations under the Sheehan and Song model (Methods), we see that no neutrally evolving regions are misclassified as sweeps (Figure 8); this is perhaps unsurprising given the much stronger signatures shown in Figure 7 than in either human model we examined. A modest fraction of the Central BGS cases are misinferred to contain sweeps (3.7% in total, with 2.3% hard and 1.4% soft), but for Real BGS these false positives occur more rarely (1%, with 0.1% hard and 0.9% soft; these fractions are 0.5% hard and 0.2% soft when selection on CNEs is weaker). Unlike in the human scenarios, the Real BGS simulations do result in an excess of sweep calls (*P* = 0.0019 and *P* = 0.015 in the standard and weak-CNE Real BGS scenarios, respectively), though this effect is quite modest (false positive rate ≤1% in either case). As in the human scenarios, we also note a sizeable fraction of simulations of the Real BGS scenario assigned to `S/HIC`’s soft-linked class (16.7% and 18.9% for the standard and weak-CNE real BGS examples, versus 2.2% under neutrality).

**Figure 8:**
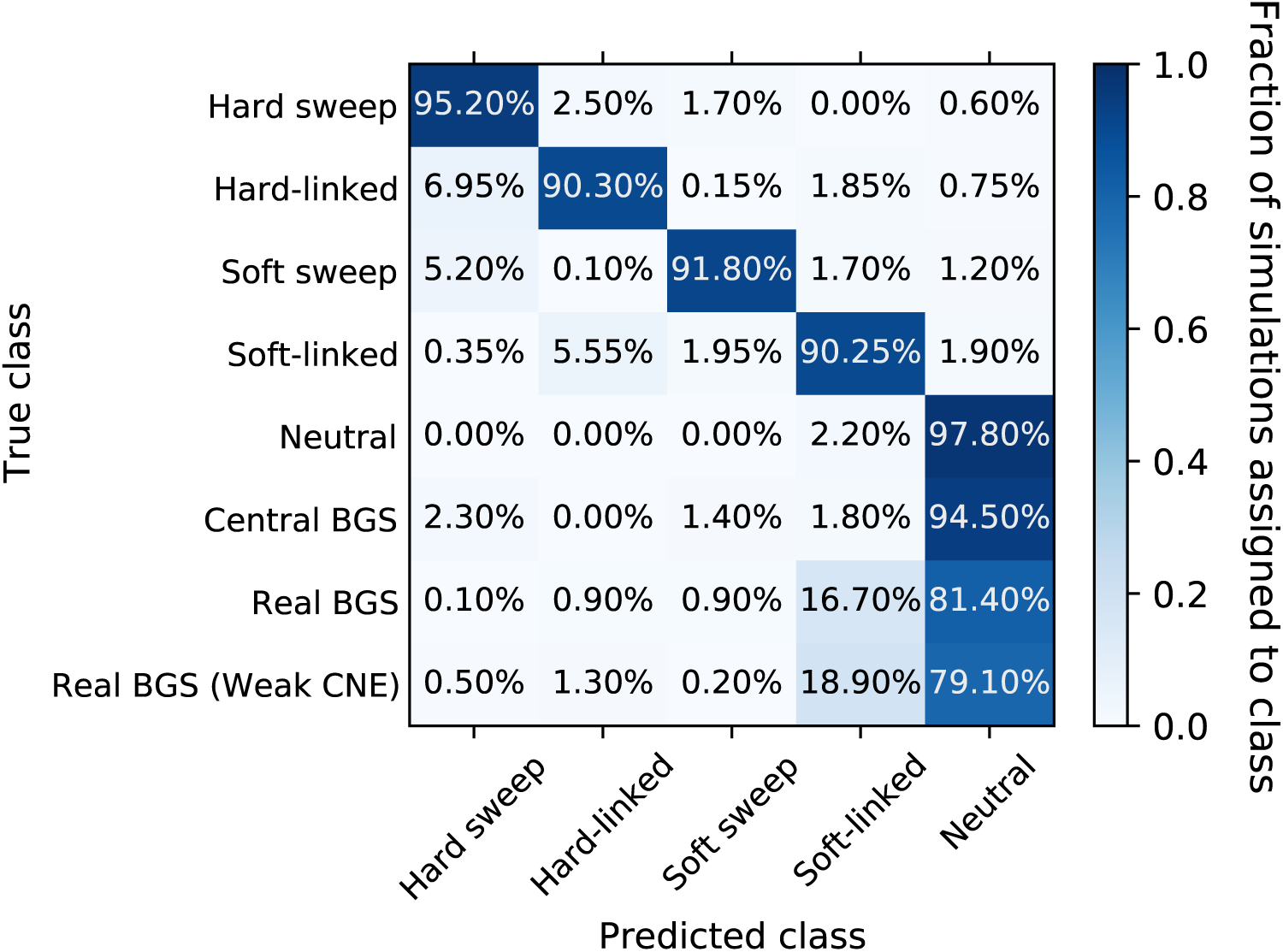
Confusion matrix showing the fraction of simulations under the Sheehan and Song (2016) model of African demography in *Drosophila* assigned to each class by `S/HIC`. The *y*-axis shows the true evolutionary model of the simulated region, and the *x*-axis shows the model selected by `S/HIC`. Thus a given cell shows the fraction of simulated examples the belong to a given class were assigned to each of `S/HIC`’s five classes. In the top five rows, the diagonal corresponds to correct classifications, while in the bottom two rows (BGS examples simulated via fwdpy) there is no possible correct classification and our concern is how often a region is misclassified as a sweep.

### 4.6 Dominance of deleterious mutations

We also examined the impact of the dominance coefficient of deleterious mutations on the propensity of regions experiencing BGS to be mistaken for selective sweeps. As shown in Supplementary Figure 33, there does not appear to be a strong relationship between dominance and the fraction of Real BGS simulations classified as a sweep by `diploSHIC`. In our simulated constant-sized human populations, the fraction of spurious sweep calls (either hard or soft) hovers around 5%, with no significant difference across dominance values and no discernible trend with increasing dominance. Similar results are observed under the European model, where the fraction of sweep calls ranges between 4%–6%, and the African *D. melanogaster* model, where this fraction is ∼1% or less in all cases, again with no significant difference among dominance classes. These results imply that, at least for models of BGS in which every deleterious mutation has the same dominance coefficient, patterns of variation are unlikely to resemble those expected under recent hitchhiking events regardless of that coefficient’s value.

### 4.7 Properties of regions misclassified as selective sweeps

Although we find that the fraction of Real BGS simulations misclassified as sweeps is not dramatically different from that of neutral simulations, it may be useful to ask whether the propensity of a genomic region under BGS to produce a signature of hitchhiking can be predicted *a priori* from the arrangement of selected sites and the recombination rate. In Table 1, we show Spearman’s correlation coefficients between the probability that a given simulation contains a sweep according to `S/HIC` (represented by the sum of the `diploSHIC`’s posterior probability estimates for the hard and soft classes) and the number of selected sites in the central window, the number of selected sites in all other windows, and the total recombination rate of the simulated region. We calculated these correlations for each combination of species, demographic history, and strength of selection on CNEs in our simulated dataset. For each data set we observe a significant negative correlation between the recombination rate and `SHIC`’s predicted posterior probability that the region is a sweep. In *Drosophila*, we also observe a significant correlation between the number of selected sites in the central window and the posterior probability of a sweep in the weak-CNE scenario. Overall our results suggest that there may be some power to predict which regions are most likely to be produce spurious sweep-like signatures—as one might expect, such regions have lower crossover rates and a greater density of selected sites in the central window (although the latter was significant in only one of our simulated scenarios).

**Table 1:**
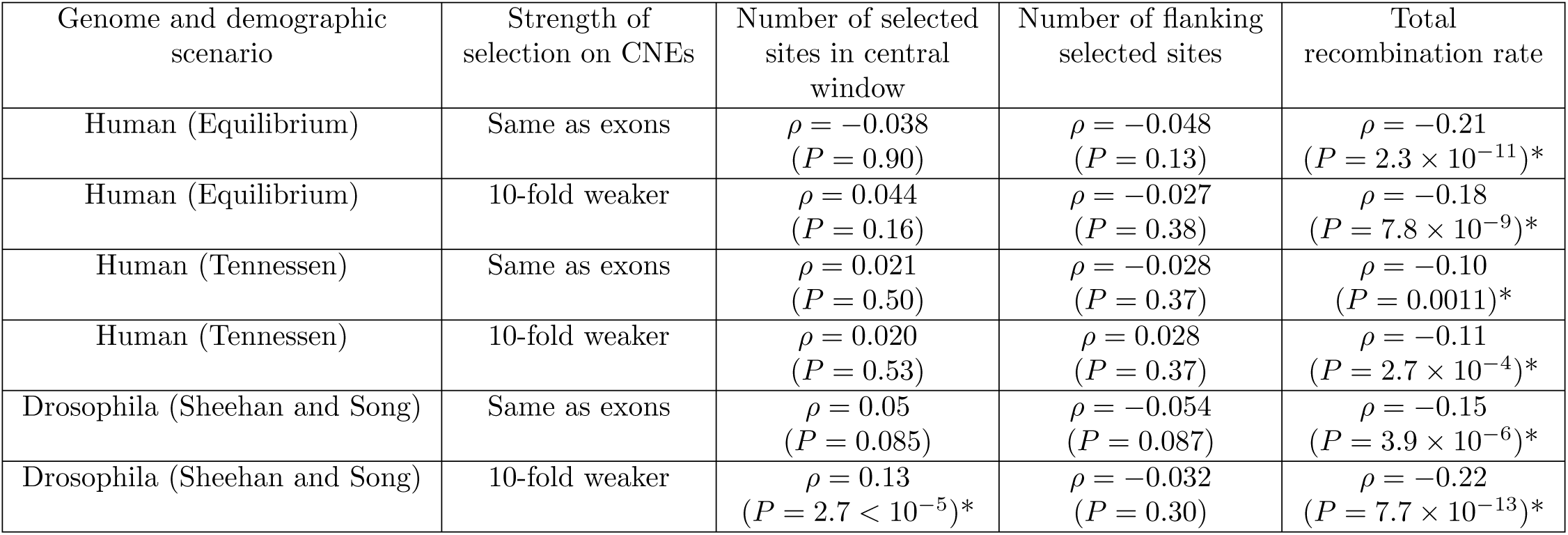
Spearman’s correlation coefficients between various properties of the simulated region and the sum of the posterior probabilities for `diploSHIC`’s hard and soft classes. Uncorrected *P*-values are shown with asterisks marking correlations that are significant with *α* = 0.05 after Bonferroni correction. All simulations were under either the Real BGS model or the Real BGS model with weaker selection on CNEs.

To more closely examine the impact of recombination rate on our results, for each demographic model we binned all of the Real BGS simulation according to mean recombination rate across the simulated chromosome. Five bins were used: one bin reserved for the fairly small number of replicates with a recombination rate of zero, and the four quartiles among simulations with a non-zero recombination rate. In Figure 9, we show the fraction of examples misclassified as selective sweeps of either type by `S/HIC` for each recombination bin and in each case compare to the misclassification rate for neutral simulations (rightmost bar). The results of this same analysis for the Real BGS–Weak CNE model is shown in Supplementary Figure 34. We see that for most of the human recombination rate bins there is no significant elevation of the false positive rate relative to neutral simulations, although this may in part be due to a inadequate statistical power, with one exception being the no-recombination bin for the weak CNE simulations (*P* = 0.04), and a trend toward higher false positive rates in replicates with less recombination is seen in the Human equilibrium simulations. However, in *Drosophila*, the effect of recombination is clear: regions with no recombination or in the lowest non-zero recombination rate bin account for the majority of false sweep calls by `S/HIC`, and outside of these two bins the false positive rate is ∼0. Thus, the small but significant elevation in false positive rate produced by BGS in *Drosophila* seems to be driven entirely by low- or non-recombining regions.

**Figure 9:**
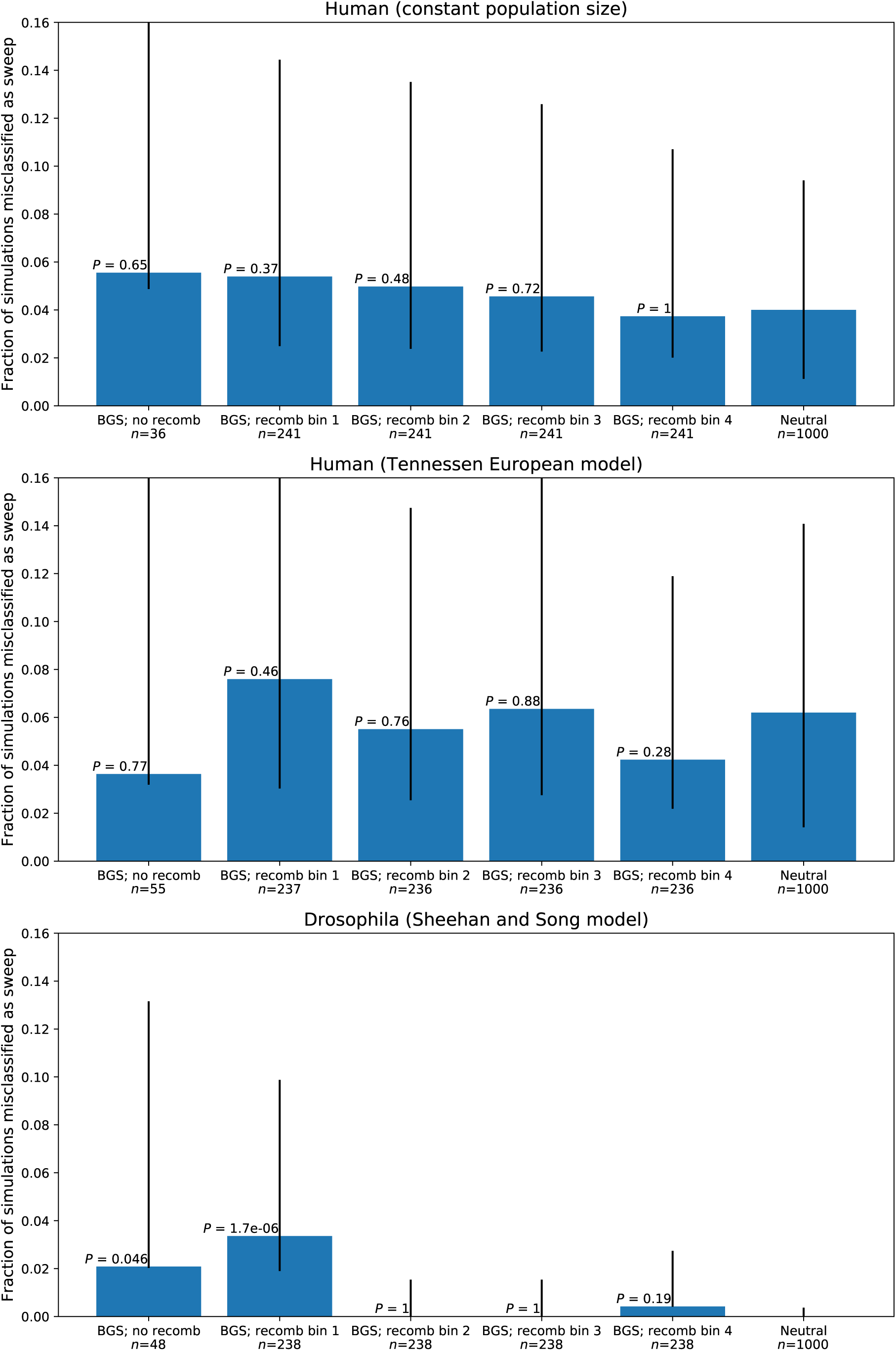
The fractions of simulations of our Real BGS model misclassified as a sweep of either type for each demographic model, shown after binning our data. The rightmost bar shows the fraction of misclassified neutral simulations for comparison, and all *P*-values show significance of the comparison with neutrality. The error bars show the 95% binomial confidence intervals.

### 4.8 BGS does not mimic sweeps in larger simulated chromosomes

Until this point we have limited our analysis to simulated chromosomes 1.1 Mb and 110 kb in length for humans and *Drosophila*, respectively—these lengths were chosen to match the region size that we trained `S/HIC` to examine. Although this allowed us to simulate thousands of replicates, the impact of BGS may be smaller in such simulations because they do not include the effect of selection in more distant linked regions which also influence diversity (Comeron 2014). Therefore, we simulated much larger (more than 10-fold larger than those described above; see Methods), with a smaller number of replicates (100 per demographic model-DFE combination) due to the greater computational demands of these simulations. These chromosomes were 12.1 Mb and 1.21 Mb in humans and Drosophila—eleven times the length of our original simulations. Average nucleotide diversity in these simulations was qualitatively similar to that of the smaller simulations: *π* per site in the central 1.1 Mb window was 4.1 × 10^−4^ and 4.4 × 10^−4^ averaged across all small-scale and large-scale human equilibrium simulations, respectively, and 2.7 × 10^−4^ and 2.6 × 10^−4^ for the small- and large-scale simulations under the Tennessen et al. (2012) European model; in the central 110 kb window of the *Drosophila* simulations average *π* was 0.0045 and 0.0044 in the small- and large-scale simulations. This suggests that the simulations used in the preceding sections may be adequate for addressing the similarity of BGS and hitchhiking models, despite the relatively small chromosomes being modeled, perhaps because the impact of selection on additional linked sites is relatively small compared to the combined effect of direct and linked selection within the focal window. Moreover, we do not observe a significantly elevated false positive rate in the large-scale simulations when examining the central window within the chromosome (10). In our human equilibrium model, we observe a nominal increase in the false positive rate when switching from smaller to larger chromosomes (4.7% of BGS windows misclassified as sweeps in small-scale simulations versus 5% in large-scale simulations), although this is not significant (*P* = 0.81). In the Tennessen et al. European model, we actually see a smaller false positive rate in our large-scale simulations (5.8% versus 2%), but again this difference is not significant (*P* = 0.16). Similarly, in our *Drosophila* simulations, we see no significant difference in false positive rates between our small- and large-scale simulations (1% versus 0%; *P* = 0.61).

Our large-scale simulations allow us to examine another potential source of bias in our results: because we are identifying false sweep signatures using `S/HIC`, which looks for the pattern of diversity consistent with a sweep at the center of its focal window, one concern may thus be that BGS produces sweep-like signatures occur fairly often, but rarely with the epicenter at the center of the region examined by `S/HIC`. Were this the case, `S/HIC` would be underpowered to detect spurious signatures resulting from BGS. To determine whether our above analyses may have underestimated the rate at which false sweep signatures appear, we adopted a sliding window approach using the large-scale simulations described in the previous section. Specifically, we moved `S/HIC` across these larger simulated chromosomes with small step sizes, asking whether `S/HIC` mistakes the focal window for a sweep at each step. Using 10 kb step sizes for our human simulations and 1 kb for *Drosophila*, we classified 1100 windows for each replicate with `S/HIC`. Importantly, by using these small step sizes we allow `S/HIC` to examine a number of possible sweep locations within each 100 kb window (or 10 kb window in *Drosophila*). We did not observe a significant increase in false positive rate relative to our examination of the central window alone (10): in our human equilibrium scenario the false positive rate is 6.5%, compared to 5% when examining the central window alone (*P* = 0.69); in the Tennessen European model the false positive rate is 3.6%, compared to 2% when examining the central window (*P* = 0.59); and 0.93% versus 0% in `Drosophila` (*P* = 1.0). (Note that these *P*-values may be anti-conservative due to the autocorrelation of tests of nearby windows within our larger simulated chromosomes; Hahn 2006.). These results imply that our primary approach of simulating larger numbers of small chromosomes experiencing BGS should randomize the location of any spurious sweep-like signatures, such that `S/HIC` should yield an unbiased estimate of their frequency of occurrence. Taken together, our findings suggest that BGS does not appear to systematically mimic hitchhiking even in larger simulated chromosomes.

### 4.9 BGS increases the false negative rate for selective sweeps

We have shown that realistic models of BGS do not frequently produce sweep-like signatures. However, BGS could also potentially confound scans for hitchhiking events by eroding the signature of positive selection, thereby increasing the false negative rate. To address this possibility, we generated forward simulations with recently completed hitchhiking events (see Methods), and asked whether sweeps occurring in concert with BGS were more difficult to detect than those occurring on an otherwise neutrally evolving background. Because this approach was fairly computationally intensive, we limited our anaysis to a single demographic history: the Tennessen et al. 2012 model of human European demography.

We wished to match the selective parameters of our coalescent simulations, namely the selection coefficient, time since fixation, and the initial selected frequency in the case of soft sweeps, all of which were drawn from uniform distributions in our coalescent simulations. We therefore subsampled our simulations by dividing them into discrete bins based on these parameter values, drawing replicates for our final data set uniformly from these bins (Methods). However, because this binning approach was fairly coarse, it may not perfectly match the uniform distributions. Therefore, we assessed the impact of BGS on `S/HIC`’s false negative rate by comparing classification results between two sets of forward simulations—those including BGS, and those without BGS (Figure 11). Nonetheless, we found that our forward simulations of sweeps without BGS were qualitatively similar to our coalescent simulations in terms of the number of sweeps detected (89.8% and 80.4% of forward- and coalescent-simulated hard sweeps detected, respectively and 71.1% and 79.2% of soft sweeps detected), although the fraction classified as hard or soft differed more substantially between the two simulated datasets.

**Figure 10:**
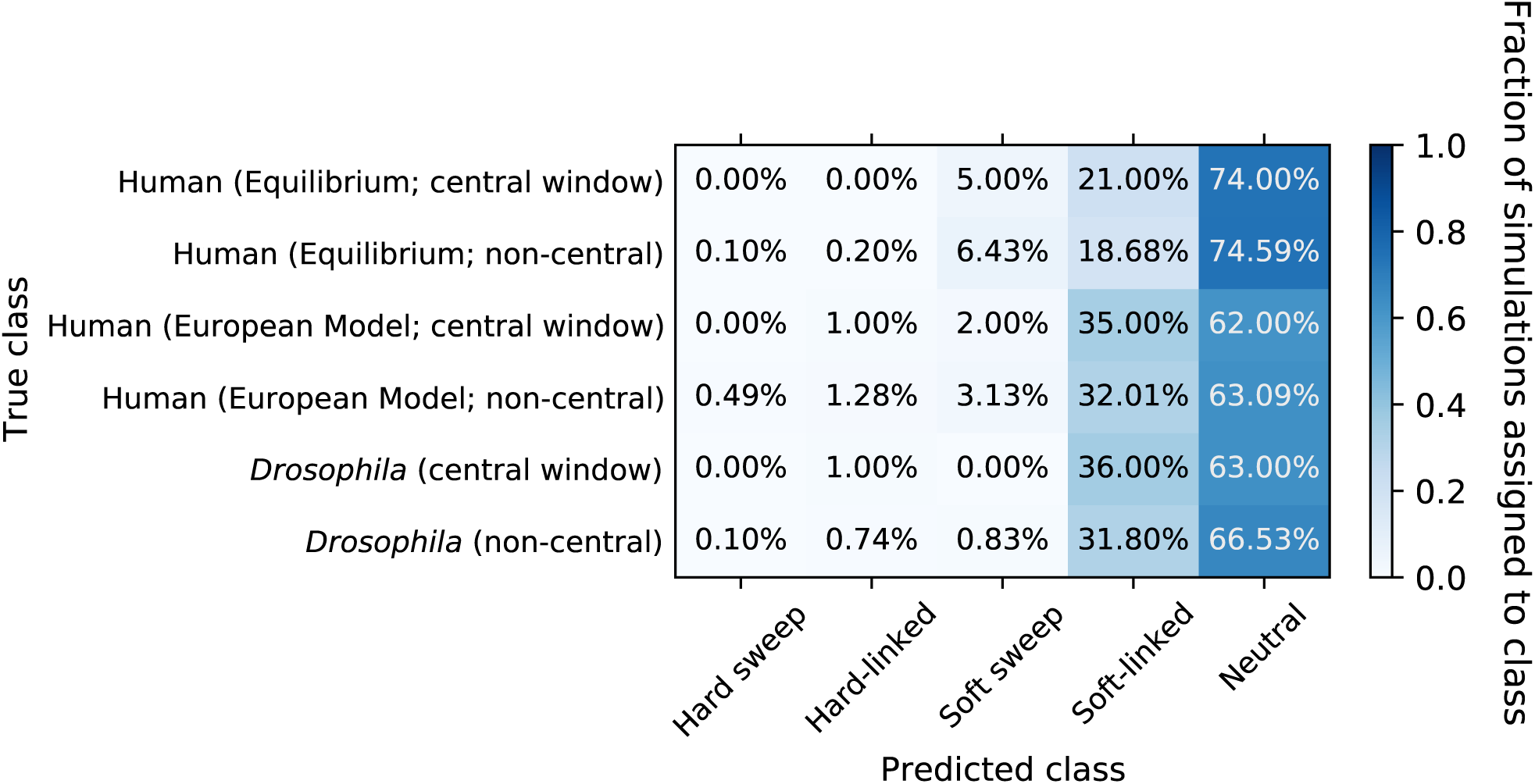
Confusion matrix showing the fraction of windows from larger simulated chromosomes that were assigned to each class by `S/HIC`. Each row shows the results from a set of 100 simulated large chromosomes. Rows marked “central window” show classification results for the central window of the simulated chromosome only, while rows marked “non-central” show results averaged across each of the remaining windows within the chromosome.

**Figure 11:**
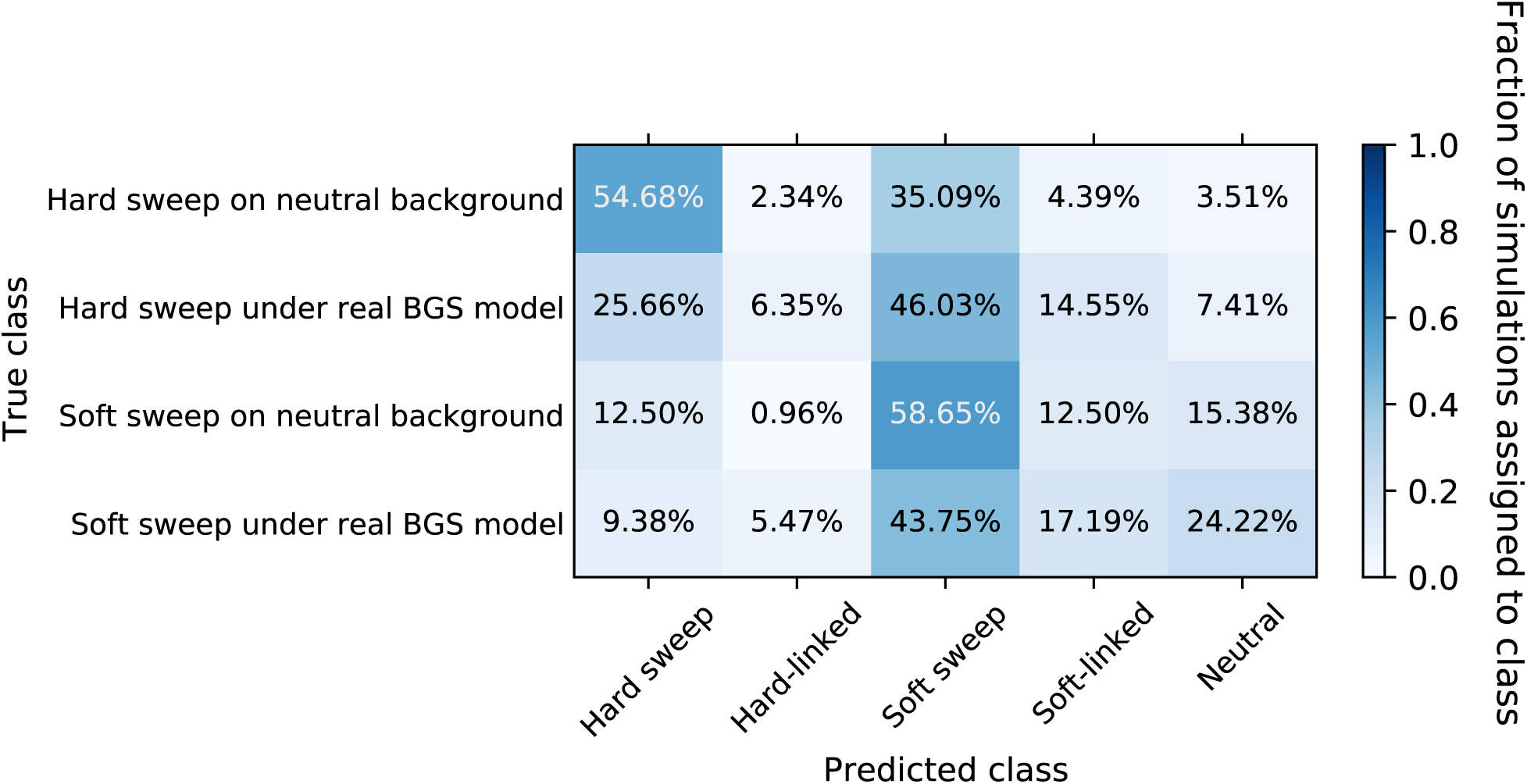
Confusion matrix showing the classification results of forward simulations of recent hard and soft selective sweeps occuring under the Tennessen et al. 2012 model with and without BGS.

We observed a substantial deficit of sweeps detected under BGS versus an otherwise neutrally evolving chromosome. For example, 71.7% of hard sweeps simulated under BGS were classified as a sweep of either type by `S/HIC`, significantly lower than the 89.8% of sweeps without BGS that were recovered (*P* = 6.72 × 10^−10^; Figure 11). Moreover, hard sweeps under BGS were more likely to be misclassified as soft (46.0% versus 35.1%; *P* = 0.0031). Similarly, soft sweeps occurring in the presence of BGS were less likely to be detected than those without BGS (71.1% versus 53.1%; *P* = 0.0066). Together, these results suggest that BGS may dull the signatures of completed selective sweeps.

## 5 Discussion

Natural selection can shape patterns of genomic diversity in many ways. BGS is a prime example of this, as its expected patterns depend on the distribution(s) of fitness effects, the locations of selected sites, and the recombination landscape. Thus this model sufficiently flexible that one should not expect a single common “signature” of BGS. This is the motivation for *B*-maps, which take the DFE, recombination map, and functional DNA element coordinates into account in order to predict the reduction in diversity produced by BGS in a genome for which all of this information has been annotated/estimated. Such maps have been touted as an important “baseline” expectation for patterns of diversity across the genome (Comeron 2014). Unfortunately, such maps are only predictive of the impact of BGS on levels of expected heterozyogisty (i.e. the degree of reduction in *π* produced by BGS). To model the full distribution of genealogies yielded by BGS on the basis of a genome annotation, one can use forward population genetic simulations, which are becoming increasingly computationally efficient (Thornton 2014; Haller *et al*. 2019; Kelleher *et al*. 2018). The present study attempts to do this by simulating large regions designed to mimic the spatial arrangement of functionally important sites across the genomes of humans and *Drosophila*, thereby modeling the effects of both direct and linked negative selection in these genomes. More extensive simulation of chromosome-sized segments under this approach could be used to produce analogs of the *B*-map for any set of summary statistics under an arbitrary demographic history. Such an approach may prove useful as a multi-dimensional baseline expectation of different summaries of diversity under BGS alone.

The goal of this study was to use forward simulation to investigate the expected patterns of diversity created by BGS in humans and *Drosophila*, and compare them to expectations under recent hard and soft selective sweeps. We find that some parameterizations of BGS do indeed yield a valley in diversity similar to that expected under a sweep. These results are consistent with a previous finding that simply taking genomic windows that are outliers with respect to *π* will result in a large number of false positives (Comeron 2014), although this approach is not commonly taken in practice. However, other statistics do not appear to be affected in the same manner as *π* (compare the Central BGS scenario to the hard and soft sweep scenarios in Figure 3 and Supplementary Figures 11 and 22).

Perhaps more importantly, in real genomes few regions have an arrangement of functional elements that coincidentally mirrors those designed with the intention of confounding scans for selection. Thus, if we examine the complete landscape of genetic variation across a chromosome, the impact of BGS on the false positive rate for selective sweeps should be minimal. Instead, our results suggest that the primary impact of BGS on scans for hitchhiking events may instead be an elevated false negative rate. This effect is probably due to a combination of Hill-Robertson interference Hill and Robertson, 1966, and the shrinking of genealogies across the chromosome, including in regions flanking selective sweeps, caused by BGS. Both of these phenomena will cause the spatial skews in patterns of polymorphism produced by hitchhiking events to be less pronounced. The effect of negative selection on selective sweep detection requires further study, and is it is possible that other approaches may be able to detect sweeps in the presence of BGS with greater sensitivity than `S/HIC`. An alternative strategy to detecting sweeps may also be to attempt to discriminate between selective sweeps and BGS, rather than solely considering neutrality as a baseline (Comeron 2014).

In recent years, several methods have been devised to use spatial patterns of multiple summaries of genetic variation around a focal region in order to detect selective sweeps (Lin *et al*. 2011; Schrider and Kern 2016; Mughal and DeGiorgio 2018), and our results suggest that these methods should be robust to realistic scenarios of BGS (consistent with results from Schrider and Kern 2017 and Mughal and DeGiorgio 2018). Indeed, a recent method for detecting hitchhiking using trend-filtered regression appears to be fairly robust even to a BGS scenario concocted to resemble selective sweeps (Mughal and DeGiorgio 2018). Thus, our conclusions about the separability between models of BGS and hitchhiking events could be viewed as conservative. In contrast to BGS, demographic history is likely to be an important confounding factor for detecting natural selection in practice (Simonsen *et al*. 1995; Jensen *et al*. 2005; Nielsen *et al*. 2005). Researchers should thus continue to focus on the development of methods that are robust to non-equilibrium demographic histories, especially in cases where the true history is unknown (Schrider and Kern 2016; Mughal and DeGiorgio 2018)—this is likely to often be the case in practice given that demographic estimates will themselves be biased by the impact of natural selection on polymorphism (Ewing and Jensen 2016; Schrider *et al*. 2016).

We modeled our BGS scenarios after two very different genome architectures: the human genome in which only 5% of sites are found within either coding or conserved non-coding elements, and the *Drosophila melanogaster* genome in which a majority of sites are under direct purifying selection. Thus, we can ask whether genome structure appears to affect the degree to which BGS mimics selective sweeps. In both our human and *Drosophila* simulations we see that in the presence of purifying selection and BGS the majority of genomic regions would not be expected to produce patterns of diversity consistent with a selective sweep. This is evidenced by the fact that we observe no elevation in our human simulations in the rate at which regions with BGS are misclassified as sweeps by `S/HIC` relative to neutrally evolving regions, and that although there is an elevation in this rate in *Drosophila*, it is quite subtle (≤ 1%) and limited to low-recombining regions as discussed below. This suggests that in both gene-dense and gene-poor genomes the “gene oasis” scenario modeled in our Central BGS simulations is relatively rare. However, we do note that in both our human- and *Drosophila*-based simulations we found that `S/HIC` classifies regions as soft-linked at an increased rate; this may imply that unmodeled sources of heterogeneity of patterns of diversity can make it more difficult to discriminate between neutral evolution and linkage to nearby sweeps. Indeed, we previously observed a similar bias for `S/HIC` in the case of demographic misspecification (Schrider and Kern 2016).

Our study has some important limitations in that we only examined two different DFEs (Boyko *et al*. 2008; Huber *et al*. 2017), four fixed dominance coefficients, and three different demographic models that contain population size changes but no migration. There are infinite possible DFEs, distributions of dominance values, and demographic histories, so we cannot rule out the possibility that our results could change qualitatively under particular models and parameterizations. However under each of the three different combinations of demographic history, DFE, dominance, and genome annotation examined here there is no evidence that BGS systematically resembles selective sweeps substantially more often than purely neutral models do. Thus we expect that our conclusions will hold in most genomes where the layout of selected sites does not frequently resemble that of our Central BGS scenario. Our results do suggest that low-recombining regions with a high density of selected sites flanked by primarily nonfunctional DNA may be somewhat more likely to be misclassified as sweeps and thus should be treated with greater caution (Table 1), although our classification results imply that such confounding examples are uncommon. Indeed, it is worth stressing that the elevated false positive rate under BGS in *Drosophila* seems to be confined to regions with little to no recombination, implying that such regions should perhaps be omitted from sweep scans based on spatial patterns of variation—this is a logical step given that such scans search for signatures produced by the interplay between selection and recombination, and if the latter is absent there is no reason to expect such a signature. Another type of region that may be problematic is that where the recombination rate is low but only in the central portion of the window to be classified. Although we did not examine this possibility here, Mughal et al. (2018) previously showed that dramatically decreasing the recombination rate only in the central portion of a region while keeping the locations of negatively selected sites constant produces a modest increase in the false positive rate.

Our analysis also focused primarily on relatively small simulated chromosomal regions, although we did simulate a number of larger chromosomes and found no evidence that they produce spurious BGS signatures at a higher rate. This result is intuitive because, although larger chromosomes result in more linked selection influencing a given focal window, there is no reason to expect that it would produce a valley of diversity near the center of this window or create other spatial signatures of a sweep—indeed, including more distant flanking selected sites should reduce diversity more on the edge of the focal window than in its center.

It is also important to note that because our strategy was to base our simulations entirely on empirical genome annotations and DFEs, we are limited to considering the effect of single nucleotide mutations. It appears to be the case that the DFE in non-coding regions is skewed toward weaker selection coefficients (Racimo and Schraiber 2014), and this is a feature of our Real BGS–Weak CNE model. However, our simulations ignore additional mutation types such as indels and transposable element insertions and other structural variants (SVs) that are probably skewed toward stronger selection coefficients. In the standard Real BGS model we have the same DFE for both coding and conserved non-coding DNA, and thus this model produces more strongly deleterious mutations than the Weak CNE model. The fact that both models produced very similar results for all three species could suggest that increases to the rates and fitness effects of deleterious mutations may not cause BGS to more closely resemble sweeps. However, the impact of indels and SVs warrants further investigation, and future efforts should incorporate both the rates and DFEs of additional mutation types once they are known more precisely.

We have also only considered scans for recent completed selective sweeps. Thus we have not examined other selection scenarios such as balancing selection or partial selective sweeps, though we have no reason to believe that BGS will systemically mirror either of these selective scenarios, with the exception of very low-frequency partial sweeps which may be indistinguishable from drifting deleterious mutations (Maruyama 1974). We also note that a recent study examining local adaptation in populations with gene flow concluded that BGS is unlikely to increase the fraction of false positives produced by scans for *F*_*ST*_ outliers (Matthey-Doret and Whitlock 2019). However, scans for other selective scenarios, including much older selective sweeps where the signature may have degraded considerably (Schrider *et al*. 2015), or polygenic selection which in some scenarios are expected to produce more subtle shifts in allele frequencies (Jain and Stephan 2017; HÖllinger *et al*. 2019; Thornton *2019*), may have different propensities to be mistaken for BGS than the hitchhiking models examined here.

Although we show that BGS effects the mean values of several population genetic summary statistics, for most of the statistics we examined it does not create spatial patterns qualitatively similar to those expected under hitchhiking. Thus, our results demonstrate that efforts to detect recent positive selection should utilize the broader genomic spatial context of high-dimensional summaries of variation. Importantly, sweep-detection methods that use this information (Lin *et al*. 2011; Schrider and Kern 2016; Mughal and DeGiorgio 2018) rather than relying on univariate summaries or examining a narrowly defined genomic region can readily detect sweeps in the presence of purifying selection and BGS. Moreover, our findings imply that attempts to disentangle the relative effects of hitchhiking and BGS on levels of diversity genome-wide (Elyashiv *et al*. 2016; Booker and Keightley 2018) could be made even more effective by incorporating additional summaries of variation, although this may necessitate a reliance on simulated data rather than the use of likelihood estimation. Such efforts could also help to answer the question of to what extent hitchhiking and BGS are responsible for the limited range of neutral diversity observed across species (Lewontin 1974; Leffler *et al*. 2013; Corbett-Detig *et al*. 2015; Coop 2016).

## 6 Acknowledgments

The author thanks Matt Hahn and Andy Kern for feedback on the manuscript, and Kevin Thornton for help with `fwdpy11`. This work was funded by the National Institutes of Health under award number R00HG008696.

---

**Supplementary Figure 1:**
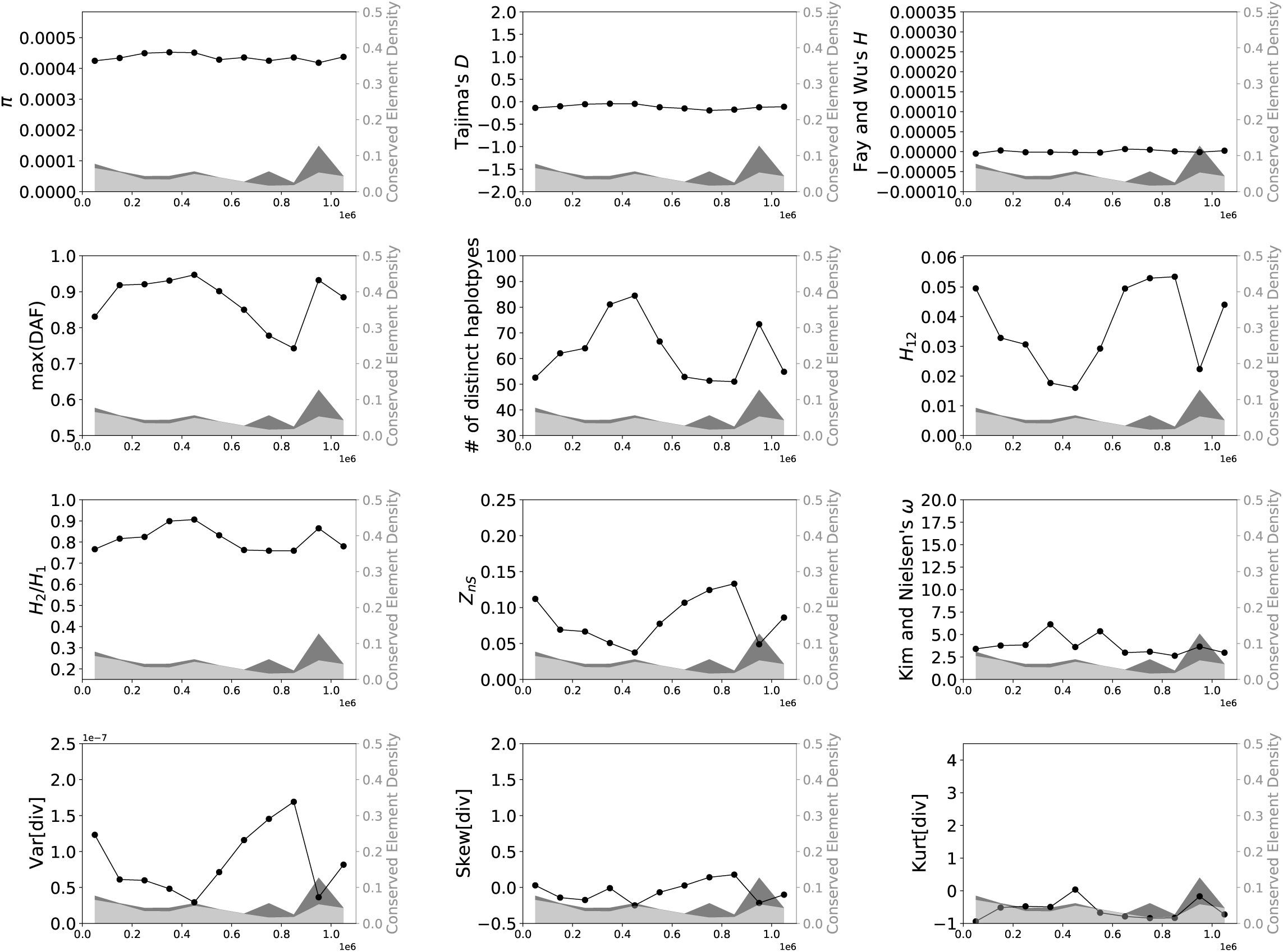
Values of 15 statistics calculated from a simulated data set of 1000 replicates designed to match the locations of conserved sites from chr7:37000001-38100000 in the human genome (hg19 assembly version), simulated under equilibrium demography. Each replicate used the same overall recombination rate and map (chosen to match this region), however the mutation rate varied from replicate to replicate in the same manner as described in the Methods. Position labels on the *x*-axes are relative to the beginning of the window. The density of exonic and conserved non-coding DNA is shown in the shaded dark gray and light gray regions, respectively.

**Supplementary Figure 2:**
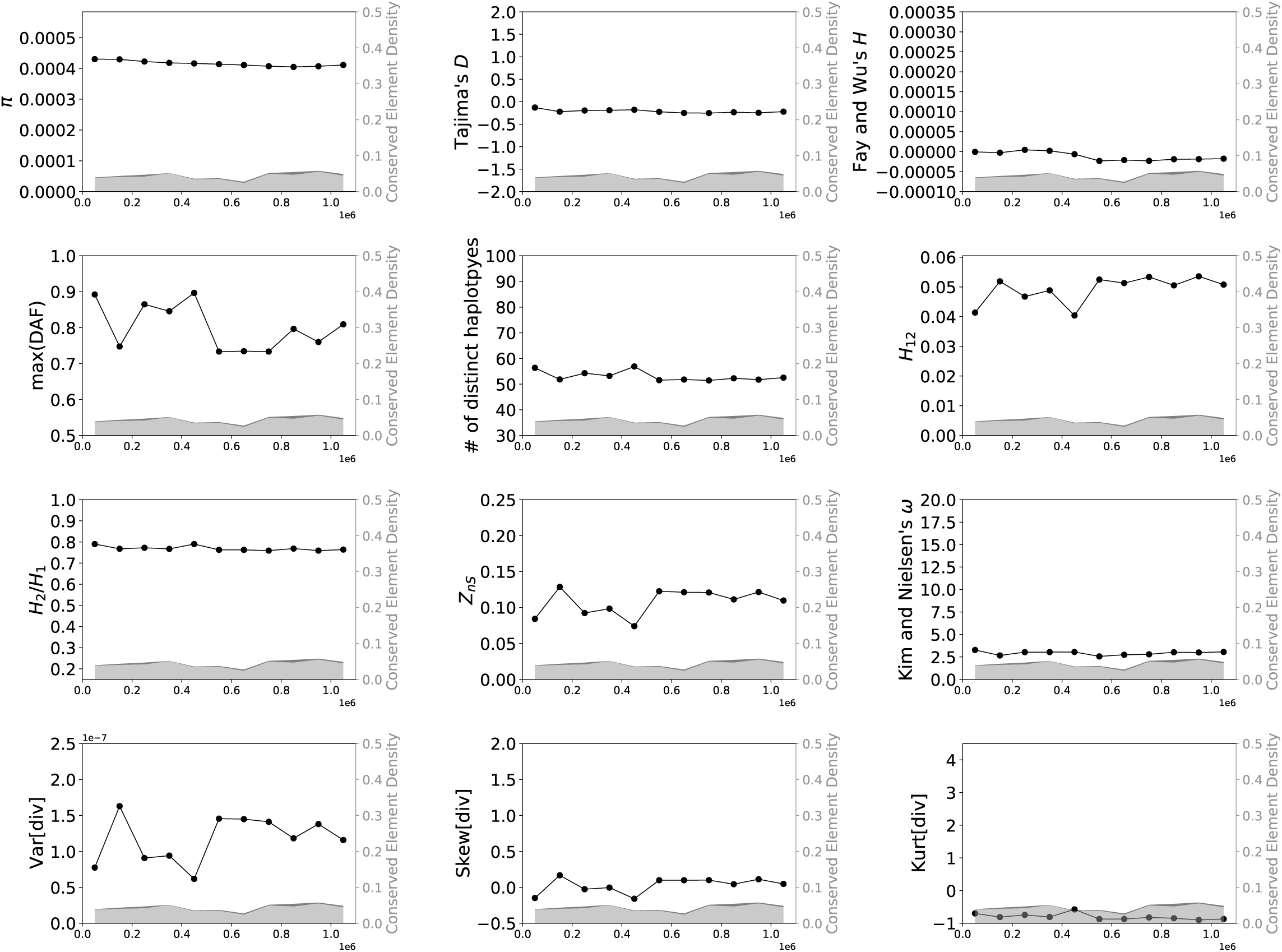
Values of 15 statistics calculated from a simulated data set designed to match the locations of conserved sites from chr4:98200001–99300000 in the human genome (hg19 assembly version), simulated under equilibrium demography. See legend of Supplementary Figure 1 for more detail.

**Supplementary Figure 3:**
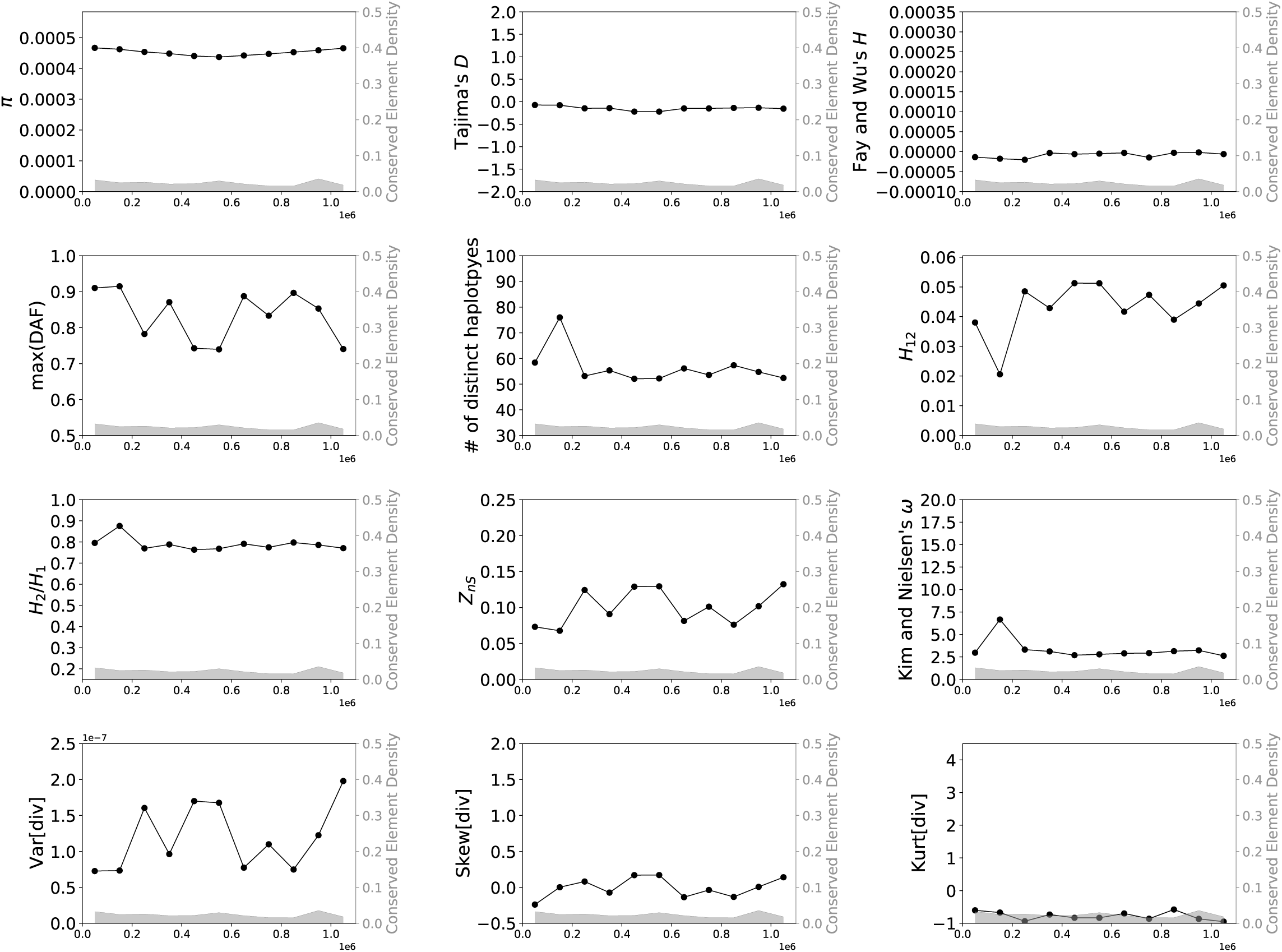
Values of 15 statistics calculated from a simulated data set designed to match the locations of conserved sites from chr4:131500001–132600000 in the human genome (hg19 assembly version), simulated under equilibrium demography. See legend of Supplementary Figure 1 for more detail.

**Supplementary Figure 4:**
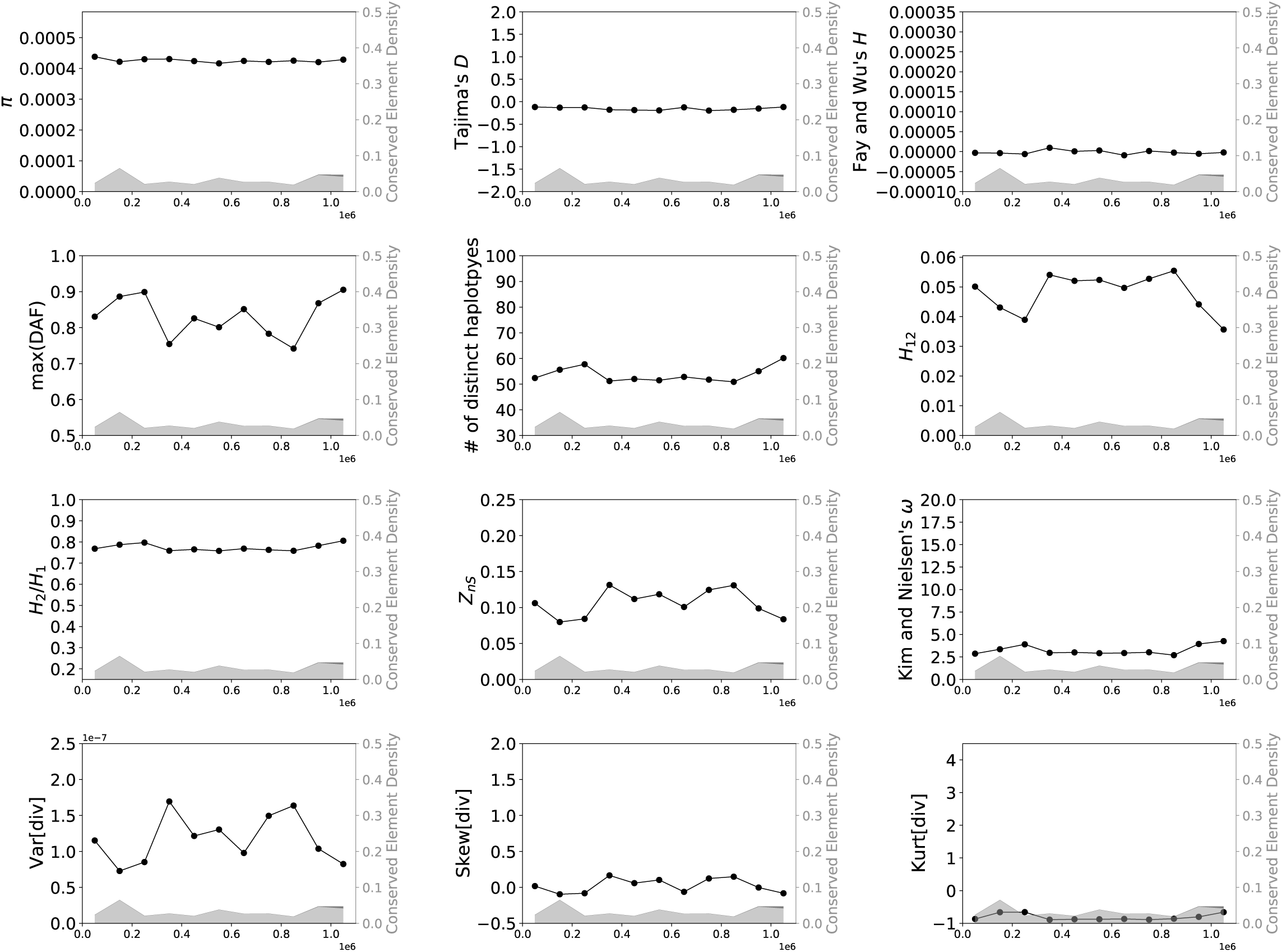
Values of 15 statistics calculated from a simulated data set designed to match the locations of conserved sites from chr3:80100001–81200000 in the human genome (hg19 assembly version), simulated under equilibrium demography. See legend of Supplementary Figure 1 for more detail.

**Supplementary Figure 5:**
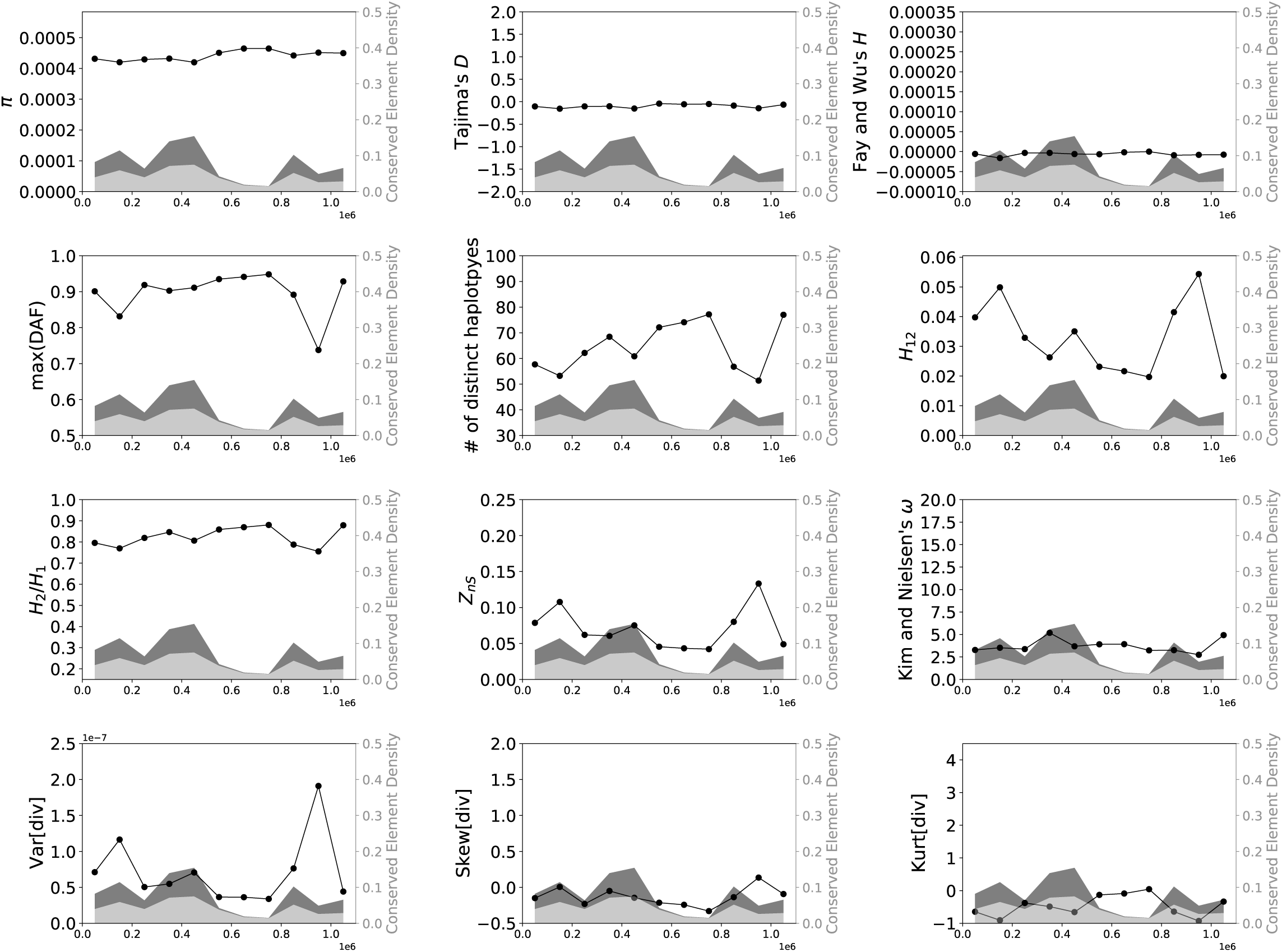
Values of 15 statistics calculated from a simulated data set designed to match the locations of conserved sites from chr3:12500001–13600000 in the human genome (hg19 assembly version), simulated under equilibrium demography. See legend of Supplementary Figure 1 for more detail.

**Supplementary Figure 6:**
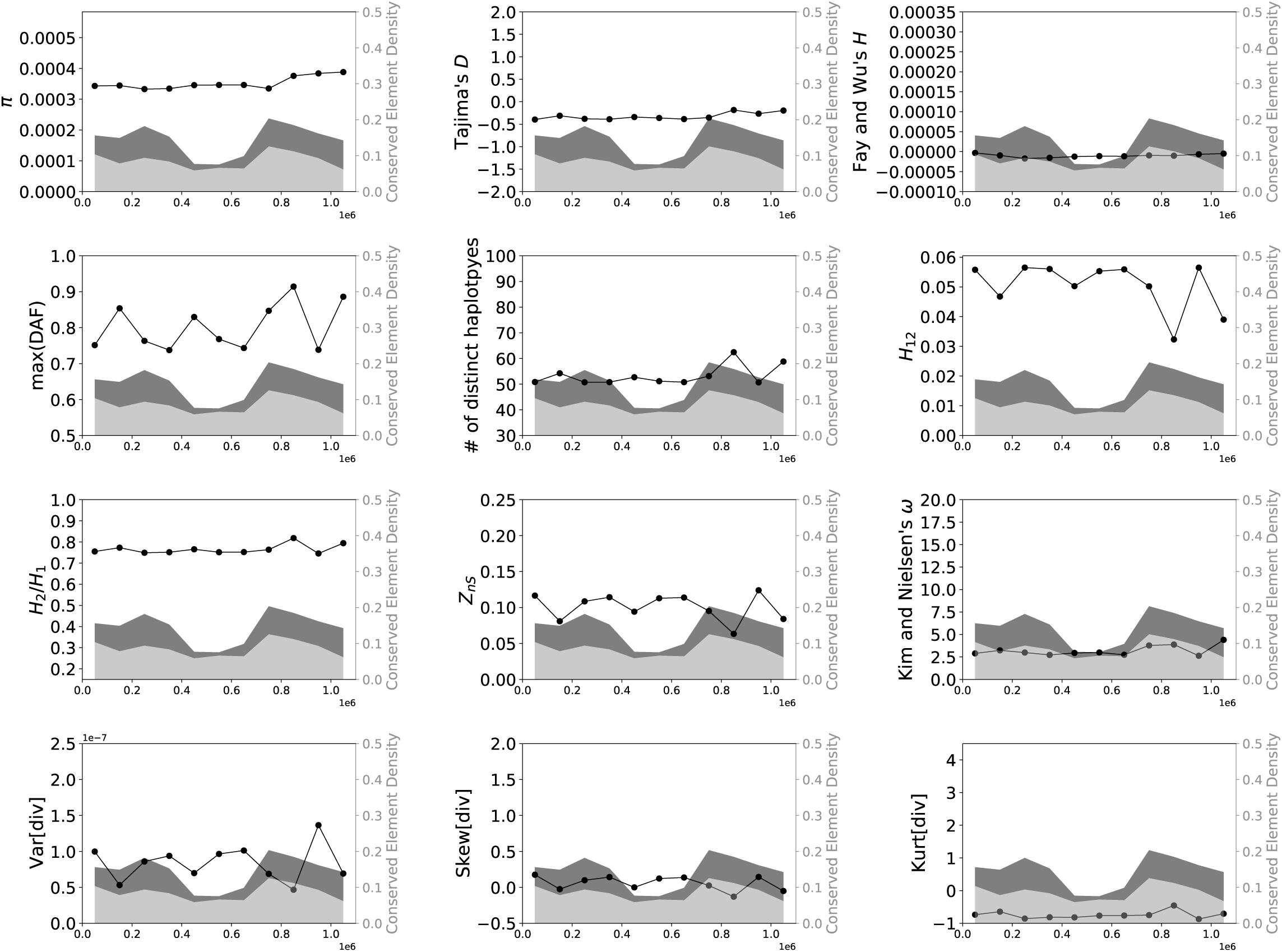
Values of 15 statistics calculated from a simulated data set designed to match the locations of conserved sites from chr1:45800001–46900000 in the human genome (hg19 assembly version), simulated under equilibrium demography. See legend of Supplementary Figure 1 for more detail.

**Supplementary Figure 7:**
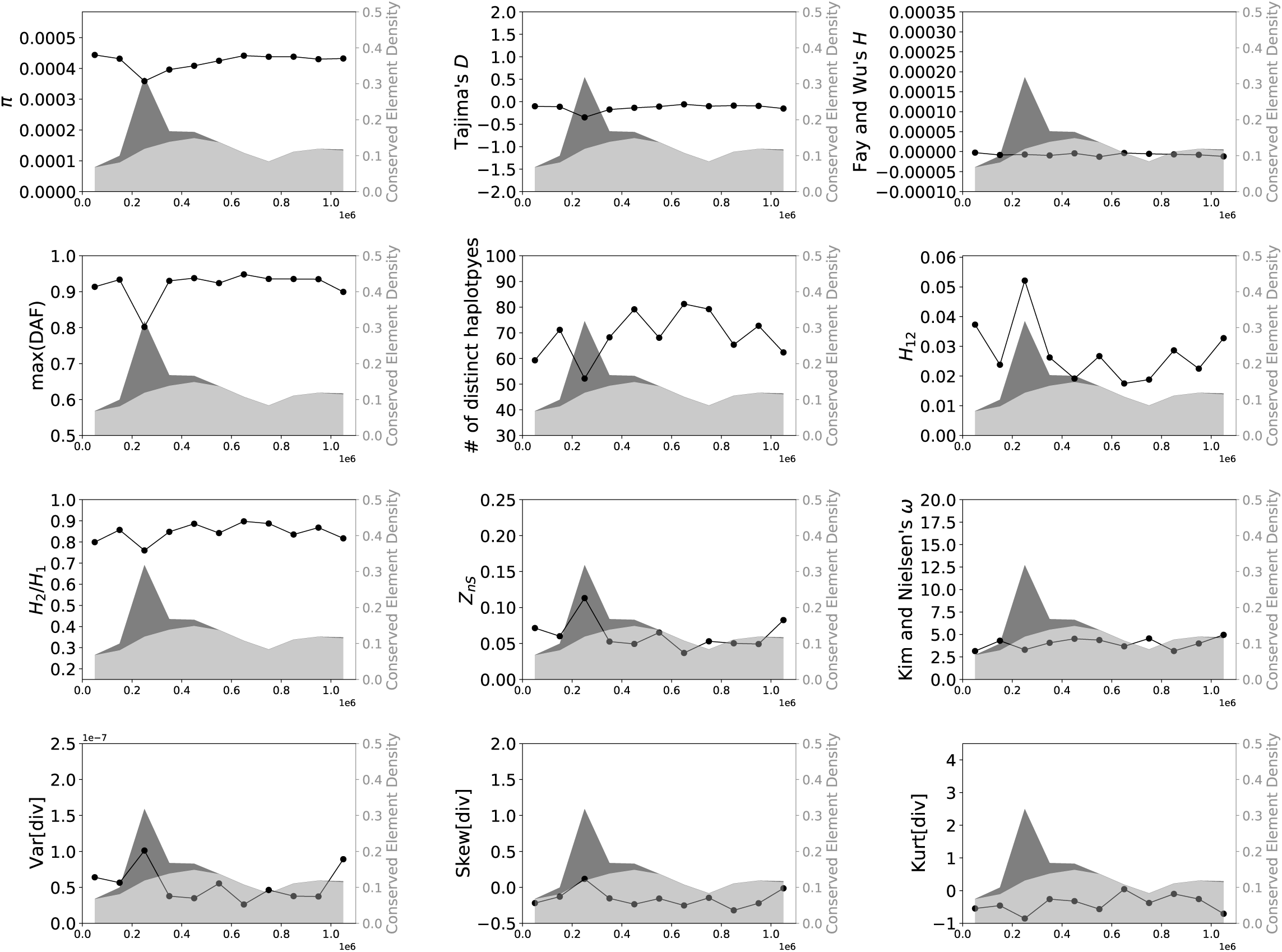
Values of 15 statistics calculated from a simulated data set designed to match the locations of conserved sites from chr15:60500001–61600000 in the human genome (hg19 assembly version), simulated under equilibrium demography. See legend of Supplementary Figure 1 for more detail.

**Supplementary Figure 8:**
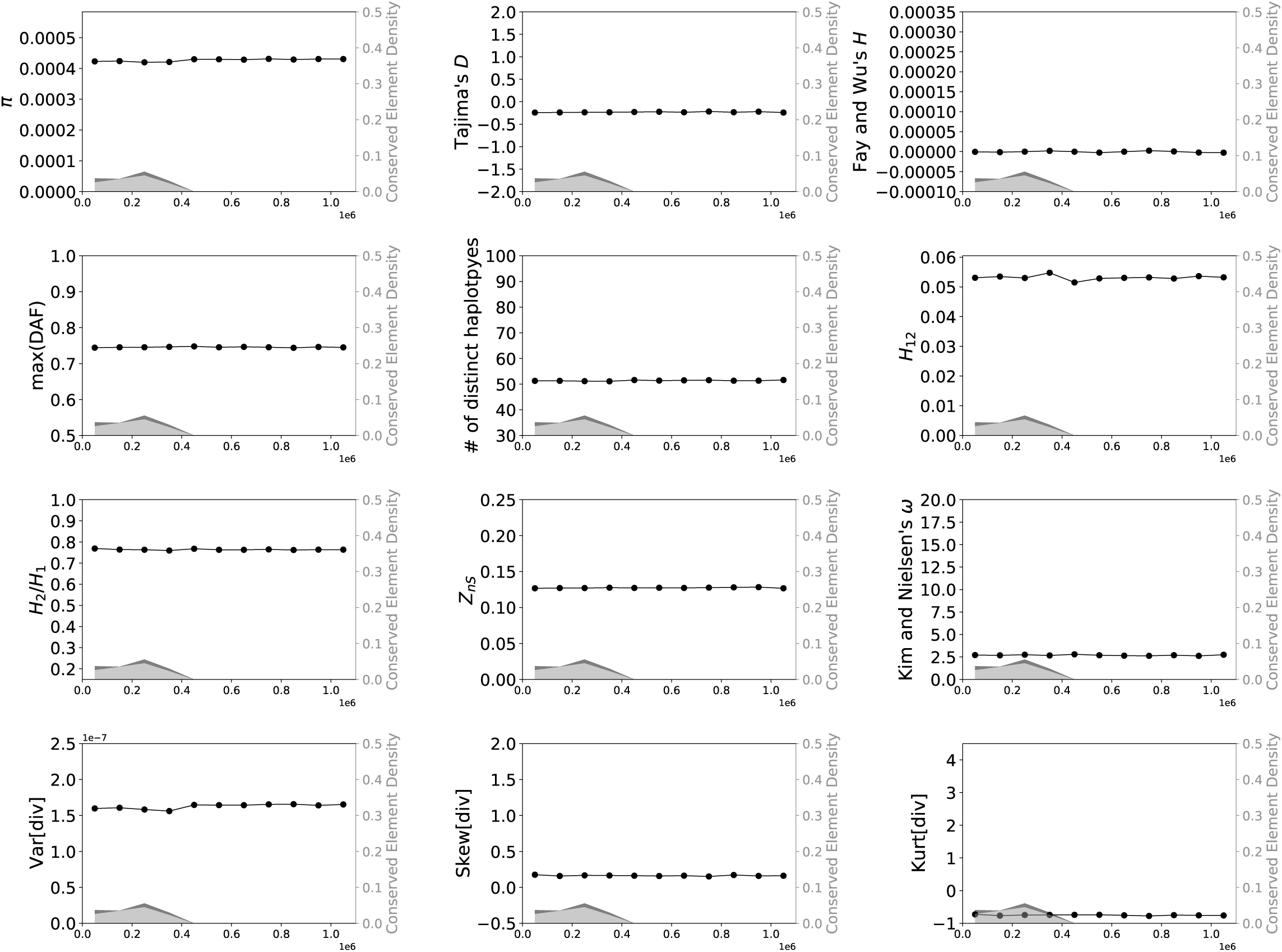
Values of 15 statistics calculated from a simulated data set designed to match the locations of conserved sites from chr11:50000001–51100000 in the human genome (hg19 assembly version), simulated under equilibrium demography. See legend of Supplementary Figure 1 for more detail.

**Supplementary Figure 9:**
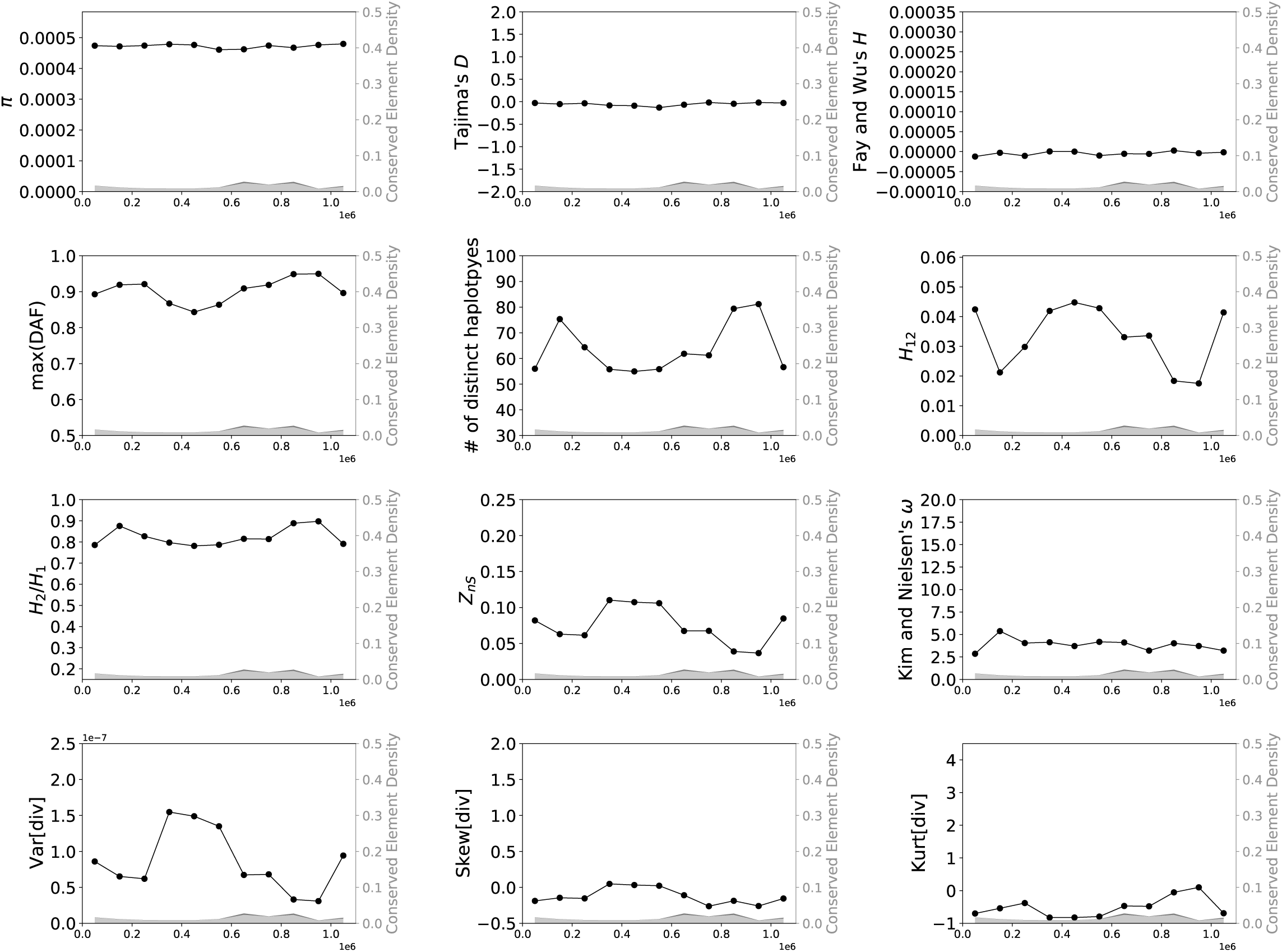
Values of 15 statistics calculated from a simulated data set designed to match the locations of conserved sites from chr11:23900001–25000000 in the human genome (hg19 assembly version), simulated under equilibrium demography. See legend of Supplementary Figure 1 for more detail.

**Supplementary Figure 10:**
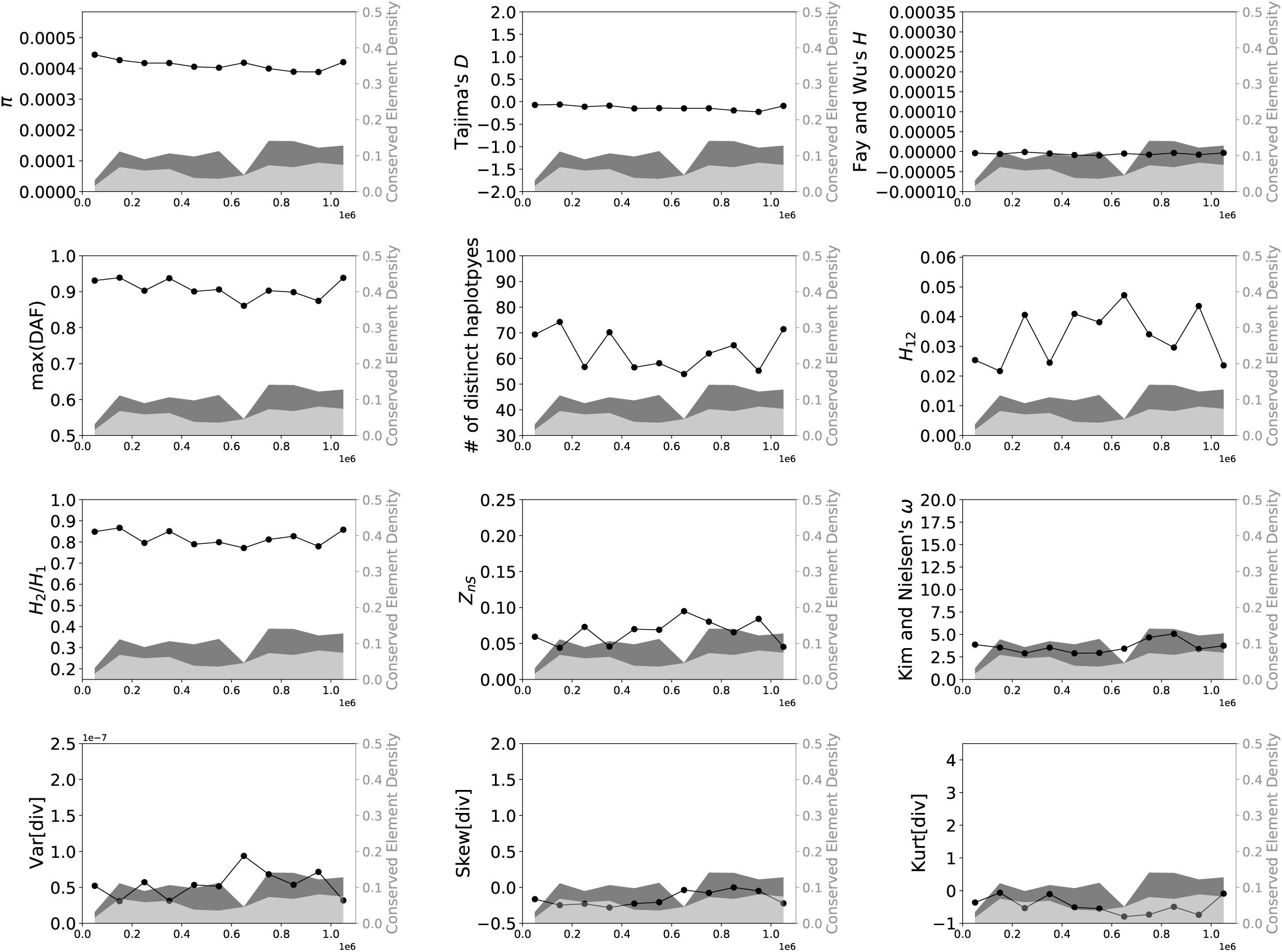
Values of 15 statistics calculated from a simulated data set designed to match the locations of conserved sites from chr10:49800001–50900000 in the human genome (hg19 assembly version), simulated under equilibrium demography. See legend of Supplementary Figure 1 for more detail.

**Supplementary Figure 11:**
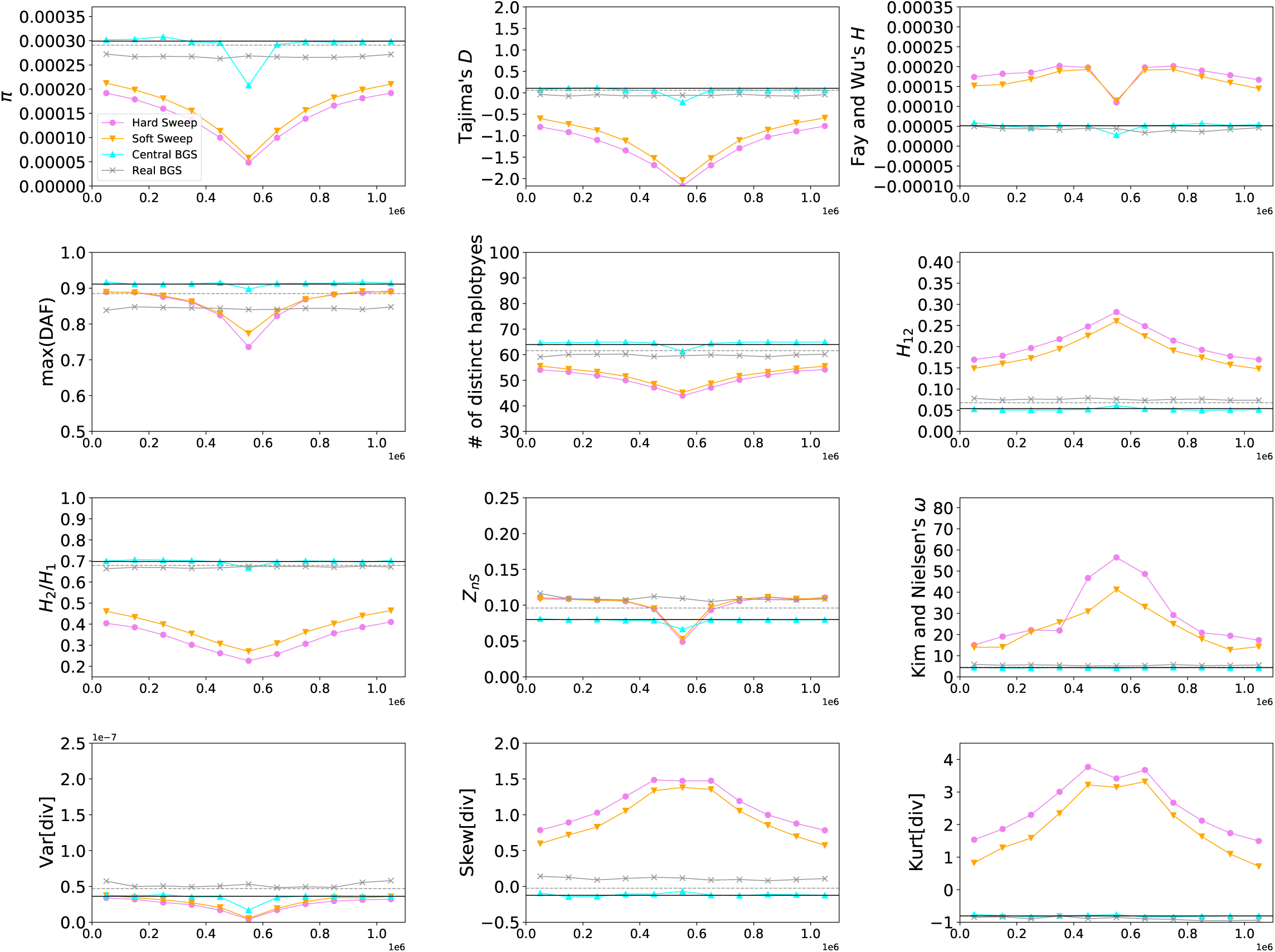
Values of 15 statistics calculated in each of the four data sets simulated under Tennessen et al.’s model of European demography (Tennessen *et al*. 2012). Here, the mean value of each statistic across each simulated replicate is shown in the 11 adjacent 100 kb windows within the 1.1 Mb chromosomal region. In each panel the neutral expectation of the statistic obtained from forward simulations are shown as a horizontal black line, while the neutral expectation from coalescent simulations is shown as a dashed gray line—although for most statistics these two expectations were essentially identical, for some there was a slight discrepancy between the two, perhaps reflecting the subtle differences in the models used by the two simulators.

**Supplementary Figure 12:**
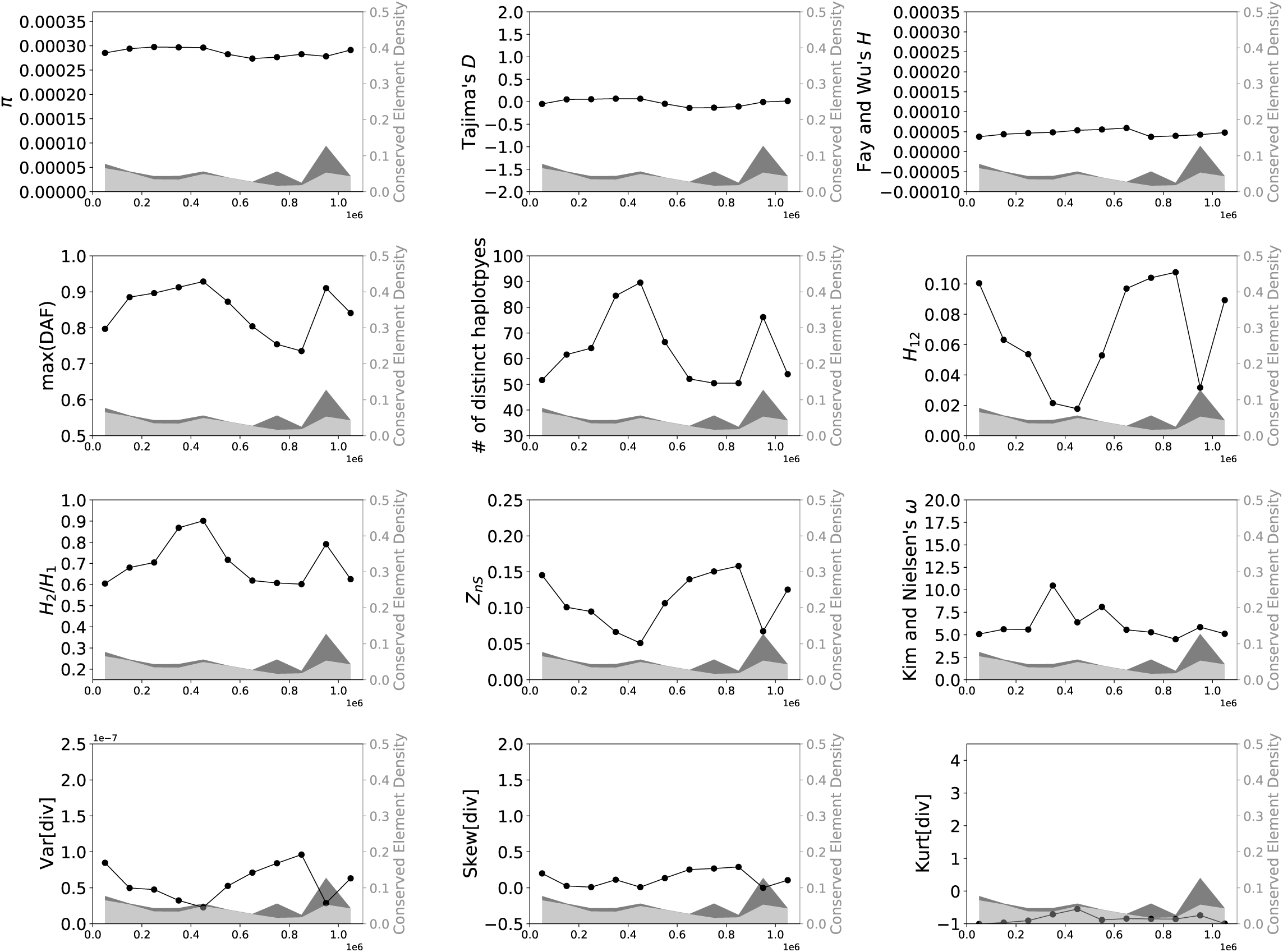
Values of 15 statistics calculated from a simulated data set designed to match the locations of conserved sites from chr7:37000001–38100000 in the human genome (hg19 assembly version), simulated under European population size history (Tennessen *et al*. 2012). See legend of Supplementary Figure 1 for more detail.

**Supplementary Figure 13:**
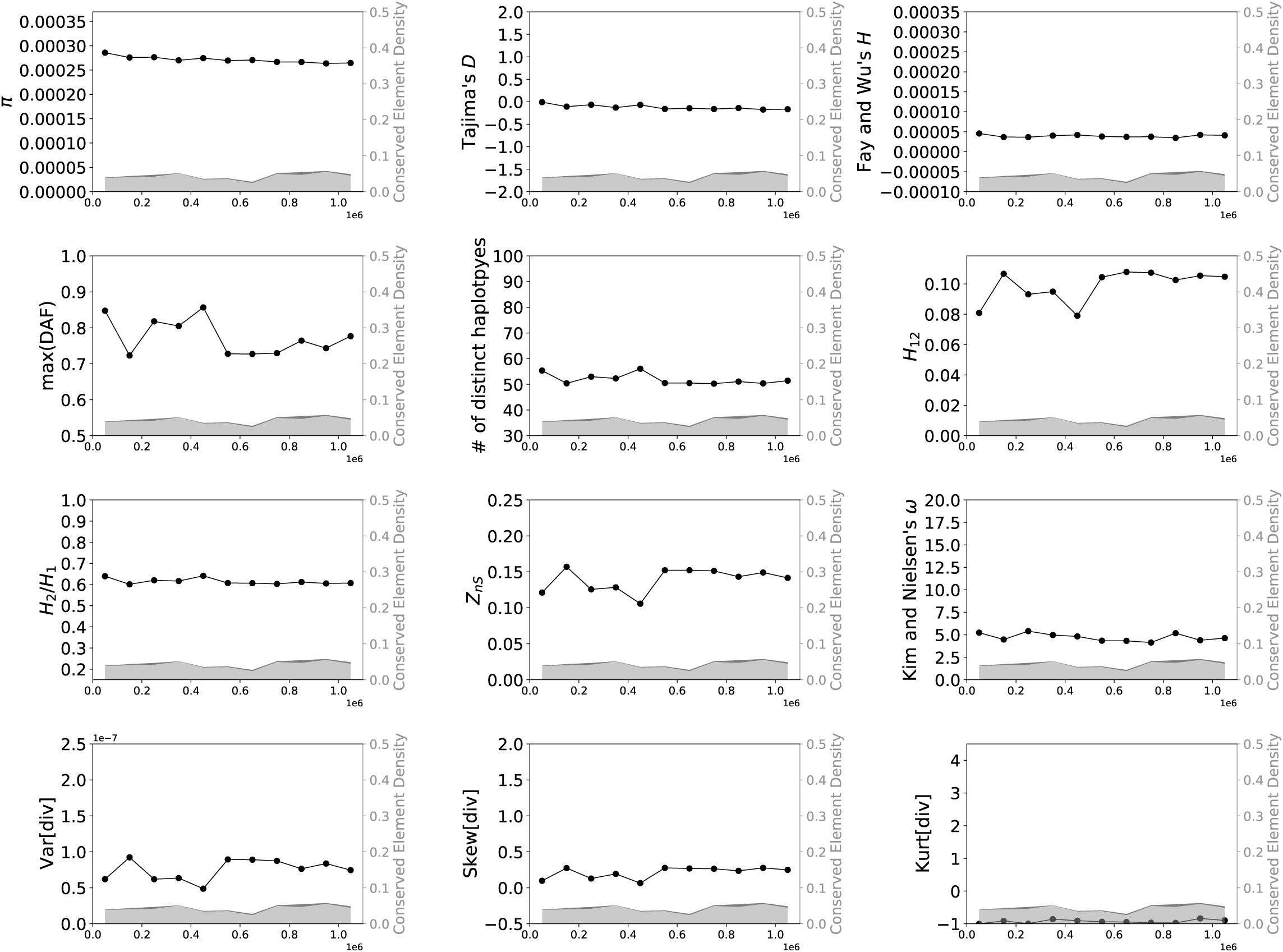
Values of 15 statistics calculated from a simulated data set designed to match the locations of conserved sites from chr4:98200001–99300000 in the human genome (hg19 assembly version), simulated under European population size history (Tennessen *et al*. 2012). See legend of Supplementary Figure 1 for more detail.

**Supplementary Figure 14:**
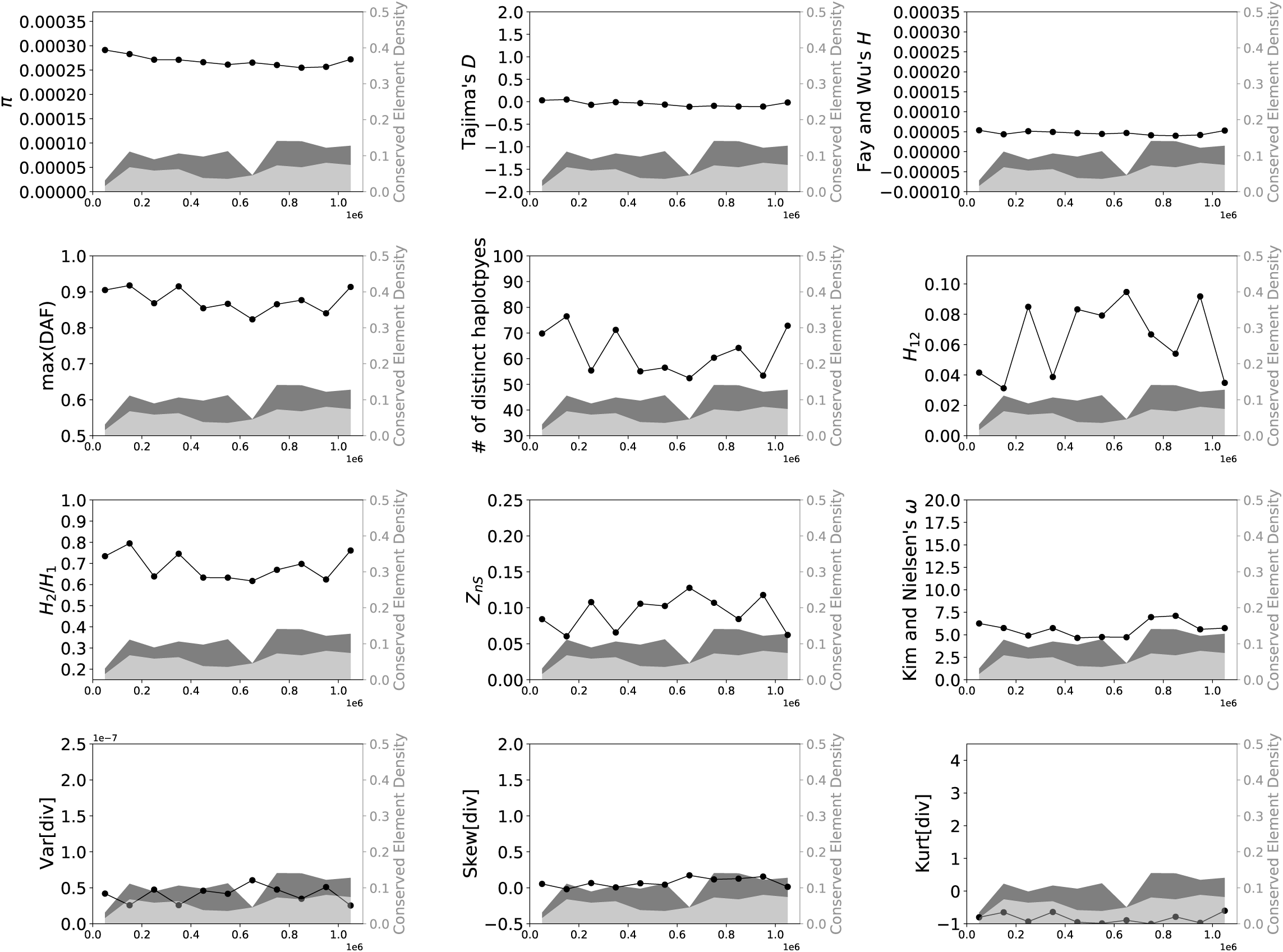
Values of 15 statistics calculated from a simulated data set designed to match the locations of conserved sites from chr10:49800001–50900000 in the human genome (hg19 assembly version), simulated under European population size history (Tennessen *et al*. 2012). See legend of Supplementary Figure 1 for more detail.

**Supplementary Figure 15:**
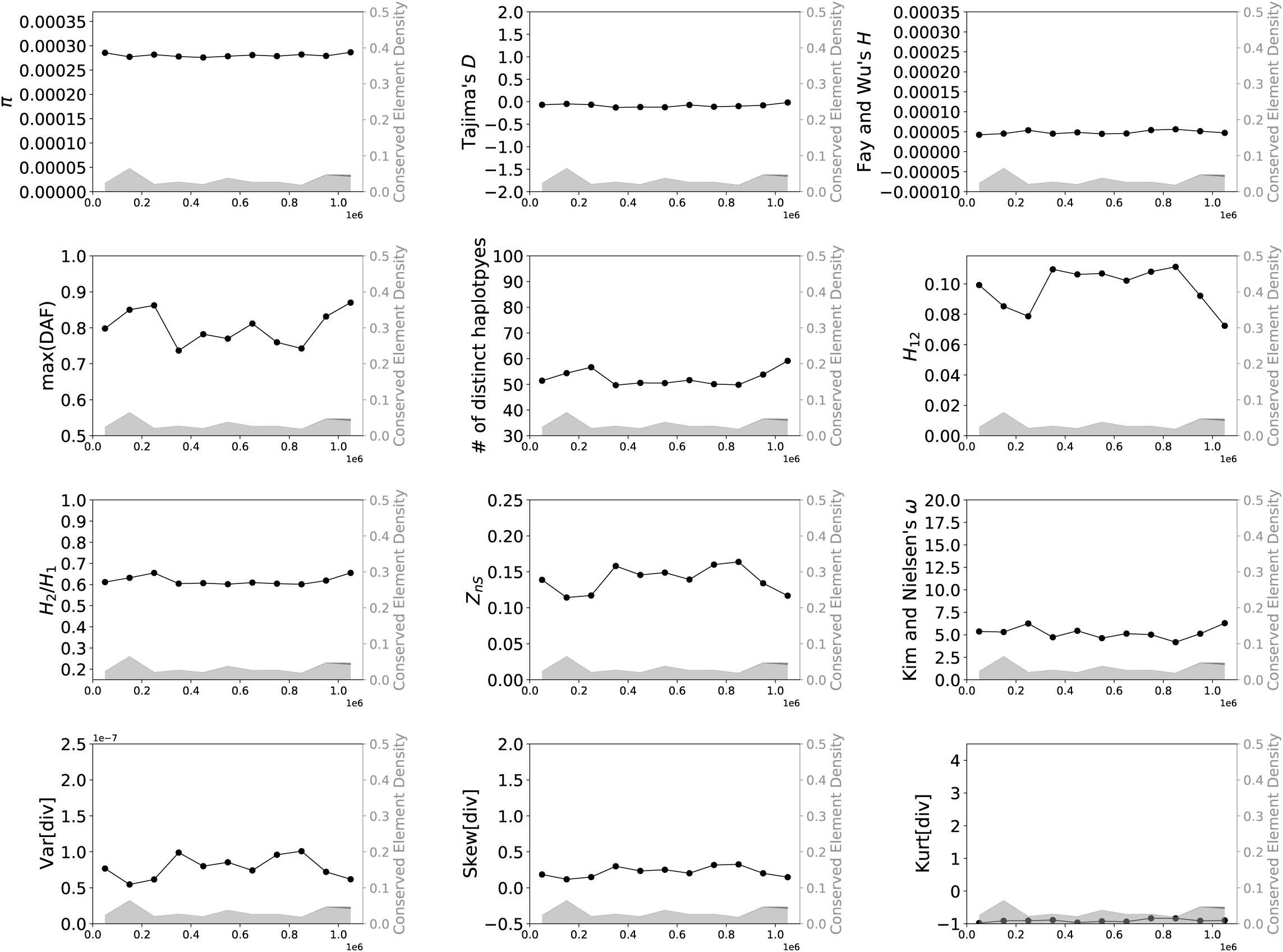
Values of 15 statistics calculated from a simulated data set designed to match the locations of conserved sites from chr3:80100001–81200000 in the human genome (hg19 assembly version), simulated under European population size history (Tennessen *et al*. 2012). See legend of Supplementary Figure 1 for more detail.

**Supplementary Figure 16:**
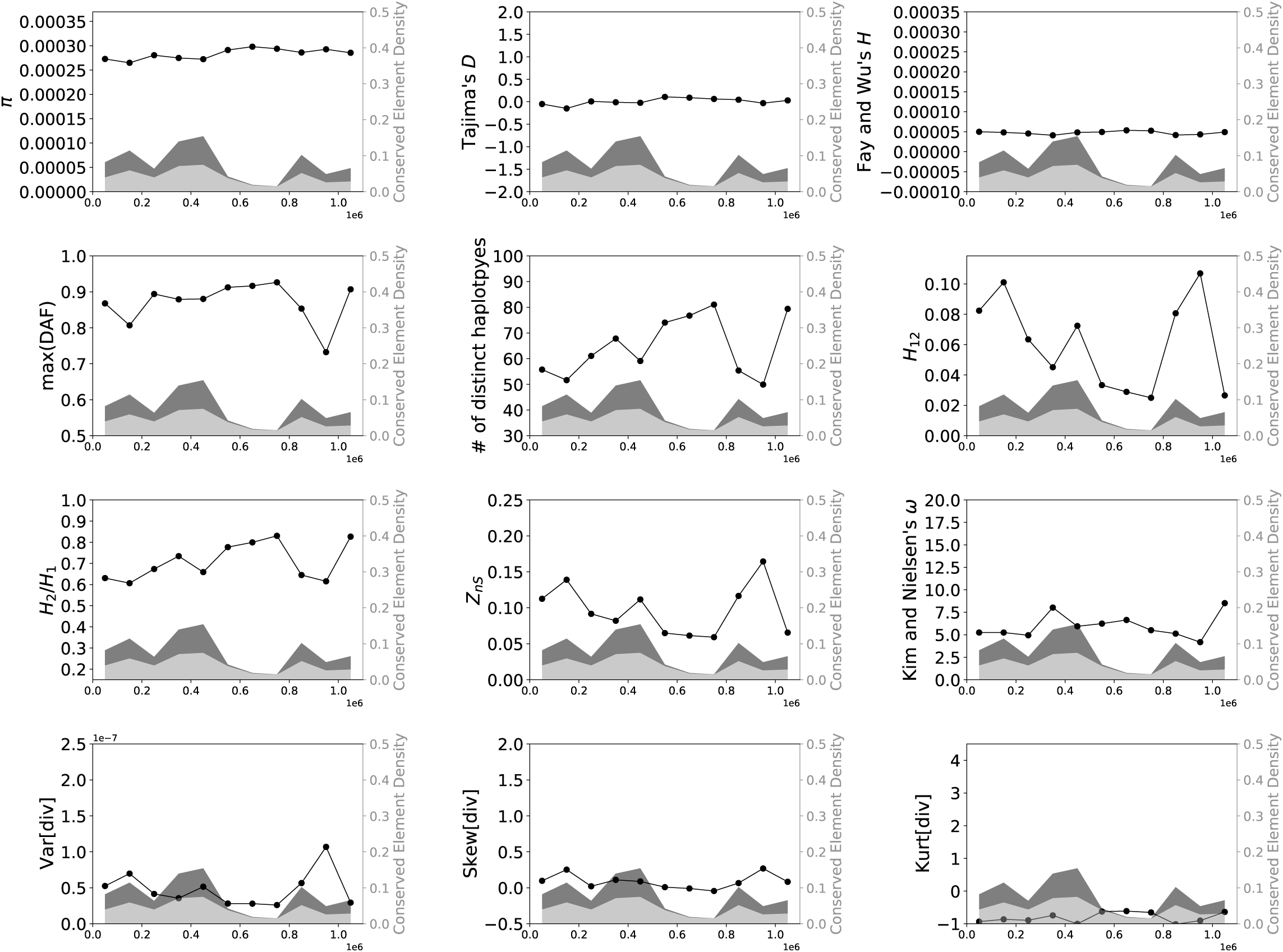
Values of 15 statistics calculated from a simulated data set designed to match the locations of conserved sites from chr3:12500001–13600000 in the human genome (hg19 assembly version), simulated under European population size history (Tennessen *et al*. 2012). See legend of Supplementary Figure 1 for more detail.

**Supplementary Figure 17:**
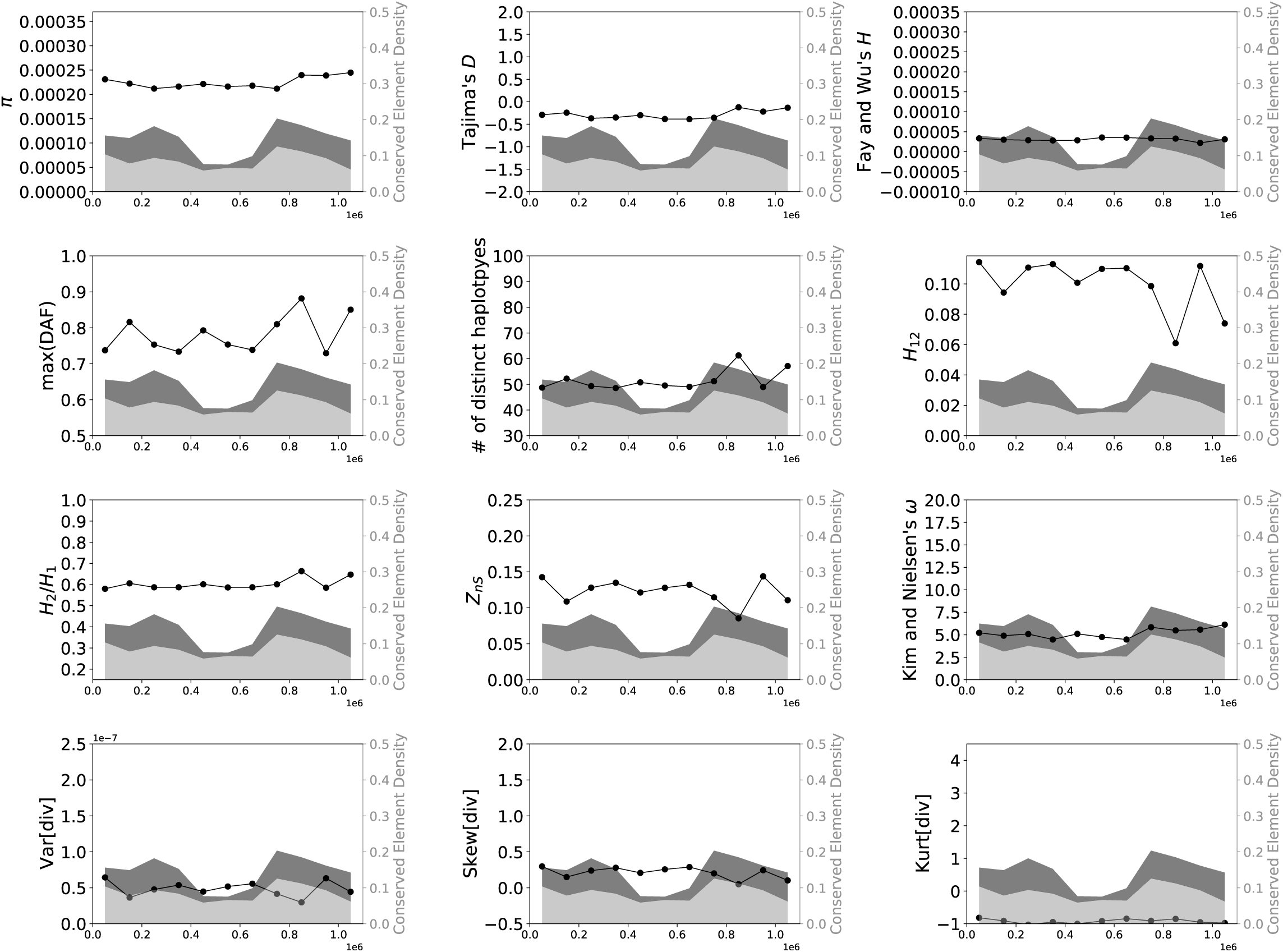
Values of 15 statistics calculated from a simulated data set designed to match the locations of conserved sites from chr1:45800001–46900000 in the human genome (hg19 assembly version), simulated under European population size history (Tennessen *et al*. 2012). See legend of Supplementary Figure 1 for more detail.

**Supplementary Figure 18:**
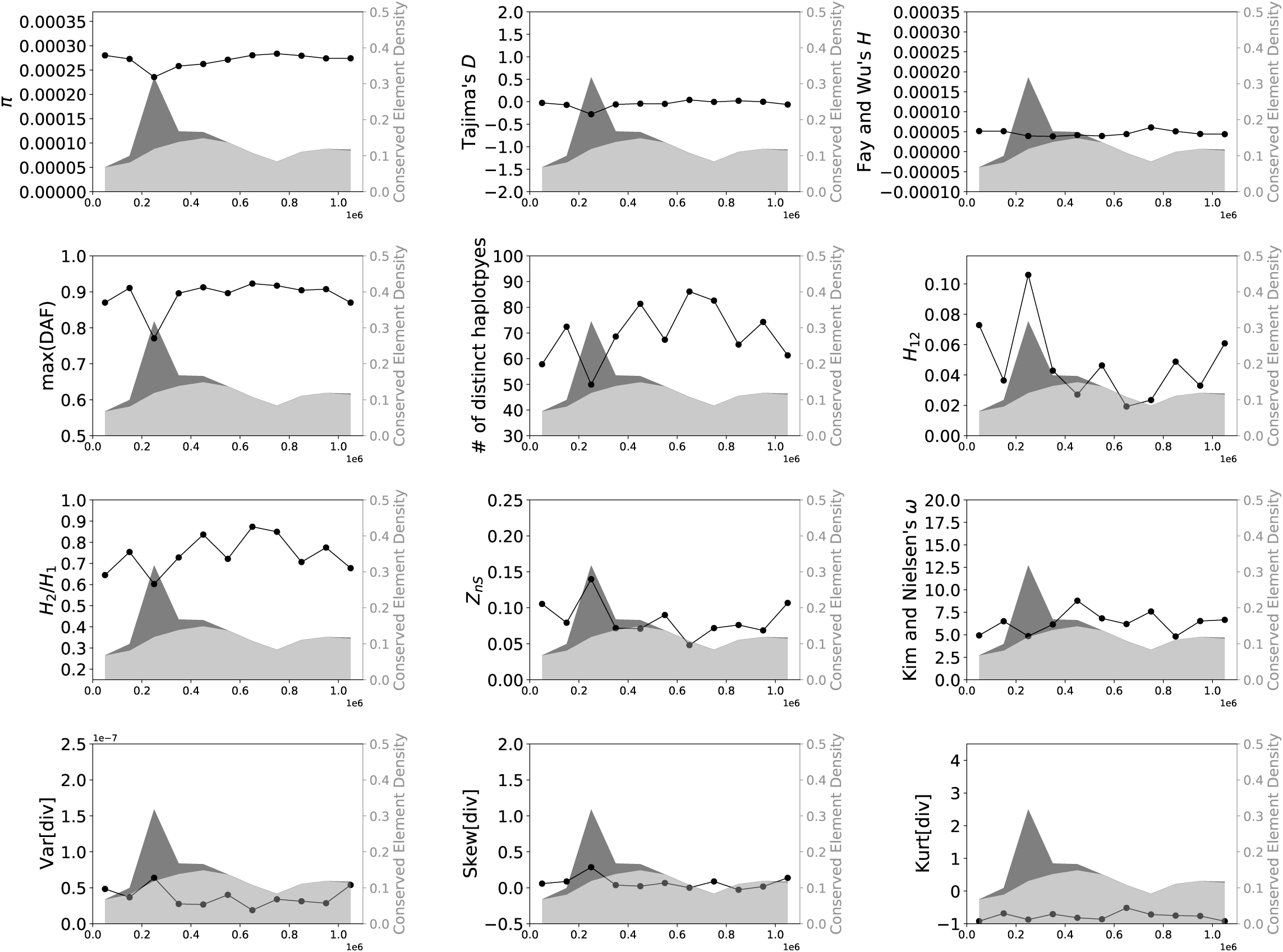
Values of 15 statistics calculated from a simulated data set designed to match the locations of conserved sites from chr15:60500001–61600000 in the human genome (hg19 assembly version), simulated under European population size history (Tennessen *et al*. 2012). See legend of Supplementary Figure 1 for more detail.

**Supplementary Figure 19:**
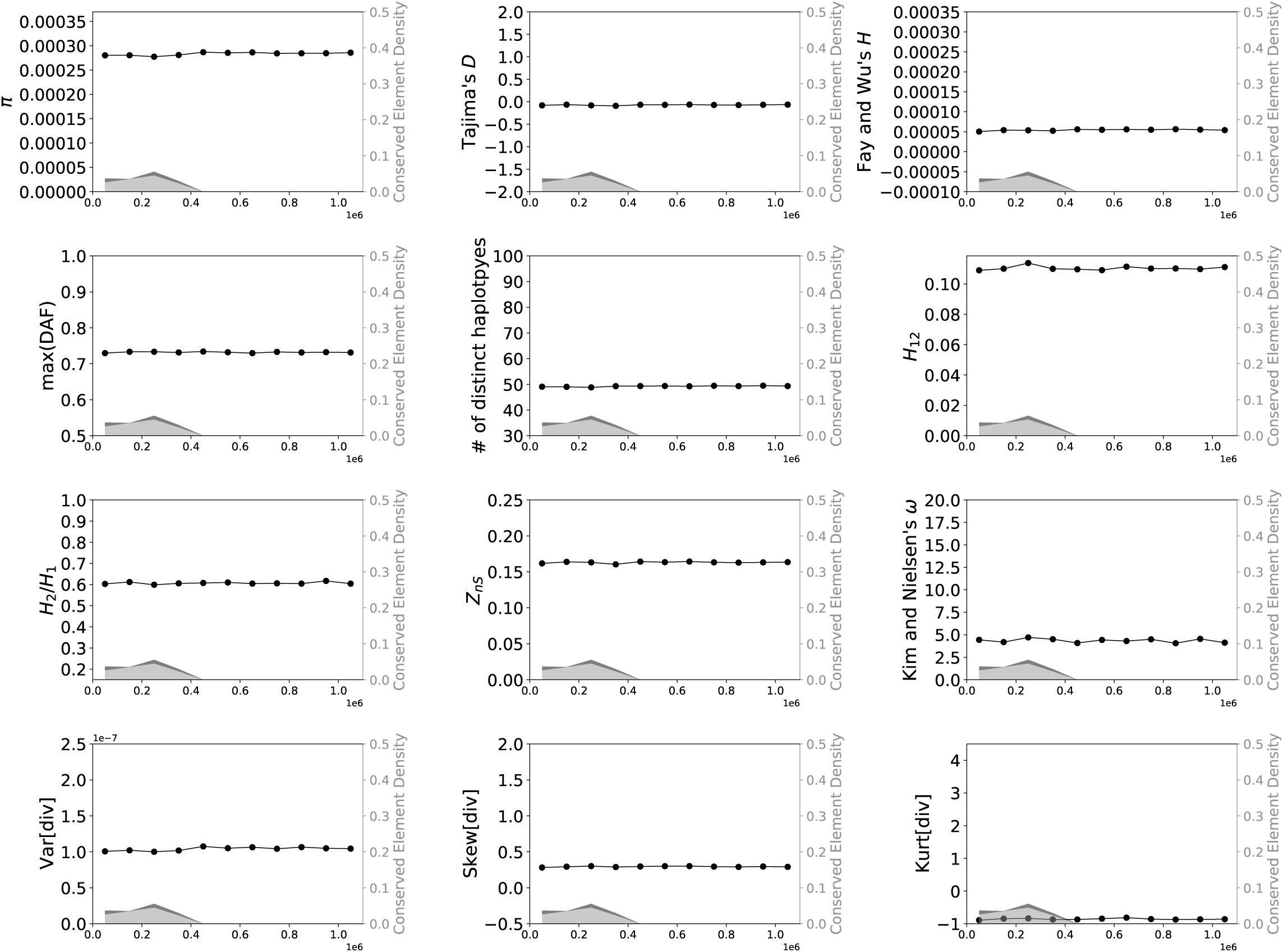
Values of 15 statistics calculated from a simulated data set designed to match the locations of conserved sites from chr11:50000001–51100000 in the human genome (hg19 assembly version), simulated under European population size history (Tennessen *et al*. 2012). See legend of Supplementary Figure 1 for more detail.

**Supplementary Figure 20:**
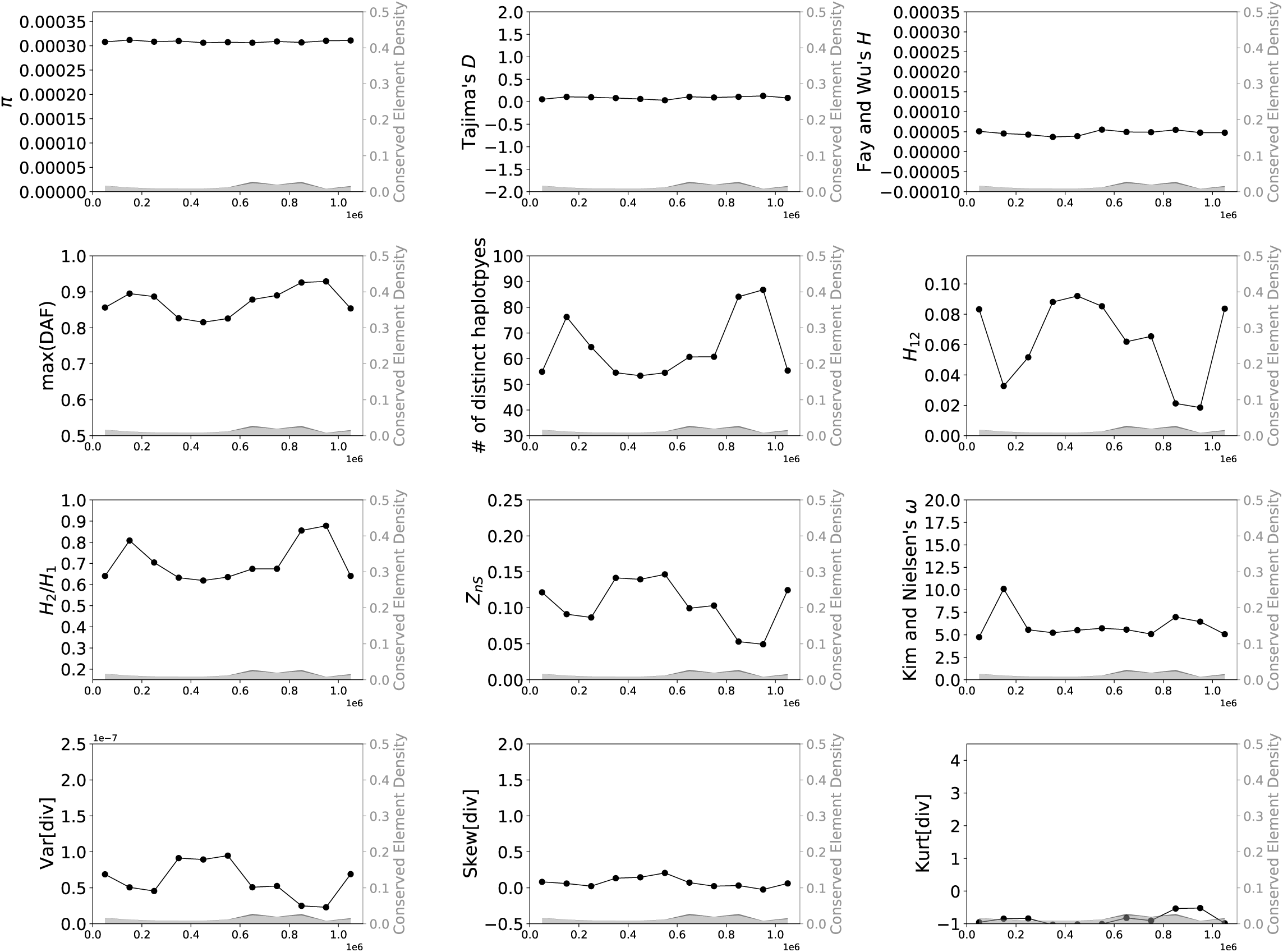
Values of 15 statistics calculated from a simulated data set designed to match the locations of conserved sites from chr11:23900001–25000000 in the human genome (hg19 assembly version), simulated under European population size history (Tennessen *et al*. 2012). See legend of Supplementary Figure 1 for more detail.

**Supplementary Figure 21:**
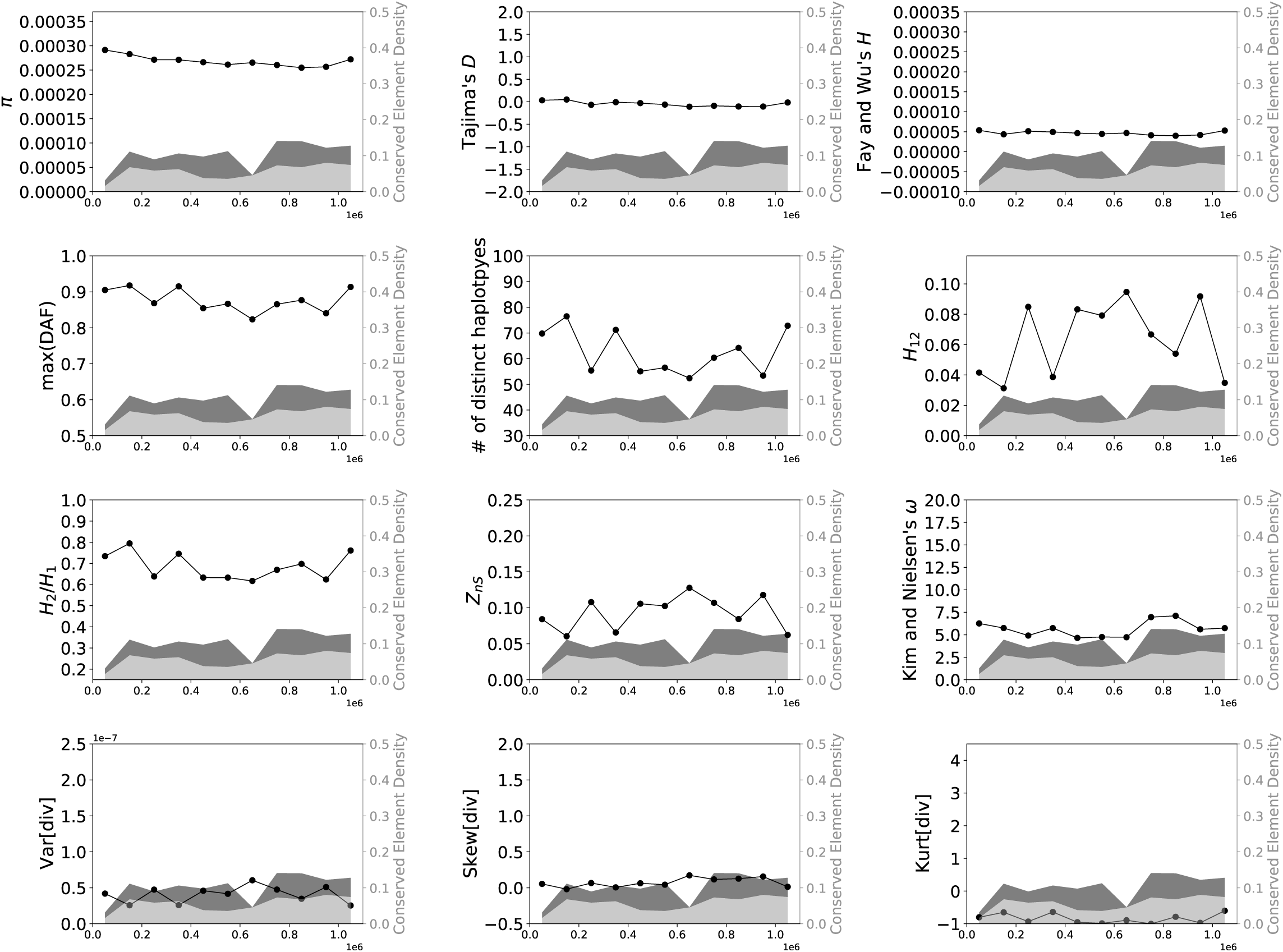
Values of 15 statistics calculated from a simulated data set designed to match the locations of conserved sites from chr10:49800001–50900000 in the human genome (hg19 assembly version), simulated under European population size history (Tennessen *et al*. 2012). See legend of Supplementary Figure 1 for more detail.

**Supplementary Figure 22:**
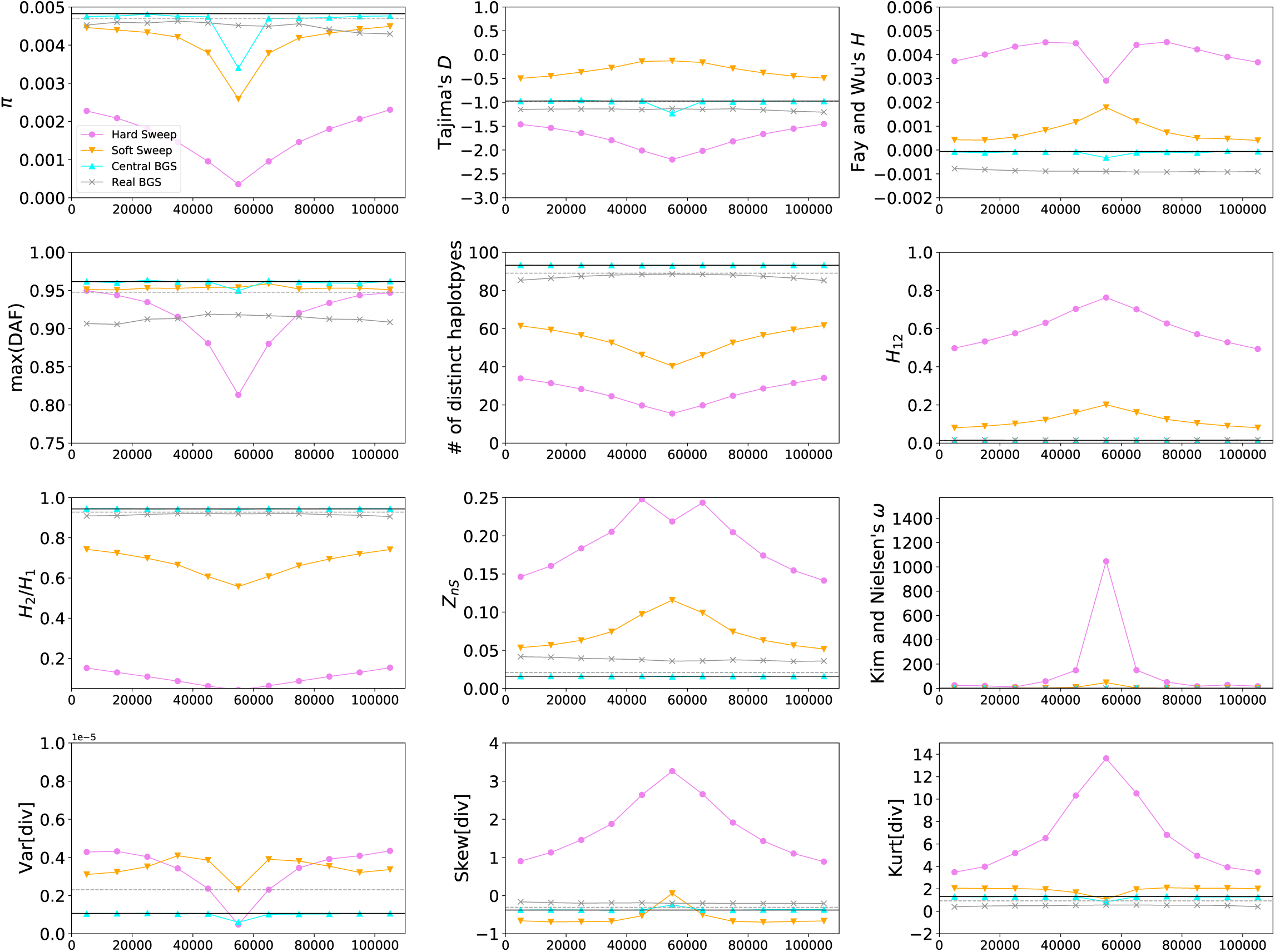
Values of 15 statistics calculated in each of the four data sets simulated under Sheehan and Song’s bottleneck model for African *Drosophila melanogaster* (Sheehan and Song 2016). Here, the mean value of each statistic across each simulated replicate is shown in the 11 adjacent 10 kb windows within the 110 kb chromosomal region. In each panel the neutral expectation of the statistic obtained from forward simulations are shown as a horizontal black line, while the neutral expectation from coalescent simulations is shown as a dashed gray line. The density of exonic and conserved non-coding DNA is shown in the shaded dark gray and light gray regions, respectively.

**Supplementary Figure 23:**
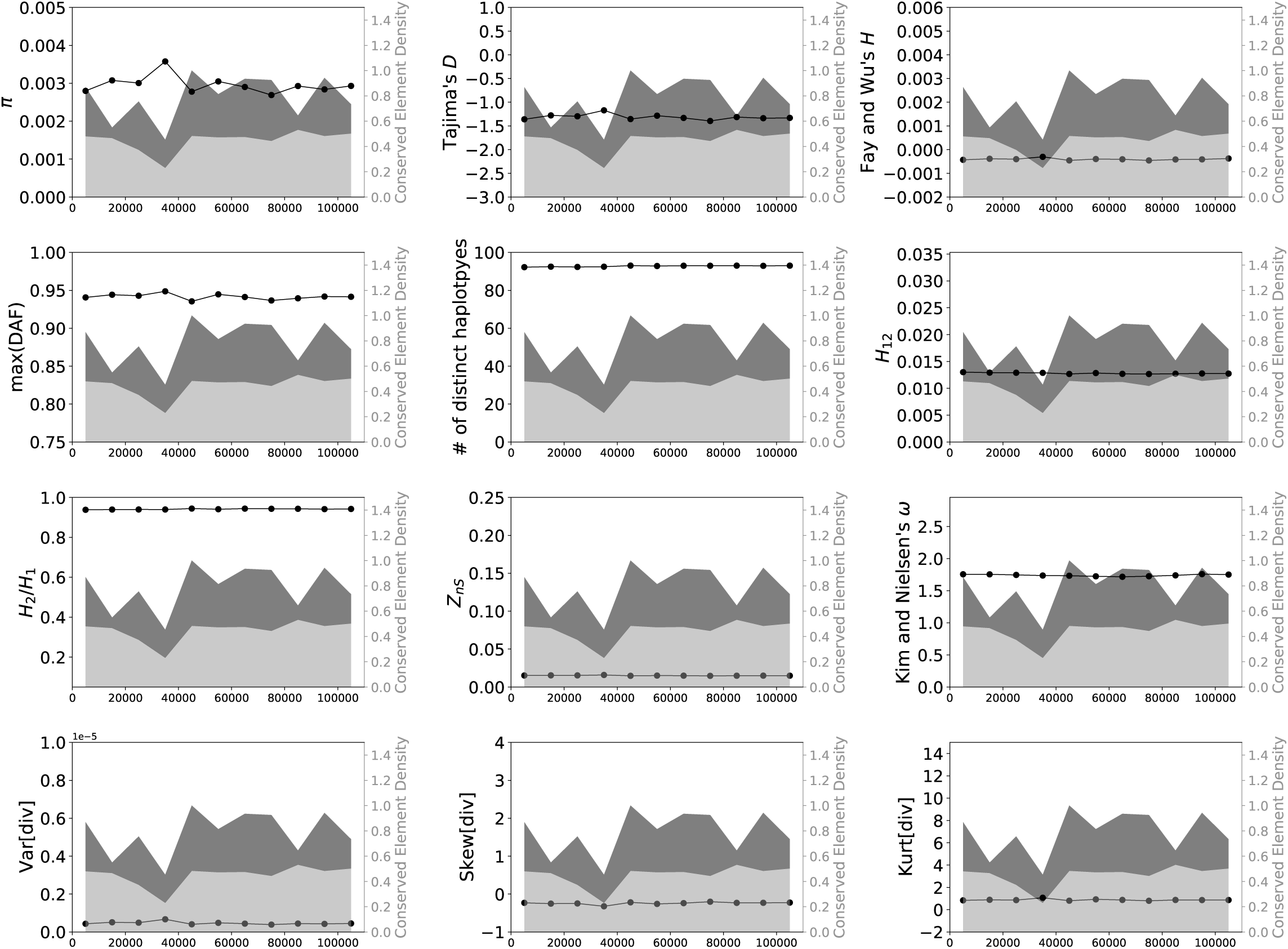
Values of 15 statistics calculated from a simulated data set designed to match the locations of conserved sites from chr3L:7460001–7570000 in the *Drosophila melanogaster* genome (assembly release 5), simulated under African population history (Sheehan and Song 2016). See legend of Supplementary Figure 1 for more detail.

**Supplementary Figure 24:**
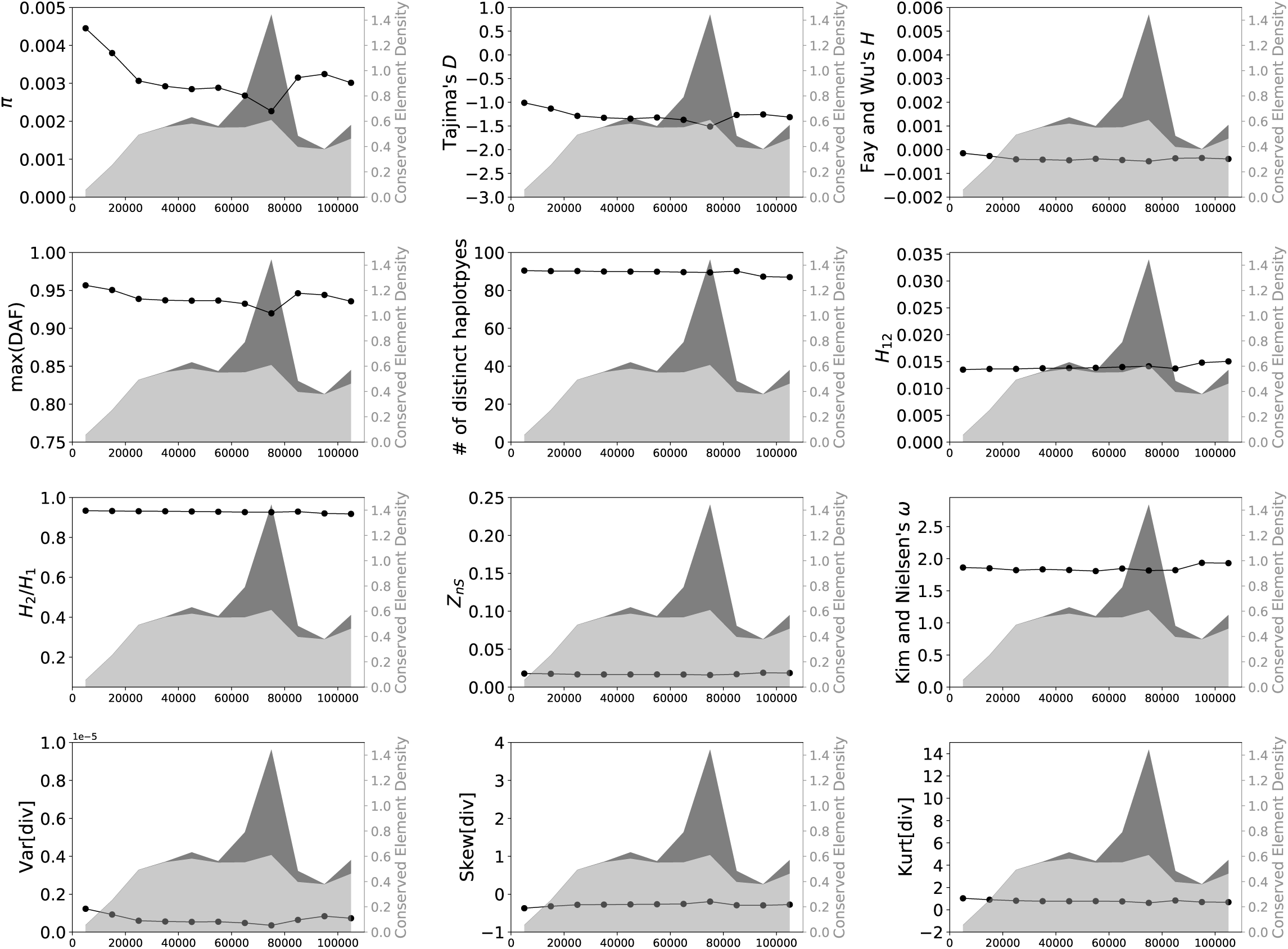
Values of 15 statistics calculated from a simulated data set designed to match the locations of conserved sites from chr3L:6410001–6520000 in the *Drosophila* genome (assembly release 5), simulated under African population history (Sheehan and Song 2016). See legend of Supplementary Figure 1 for more detail.

**Supplementary Figure 25:**
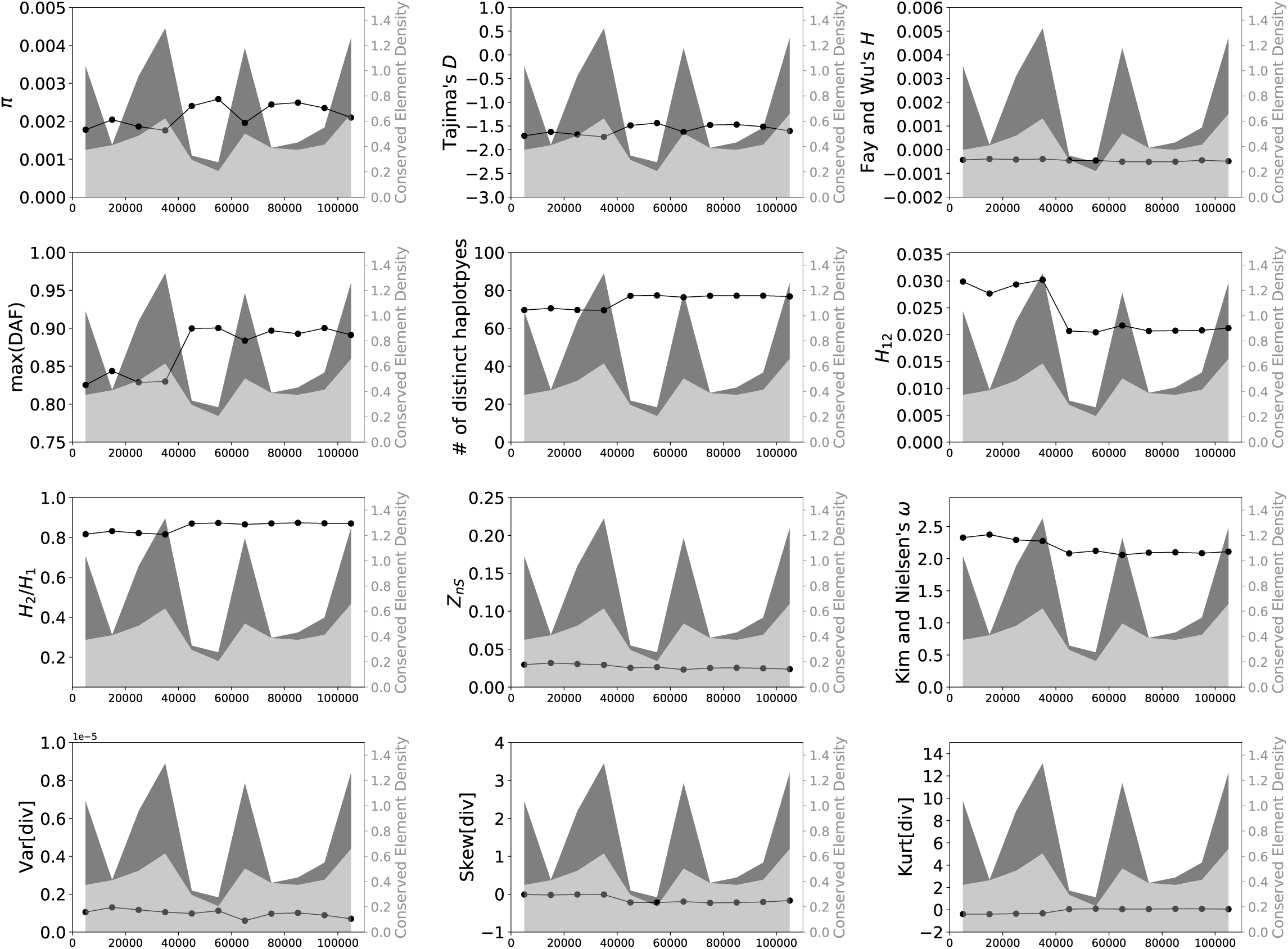
Values of 15 statistics calculated from a simulated data set designed to match the locations of conserved sites from chr3L:20960001–21070000 in the *Drosophila* genome (assembly release 5), simulated under African population history (Sheehan and Song 2016). See legend of Supplementary Figure 1 for more detail.

**Supplementary Figure 26:**
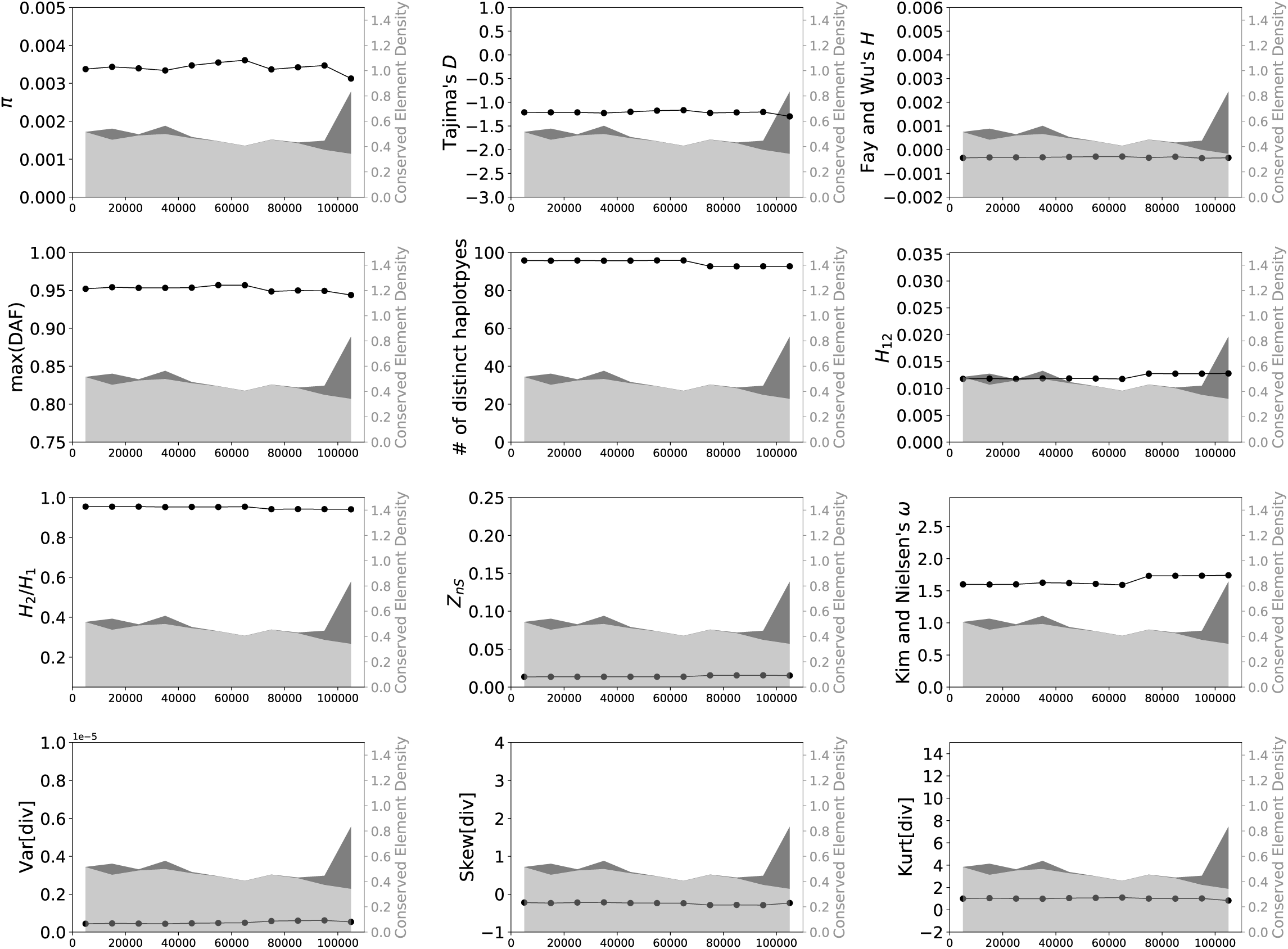
Values of 15 statistics calculated from a simulated data set designed to match the locations of conserved sites from chr3L:17130001–17240000 in the *Drosophila* genome (assembly release 5), simulated under African population history (Sheehan and Song 2016). See legend of Supplementary Figure 1 for more detail.

**Supplementary Figure 27:**
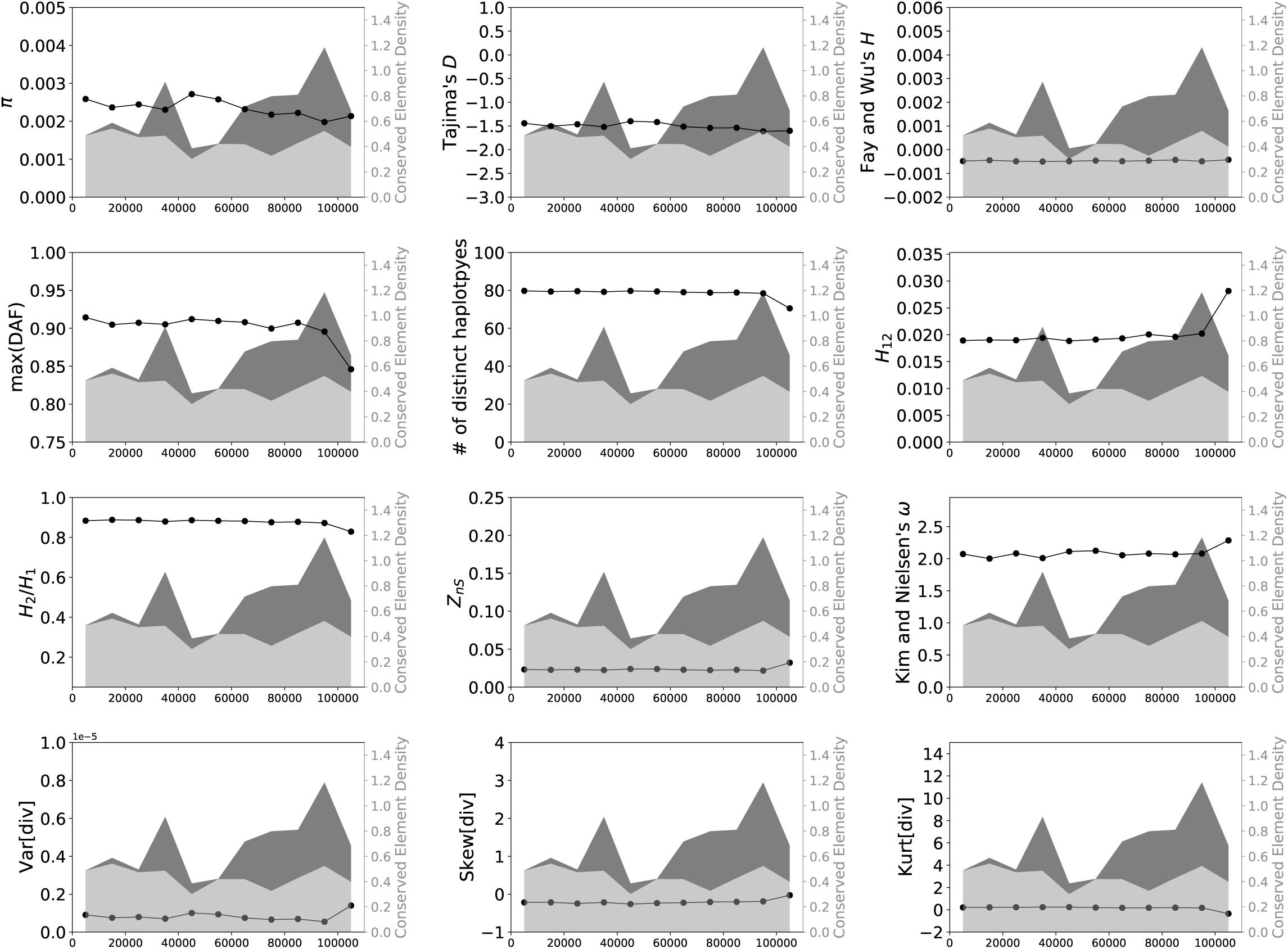
Values of 15 statistics calculated from a simulated data set designed to match the locations of conserved sites from chr3L:14900001–15010000 in the *Drosophila* genome (assembly release 5), simulated under African population history (Sheehan and Song 2016). See legend of Supplementary Figure 1 for more detail.

**Supplementary Figure 28:**
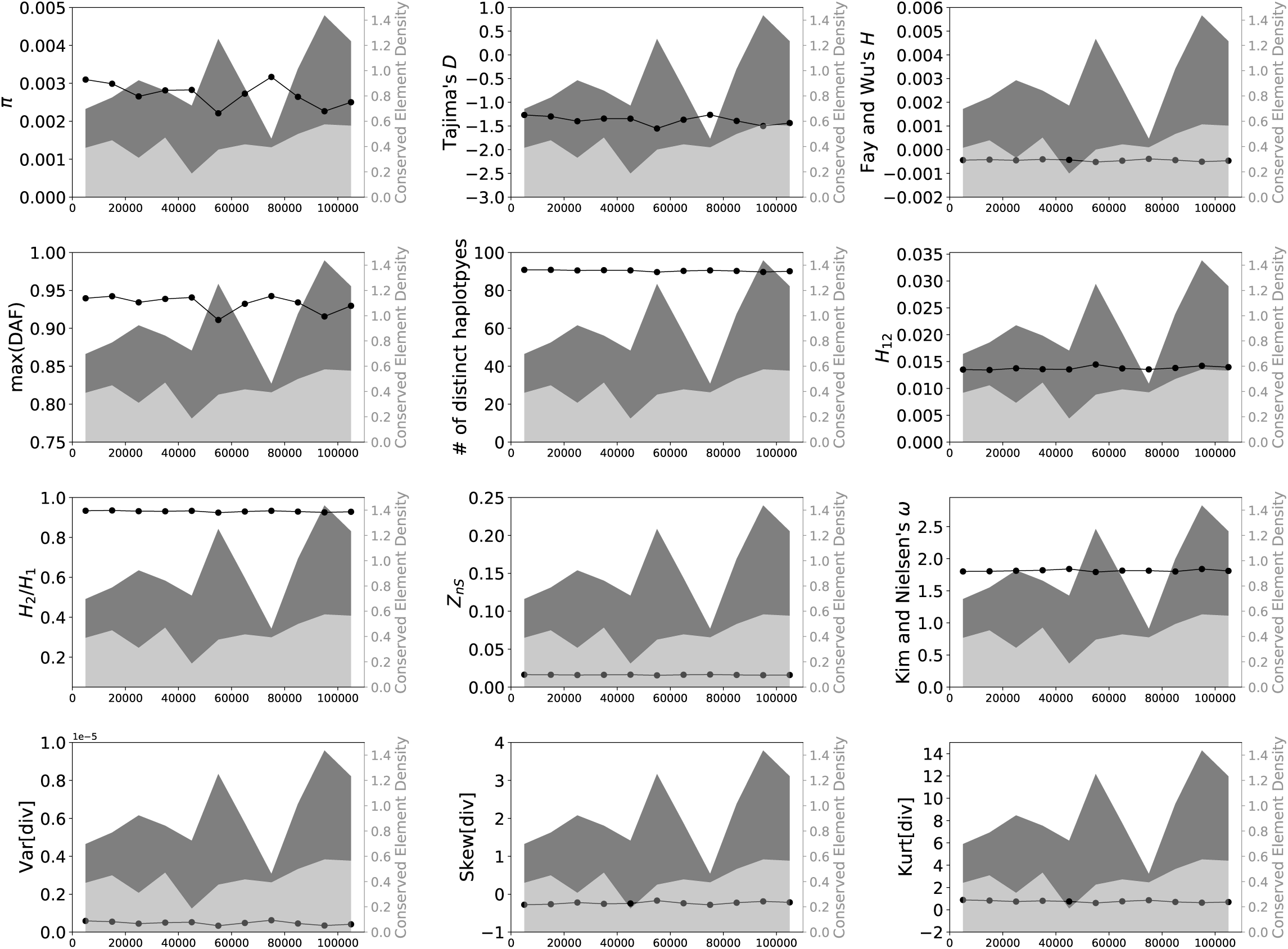
Values of 15 statistics calculated from a simulated data set designed to match the locations of conserved sites from chr2R:20360001–20470000 in the *Drosophila* genome (assembly release 5), simulated under African population history (Sheehan and Song 2016). See legend of Supplementary Figure 1 for more detail.

**Supplementary Figure 29:**
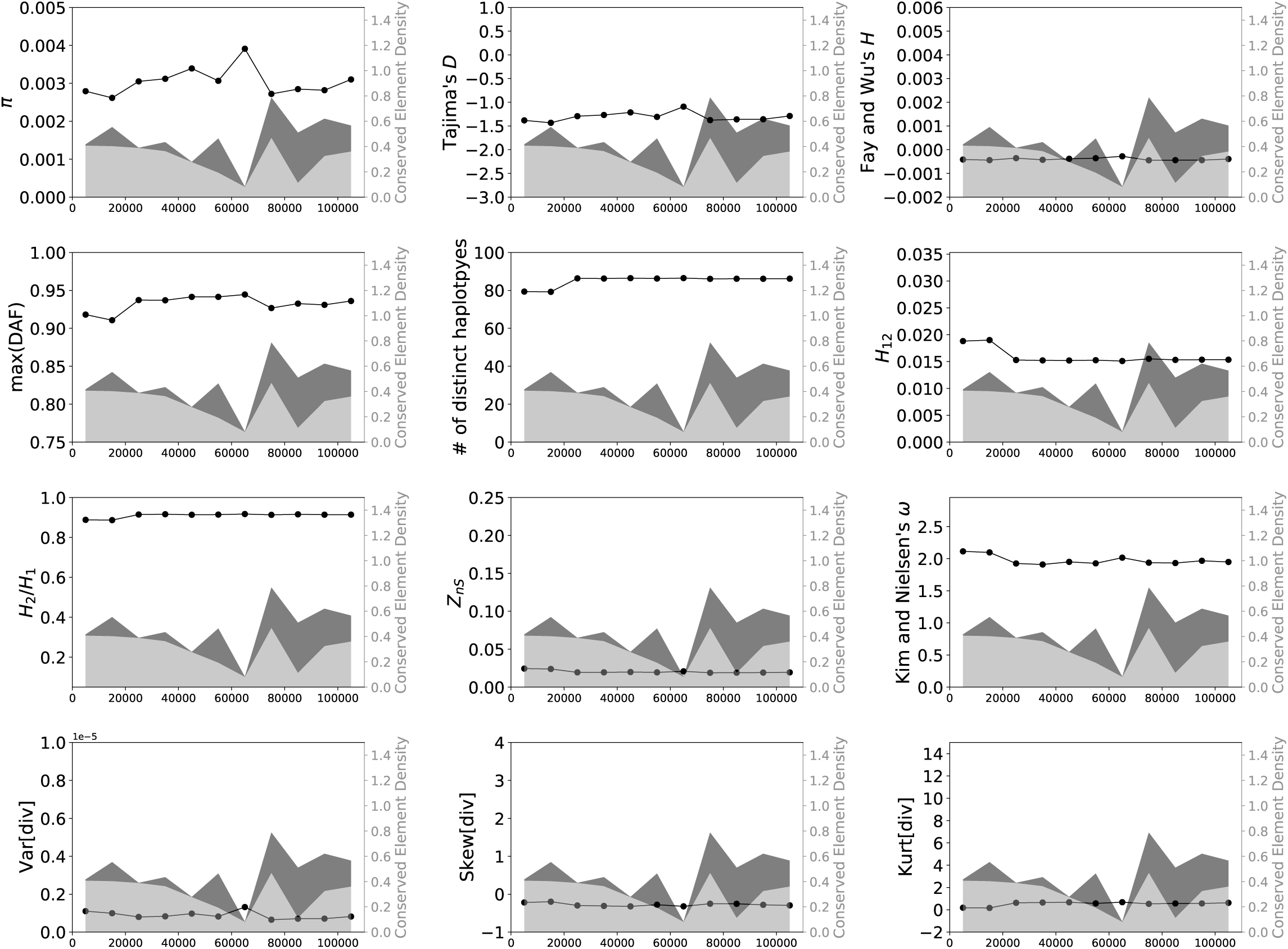
Values of 15 statistics calculated from a simulated data set designed to match the locations of conserved sites from chr3L:17980001–18090000 in the *Drosophila* genome (assembly release 5), simulated under African population history (Sheehan and Song 2016). See legend of Supplementary Figure 1 for more detail.

**Supplementary Figure 30:**
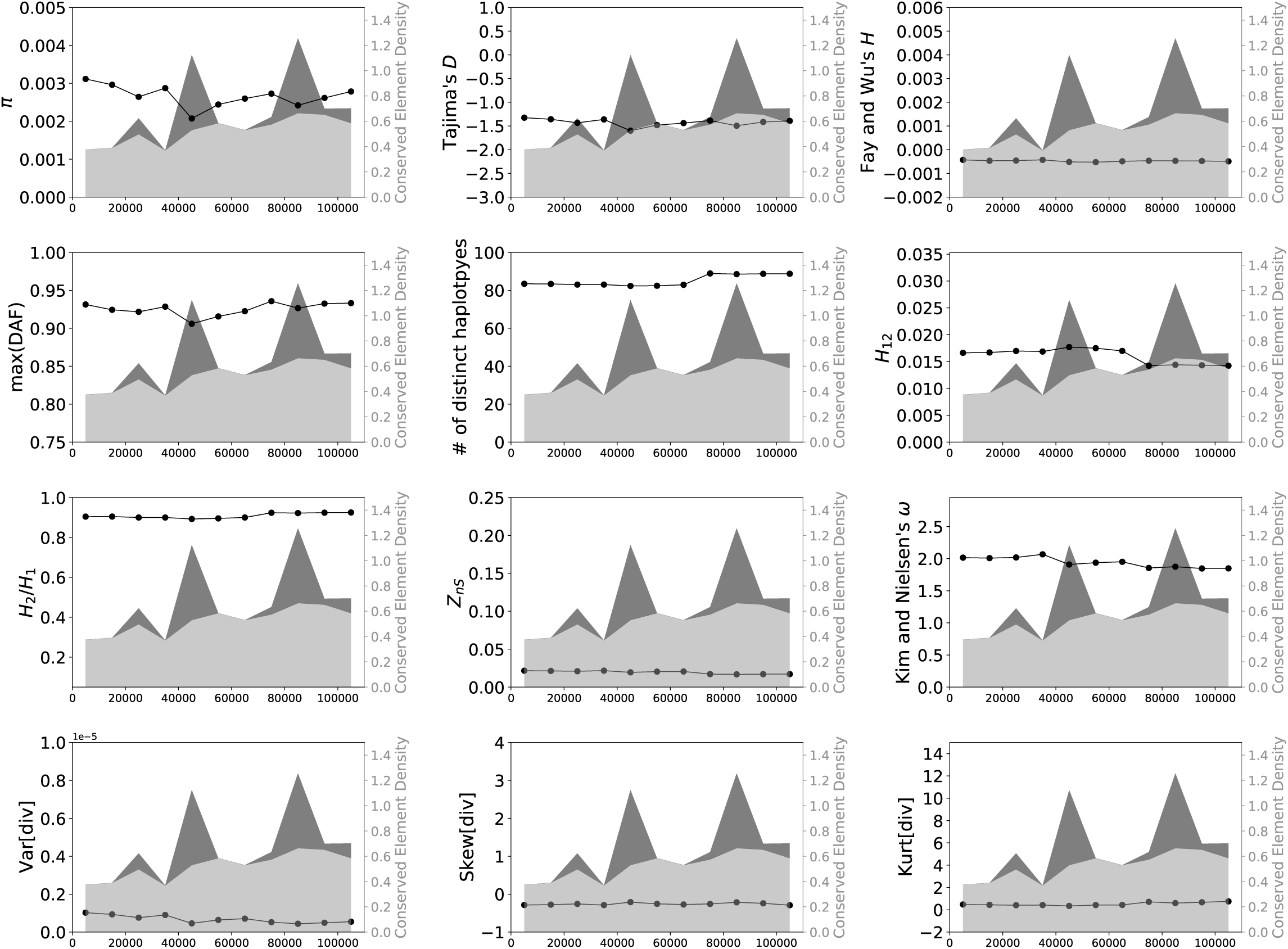
Values of 15 statistics calculated from a simulated data set designed to match the locations of conserved sites from chr2R:15630001–15740000 in the *Drosophila* genome (assembly release 5), simulated under African population history (Sheehan and Song 2016). See legend of Supplementary Figure 1 for more detail.

**Supplementary Figure 31:**
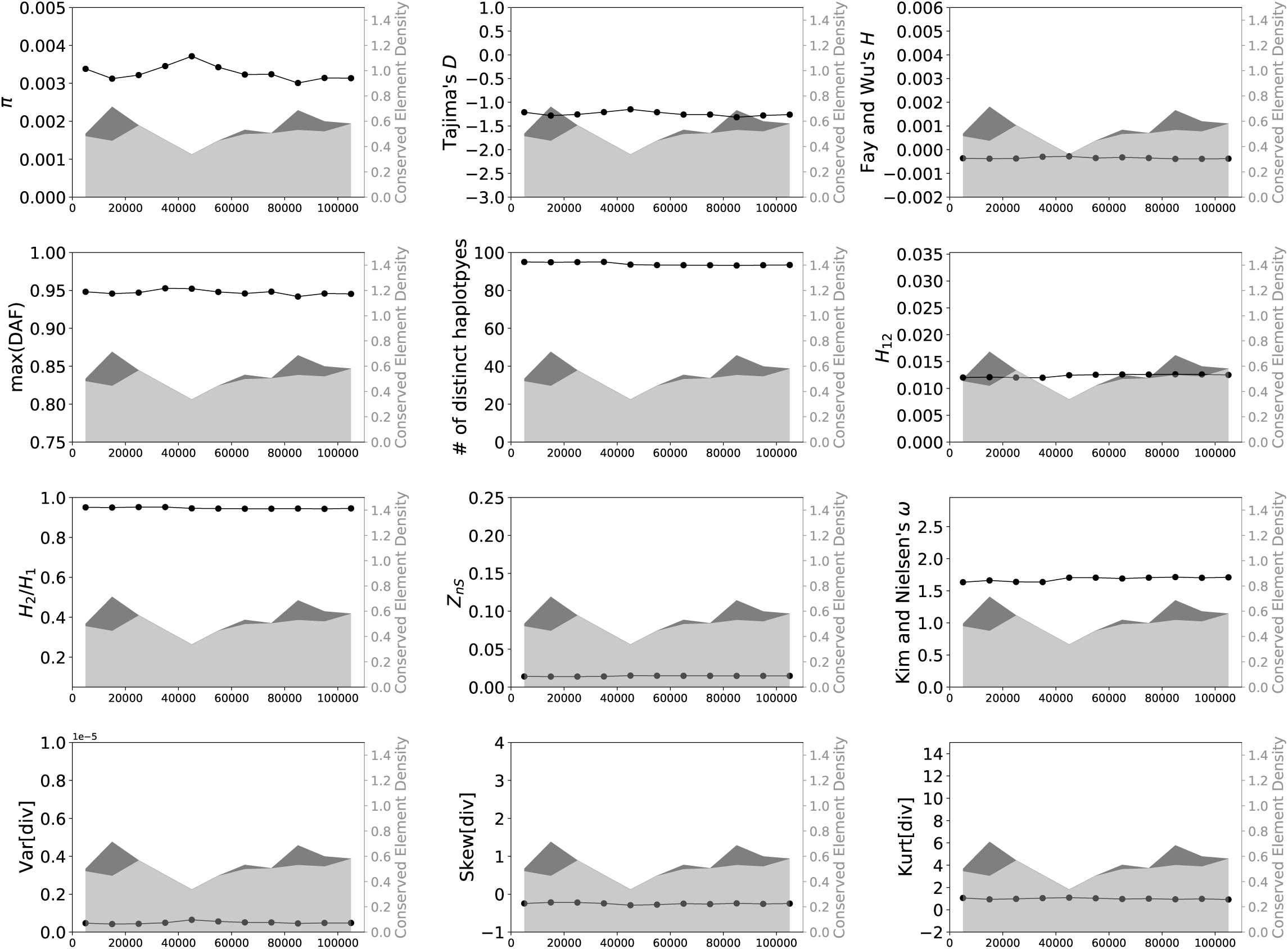
Values of 15 statistics calculated from a simulated data set designed to match the locations of conserved sites from chr3R:18660001–18770000 in the *Drosophila* genome (assembly release 5), simulated under African population history (Sheehan and Song 2016). See legend of Supplementary Figure 1 for more detail.

**Supplementary Figure 32:**
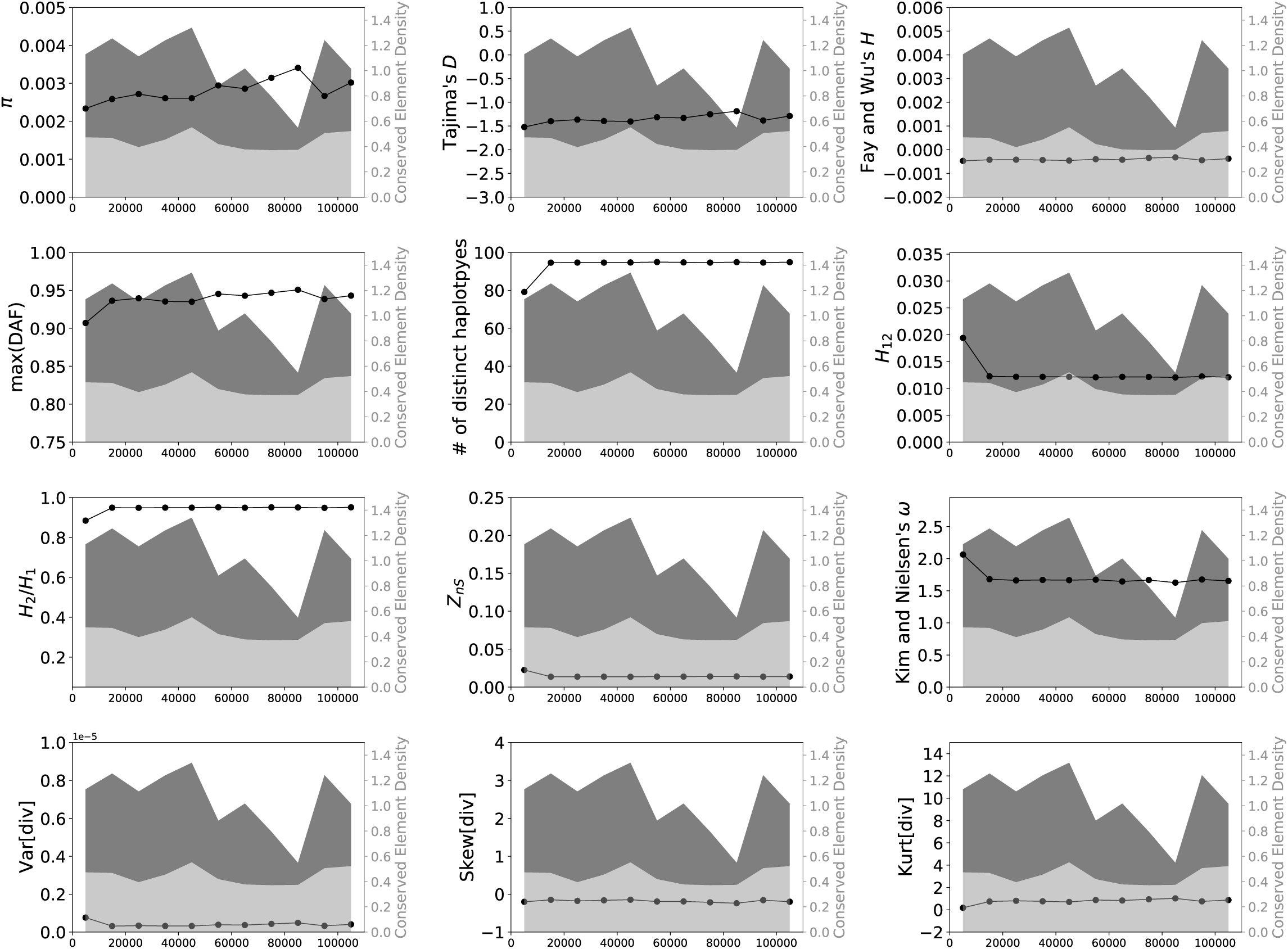
Values of 15 statistics calculated from a simulated data set designed to match the locations of conserved sites from chr2L:10390001–10500000 in the *Drosophila* genome (assembly release 5), simulated under African population history (Sheehan and Song 2016). See legend of Supplementary Figure 1 for more detail.

**Supplementary Figure 33:**
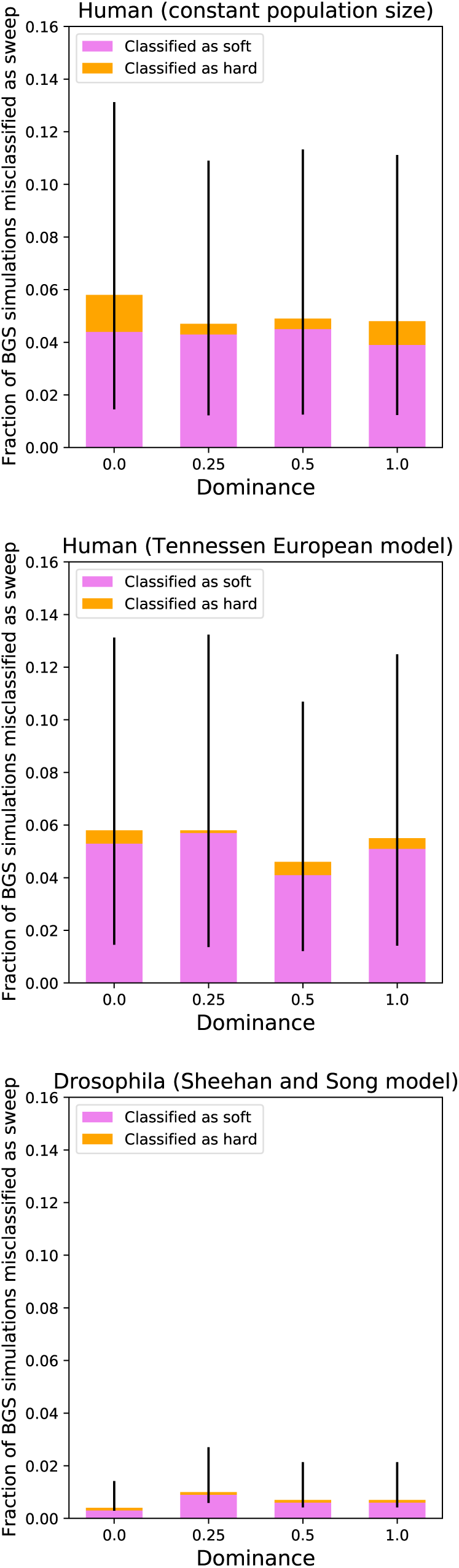
The fractions of simulations of our Real BGS model misclassified as hard and soft sweeps, shown for simulated data sets with different fixed dominance coefficients for deleterious mutations. The results for a dominance coefficient of 0.25 summarizes results from the same set of simulations reported elsewhere in the text, while the other simulations were generated by repeating the same procedure described in the Methods but altering the dominance coefficient accordingly. The error bars show the 95% binomial confidence intervals for our estimates of the total fraction of simulations misclassified as a sweep of either type.

**Supplementary Figure 34:**
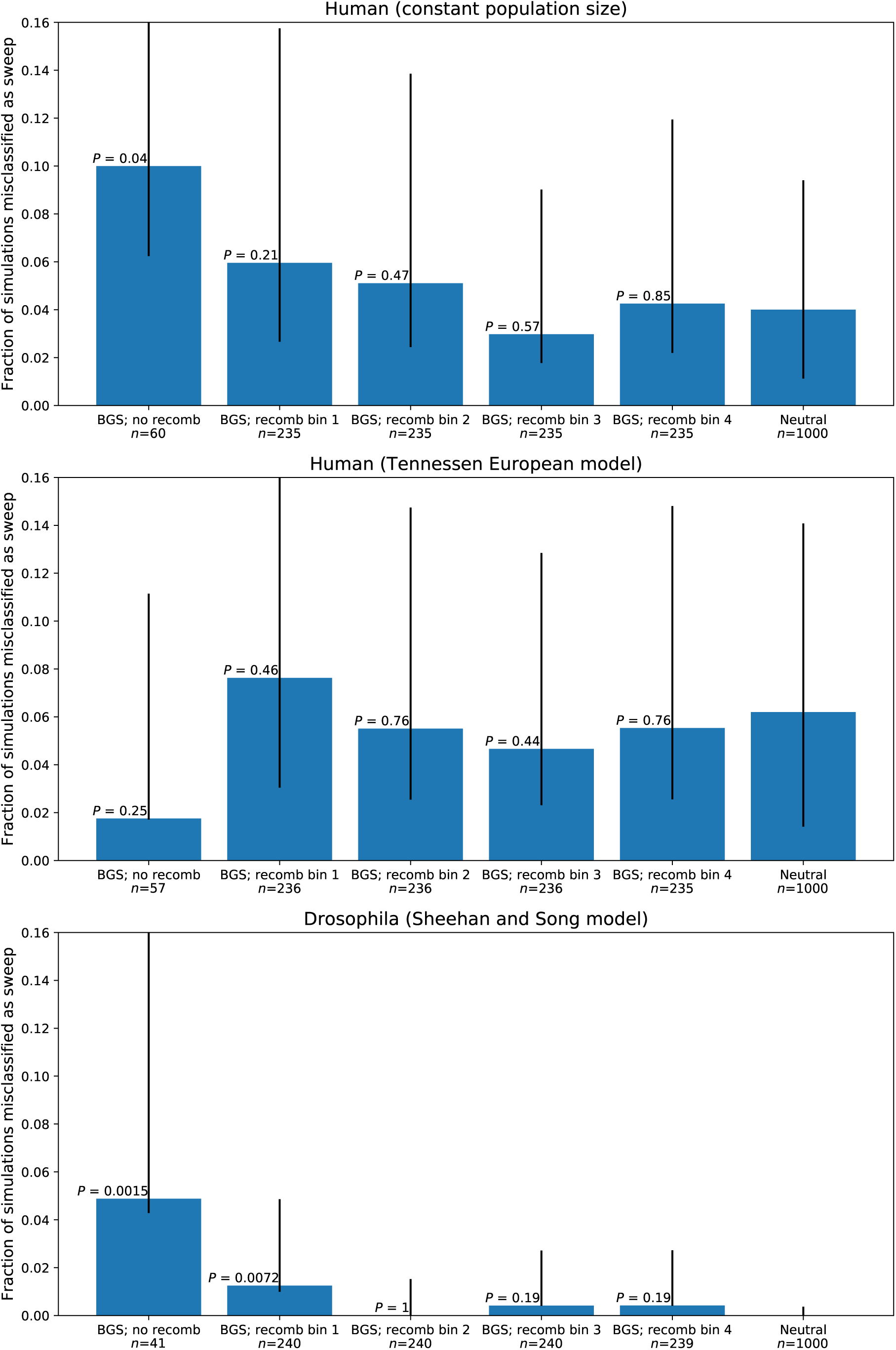
The fractions of simulations of our Real BGS–weak CNE model misclassified as a sweep of either type for each demographic model, shown after binning our data. The rightmost bar shows the fraction of misclassified neutral simulations for comparison, and all *P*-values show significance of the comparison with neutrality. The error bars show the 95% binomial confidence intervals.

